# Novel psilocin prodrugs with altered pharmacological properties as candidate therapies for treatment-resistant anxiety disorders

**DOI:** 10.1101/2023.05.16.540994

**Authors:** Sheetal A. Raithatha, Jillian M. Hagel, Kaveh Matinkhoo, Lisa Yu, David Press, Sarah G. Cook, Govinda Sharma, Dhananjaya D., Glynnis Jensen, Jessica B. Lee, Charlie Cai, Jonathan Gallant, Jaideep S. Bains, Joseph E. Tucker, Peter J. Facchini

## Abstract

The psychedelic compound psilocybin has shown therapeutic benefit in the treatment of numerous psychiatric diseases. A recent randomized clinical trial conducted at Johns Hopkins Bayview Medical Center demonstrated the efficacy of psilocybin-assisted therapy in the treatment of Major Depressive Disorder (MDD). Similarly, a phase IIb study evaluating psilocybin-assisted therapy for treatment-resistant depression (TRD) presented statistically meaningful and long-term reduction in depressive symptoms. Also, many studies have reported the successful treatment of severe anxiety after a single oral dose of psilocybin, especially in patients struggling with cancer-related distress (CRD). Despite these compelling clinical results, concerns regarding the duration of the psychedelic experience produced by psilocybin pose a significant barrier to its widespread therapeutic application. Psilocybin, derived from magic mushrooms is the naturally occurring prodrug of the neuroactive compound psilocin. When orally administered, exposure to the acidic gastrointestinal (GI) environment together with enzymatic processing by intestinal and hepatic alkaline phosphatase lead to the dephosphorylation of psilocybin producing elevated levels of systemic psilocin. These plasma levels are detectable up to 24 h and produce a psychoactive episode lasting as long as 6 h post-ingestion. In order to positively modify the kinetics of the acute psychedelic response, we have engineered a library of novel prodrug derivatives (NPDs) of psilocin, introducing a diversity of alternative metabolically cleavable moieties modified at the 4-carbon position of the core indole ring. This library consists of twenty-eight unique compounds represented by nine distinct prodrug classes. Each molecule was screened *in vitro* for metabolic stability using isolated human serum, and human cellular fractions derived from liver and intestinal tissues. This screen revealed fifteen prodrugs that produced measurable levels of psilocin *in vitro*, with ester and thiocarbonate-based prodrug derivatives significantly represented. These fifteen NPDs were further evaluated for pharmacokinetic (PK) profiles in mice, assessing plasma levels of both residual prodrug and resultant psilocin. PK results confirmed the efficiency of ester and thiocarbonate-based prodrug metabolism upon oral and intravenous administration, achieving levels reduced, albeit comparable to levels of psilocybin-derived psilocin. Of note, almost all NPDs tested maintained reduced overall exposure of psilocin relative to psilocybin, with no measurable levels detected at 24 h post-dose. Finally, all NPDs were screened for CNS bioavailability in healthy mice using the Head Twitch Response (HTR), a behavioural biomarker of 5-HT_2A_ receptor stimulation and an established proxy for psychoactive potential. Interestingly, five NPDs produced peak HTR that approached or exceeded levels induced by an equivalent dose of psilocybin. Among these bioactive prodrugs, an ester-based and thiocarbonate-based molecule produced long-term anxiolytic benefit in chronically stressed mice evaluated in the marble burying psychiatric model. Overall, this screening campaign identified novel candidate prodrugs of psilocin with altered metabolic profiles and reduced pharmacological exposure, potentially attenuating the duration of the psychedelic response. These molecules still maintained the long-term psychiatric and physiological benefits characteristic of psilocybin therapy. Additionally, these modified parameters also offer the opportunity for altered routes of administration bypassing conventional oral dosing.

## Introduction

Treatment-resistant depression and anxiety disorders remain significant unmet medical needs, affecting hundreds of millions of people worldwide (World Health Organization [WHO], 2023). This reality warrants the continuous search for rapid-acting anti-depressant and anxiolytic medicines. For thousands of years, psychedelic substances have been used by human cultures for holistic healing and spiritual purposes. In recent years, numerous drug development efforts have focused on the neuroactive, serotonergic functions of these natural hallucinogens for their therapeutic potential (1). Amongst these agents, there has been an increased interest in the use of psilocybin for the treatment of numerous psychiatric disorders (2). Found in various species of fungi (3), psilocybin is a naturally occurring prodrug of the psychedelic compound psilocin (4–6). Recent clinical research has reported compelling results demonstrating psilocybin’s therapeutic efficacy in addressing treatment resistant depression (TRD) (7–10). Significant and meaningful improvement in depression scores were presented, with long-term benefits lasting weeks, and in some individuals up to six months post-treatment. Similarly, a randomized clinical trial enrolling patients with moderate to severe major depressive disorder (MDD) demonstrated substantial and durable anti-depressant effects from psilocybin-assisted therapy, lasting up to 12 months (11,12). Additionally, in a randomized, double-blind, cross-over trial, a single dose of psilocybin was shown to produce substantial and enduring improvements in depressed mood, severe anxiety, and quality of life in patients struggling with life-threatening cancer diagnoses (13). These rapid and persistent anti-depressive and anxiolytic effects produced by psilocybin are characteristically accompanied by a potent psychedelic episode lasting up to 6 h from the time of dosing (4,7,14–16). The long duration of this altered state is attributed to the PK profile of psilocybin (6,17). Upon ingestion, exposure to acidic conditions within the GI tract, along with intestinal and hepatic alkaline phosphatase lead to the dephosphorylation of psilocybin, producing elevated levels of systemic psilocin. Participants consuming 25 mg of psilocybin experience significant systemic exposure to psilocin, with plasma levels peaking at 2 h, and enduring for up to 24 h post-administration.

Concerns around the duration of the psychedelic experience, which inevitably requires hours-long clinical supervision, significantly limits the widespread therapeutic application of psilocybin. Of note, comprehensive studies to determine the low threshold dose of psilocybin required to induce therapeutic benefit have not been conducted. However, subjective ratings for the acute drug effects reported at the conclusion of psilocybin-induced psychedelic sessions conducted at various dose levels have revealed interesting trends. These reports suggest that broad experiences, ranging from acute mystical-based effects to challenging experiences of paranoia, grief and fear, prove to produce an enduring sense of well-being and positive emotional benefits (reviewed in 18). To this end, it would stand to reason that psilocybin-based treatment maintains a wide therapeutic index with respect to effective dose range. Indeed, no significant correlation was observed between subjective drug effects and body weight (broadly ranging from 49 kg to 113 kg). This supports the fixed dose strategy used by current clinical trials (7,9–12,18). Owing to the somewhat dose-independent responses associated with the therapeutic effect of psilocybin, an alternative approach to reduce the extent and intensity of the psychedelic experience may be to modify the prodrug pharmacokinetics of systemic psilocin exposure. It can be argued that establishing a rapid onset and shorter duration of the acute psychoactive effects, while still retaining the long-term benefits reported for psilocybin-assisted treatment, will accommodate a more amendable psychotherapeutic process. To investigate this possibility, we engineered a library of novel prodrug derivatives (NPDs) of psilocin, incorporating a diversity of established metabolically amenable moieties substituted at the 4-hydroxyl group attached to the core indole ring. This library consists of twenty-eight unique compounds represented by nine distinct prodrug classes. This level of prodrug diversity encompasses derivatives with diverse physiochemical and biopharmaceutical properties, producing variable pharmacokinetic profiles, likely resulting in altered absorption and metabolic parameters. To investigate this diversity, we first screened the full collection of NPDs for metabolic stability using human intestine, liver and serum extracts. Those NPDs that produced measurable levels of psilocin *in vitro* were evaluated *in vivo* for systemic pharmacokinetic levels of psilocin, and resultant pharmacodynamic neuroactivity against the 5-HT_2A_ target receptor in orally treated healthy mice. Outcomes of these screens revealed five unique prodrug derivatives that produced notable acute levels of plasma psilocin, and reduced overall systemic exposure, as well as acute neuroactive responses approaching or exceeding levels induced by an equivalent dose of psilocybin. Of these active prodrugs, two distinct molecules produced long-term anxiolytic benefit in chronically stressed mice evaluated in the marble burying psychiatric model. Overall, this screening campaign identified novel candidate prodrugs of psilocin with altered metabolic profiles and reduced pharmacological exposure. These molecules still maintained the long-term behavioral benefits in mice characteristic of psilocybin therapy.

## Results

### Chemical synthesis

In line with our objective of modulating systemic psilocin exposure and pharmacokinetic properties of a psilocin prodrug, we embarked on the design and synthesis of unique prodrugs of this psychoactive molecule. An aromatic alcohol, psilocin bears a hydroxyl group at the 4-position of its indole core. The hydroxyl group of alcohol-based drugs have been extensively utilized as a handle for further derivatization, particularly to produce prodrugs (19). Taking advantage of the anticipated wide range of *in vivo* stability of linkers that can release an alcohol-based drug, several types of linkers were considered in designing our pool of NPDs, including esters, carbonates, phosphates, among others, which were all attached to the 4-hydroxy group of psilocin (Figure 1). Additionally, within each distinct group of prodrug linkers, various electronic properties and hydrophilicities were introduced to further modify physical and chemical features of the final prodrugs, namely their solubility and chemical stability.

**Figure 1.**
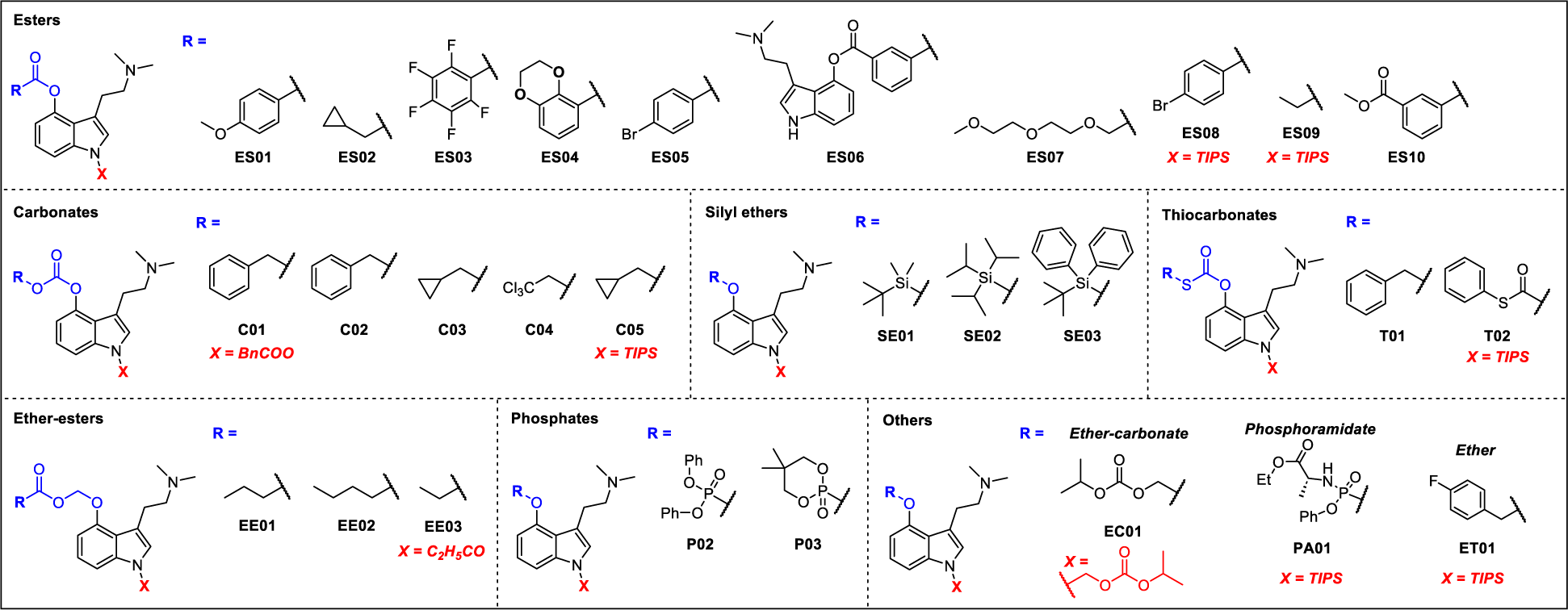
The chemical structures of the synthetic novel prodrug derivatives (NPDs) of psilocin. Unless shown otherwise, X = H.

To begin, psilocin was synthesized following the procedures found in the existing literature (20) with slight modifications that allowed us to utilize O-benzyl-protected 4-hydroxy-indole as the building block (Figure S1A). Following the production of psilocin, different prodrug linkers were installed simply *via* reacting the corresponding activated linker precursors with the 4-hydroxy group of psilocin (Figure S1B). This strategy was successful in generating the majority of our designed prodrugs; however, in some cases, the indole-NH interfered with modifications on the 4-OH and orthogonal derivatization was not feasible. To circumvent this issue, a triisopropylsilyl (TIPS) protection step was introduced to the synthesis to mask the indole-NH that was revealed following the installation of the corresponding linker (Figure S1C) (21). Subsequent to the 4-OH derivatization, in most cases, the *N*-TIPS protected moiety was also tested as an additional unique prodrug. Moreover, attempts to synthesize some of the designed prodrugs from psilocin led to the adventitious formation of NH-derivatized species which were tested separately. Following these two general synthetic routes, a total of 28 psilocin prodrugs within 9 distinct structural groups were synthesized (Figure 1).

### *In vitro* prodrug metabolism

Orally administered psilocybin is subject to significant first-pass metabolism, involving dephosphorylation at the 4-hydroxyl position producing the bioactive form psilocin (6,4,17,22–27). This dephosphorylation occurs in various compartments within the body. Chemical hydrolysis of the phosphate group readily occurs within the acidic environment of the GI tract (6,17,27). Enzymatic cleavage is mediated by alkaline phosphatase and non-specific esterases when psilocybin is absorbed through the intestinal lining and upon passage through the liver (4,17,25,26). Ultimately, generation of systemic levels of the more lipid soluble psilocin enhances bioavailability throughout the body, specifically making it amenable for transport across the blood-brain barrier and stimulation of receptors in the brain to induce its neuroactive effects (16,28,29).

Chemical modification of the cleavable moiety on the 4-hydroxyl position of these NPDs has the potential to modify the overall metabolic profile of these molecules. This altered metabolic potential will ultimately affect the pharmacokinetics (PK) of plasma psilocin and its overall bioavailability. Various assays are available to evaluate the potential metabolism of candidate drugs *in vitro* (30,31). A commonly established approach is the use of subcellular fractions derived from relevant organs involved in drug metabolism. Human microsomes, isolated from liver and intestinal tissues contain a wide array of relevant enzymes involved in biotransformation and drug metabolism (32–34). S9 subcellular fractions from either liver or intestinal tissues contain both microsomal and cytosolic components. Except for membrane-associated factors, S9 fractions essentially encompass the same metabolic enzyme profile as intact cells (35,36). To determine the *in vitro* metabolic parameters that might predict the pharmacokinetics and drug reactivity profile of each prodrug, all molecules, prepared at an assay concentration of 2.5 µM, were incubated independently in human serum, and in S9 or microsomal cell free fractions derived from both human liver and intestinal donor organ. Each molecule was also assessed for aqueous stability independent of enzyme activity in assay buffer alone. Biotransformation of the parent prodrug into the psychoactive metabolite psilocin was monitored for each molecule in 20-min intervals for a duration of 2 h by liquid chromatography mass spectrometry (LC-MS). As a relevant comparator, the natural prodrug psilocybin (PCB) was also assessed for its *in vitro* metabolic stability. Table 1 presents the *in vitro* half-life (T_1/2_) for each novel prodrug evaluated independently in each biological fraction. Table 2 lists the emergent concentration of psilocin (in µM) at the endpoint of each assay. Psilocybin, which is phosphorylated on the 4-hydroxyl of the indole core, proved to be considerably resilient to metabolism under these *in vitro* conditions, with only a marginal reactivity seen in intestinal-based fractions (Table 1 and 2, Figure S2). In contrast, many NPDs designed to be specifically targeted by esterase-directed cleavage were readily metabolized into psilocin in all biological fractions tested. Of these, the ester-based prodrug derivatives ES01 and ES02 produced notable enzyme-mediated cleavage profiles (Figures S3 and S4 respectively), with significant metabolism detected in liver- and serum-based fractions and appreciable activity seen in intestinal isolates. Notable metabolic conversion was also seen in one carbonate-based prodrug, C03 (Figure S5), and one thiocarbonate derivative, T01 (Figure S6). The chemical diversity in the R groups within this collection of compounds does not suggest any structural feature is fundamentally integral in promoting greater cleavage of the prodrug moiety to release psilocin; however, aromatic (e.g., R = benzyl, methoxybenzyl, halogenated benzyls) and aliphatic (e.g., R = ethyl, cyclopropyl) side chains are increasingly represented in metabolically labile compounds. Additionally, the removal of a zwitterionic state may make these compounds better esterase substrates. A compelling outcome in the above analysis is the notable metabolic activity detected in human serum samples against many of the NPDs (Table 1 and 2, Figure S3, S4, S5 and S6). This is in stark contrast to psilocybin, which is considerably stable in this biological fraction (Figure S2). These NPDs, proving amenable to serum-based metabolism, present an interesting therapeutic opportunity to circumvent first-pass processing by direct systemic administration.

**Table 1.** *In vitro* metabolic stability of psilocybin and twenty-eight novel prodrug derivatives (NPDs) upon exposure to human serum, and human S9 and microsomal fractions derived from donor liver and intestinal tissues. Enzymatic stability of each NPD is presented as *in vitro* half-life (T_1/2_). The chemical structure of each indicated NPD is presented in Figure 1. Each NPD was prepared at an assay concentration of 2.5 µM and incubated in each biological fraction independently for a total of 120 min. LC-MS-based quantification of remaining (non-metabolized) compound was performed at timepoints 0, 20, 40, 60, 80, 100 and 120 min. Plotted data was analyzed using exponential one-phase decay non-linear fit to determine T_1/2_. (aq) indicates assay conducted without biological fraction; “∞” indicates compound is stable over the 120-min incubation period.

| NPD* | *in vitro* Half-life ($T_{1/2}$) [min] | | | | | |
| --- | --- | --- | --- | --- | --- | --- |
|  | (aq) | Liver S9 Fraction | Liver Microsomes | Intestine S9 Fraction | Intestine Microsomes | Serum |
| PCB | ∞ | ∞ | ∞ | ∞ | 195.0 | ∞ |
| ES01 | ∞ | 21.0 | 17.0 | 132.0 | 315.0 | 1.0 |
| ES02 | ∞ | 3.0 | 1.0 | 104.0 | 62.0 | 1.0 |
| ES03 | 85.0 | 1.0 | 1.0 | 56.0 | 32.0 | 0.7 |
| ES04 | ∞ | 529.0 | 390.0 | ∞ | 113.0 | ∞ |
| ES05 | ∞ | 39.0 | 87.0 | 69.0 | 71.0 | 19.0 |
| ES06 | ∞ | ∞ | 241.0 | 445.0 | 621.0 | 101.0 |
| ES07 | 80.0 | 16.0 | 5.0 | 72.0 | 60.0 | 0.8 |
| ES08 | ∞ | 15.0 | 25.0 | 16.0 | 11.0 | ∞ |
| ES09 | ∞ | 3.0 | 2.0 | 2.0 | 2.0 | 49.0 |
| ES10 | ∞ | 33.0 | 22.0 | ∞ | ∞ | 33.0 |
| C01 | ∞ | 22.0 | 52.0 | 16.0 | 13.0 | 0.8 |
| C02 | ∞ | ∞ | 113.0 | 529.0 | 211.0 | ∞ |
| C03 | ∞ | 9.0 | 4.0 | 327.0 | 271.0 | 3.0 |
| C04 | ∞ | ∞ | 277.0 | 221.0 | 150.0 | 182.0 |
| C05 | 18.0 | 49.0 | 31.0 | 24.0 | 24.0 | 26.0 |
| T01 | ∞ | 5.0 | 6.0 | 50.0 | 71.0 | 14.0 |
| T02 | 6.0 | 7.0 | 9.0 | 24.0 | 13.0 | 32.0 |
| ET01 | ∞ | 60.0 | 27.0 | 64.0 | 50.0 | ∞ |
| EE01 | ∞ | 4.0 | 3.0 | 241.0 | 74.0 | 2.0 |
| EE02 | ∞ | 5.0 | 4.0 | 39.0 | 112.0 | 3.0 |
| EE03 | 145.0 | 16.0 | 13.0 | 52.0 | 22.0 | 0.8 |
| EC01 | ∞ | 293.0 | 104.0 | 266.0 | 894.0 | 3.0 |
| SE01 | ∞ | 191.0 | 347.0 | ∞ | ∞ | ∞ |
| SE02 | ∞ | 96.0 | 104.0 | 57.0 | 27.0 | ∞ |
| SE03 | ∞ | 391.0 | 229.0 | ∞ | ∞ | ∞ |
| P02 | ∞ | 96.0 | 43.0 | 52.0 | 28.0 | ∞ |
| P03 | ∞ | ∞ | ∞ | ∞ | ∞ | ∞ |
| PA01 | 10.0 | 26.0 | 22.0 | 39.0 | 29.0 | 145.0 |

\*ES = ester; C = carbonate; T = thiocarbonate; ET = ether; EE = ether-esters; EC = ether-carbonate; SE – silyl-ether; P = phosphate; PA = phosphoramidate; PCB = psilocybin

**Table 2.** *In vitro* metabolic stability of psilocybin and twenty-eight novel prodrug derivatives (NPDs) upon exposure to human serum, and human S9 and microsomal fractions derived from donor liver and intestinal tissues. Biotransformation into active metabolite, psilocin after 120-min incubation is presented in µM (endpoint concentration). The chemical structure of each indicated NPD is presented in Figure 1. Each NPD was prepared at an assay concentration of 2.5 µM and incubated in each biological fraction independently for a total of 120 min. LC-MS-based quantification of resultant psilocin metabolite was performed at timepoints 0, 20, 40, 60, 80, 100 and 120 min. (aq) indicates assay conducted without biological fraction.

| NPD* | Endpoint *in vitro* Psilocin Concentration [$\mu\text{M}$] | | | | |
| --- | --- | --- | --- | --- | --- |
|  | Liver S9 Fraction | Liver Microsomes | Intestine S9 Fraction | Intestine Microsomes | Serum |
| PCB | 0.000 | 0.000 | 0.621 | 0.172 | 0.000 |
| ES01 | 1.950 | 2.718 | 0.114 | 0.041 | 1.097 |
| ES02 | 1.343 | 1.890 | 1.384 | 0.927 | 0.778 |
| ES03 | 1.244 | 2.554 | 0.533 | 0.786 | 0.450 |
| ES04 | 0.145 | 0.329 | 0.000 | 0.000 | 0.000 |
| ES05 | 0.689 | 1.310 | 0.000 | 0.018 | 2.010 |
| ES06 | 0.000 | 0.000 | 0.000 | 0.000 | 0.000 |
| ES07 | 0.000 | 0.494 | 0.000 | 0.000 | 0.000 |
| ES08 | 0.656 | 1.659 | 0.195 | 0.887 | 0.000 |
| ES09 | 0.204 | 0.244 | 0.000 | 0.053 | 0.000 |
| ES10 | 0.451 | 0.919 | 0.166 | 0.241 | 3.148 |
| C01 | 0.000 | 0.000 | 0.000 | 0.000 | 0.000 |
| C02 | 0.084 | 0.114 | 0.044 | 0.038 | 0.000 |
| C03 | 1.177 | 1.929 | 0.224 | 0.157 | 0.361 |
| C04 | 0.133 | 0.057 | 0.000 | 0.000 | 0.000 |
| C05 | 0.072 | 0.186 | 0.018 | 0.096 | 0.000 |
| T01 | 1.233 | 2.105 | 0.000 | 0.156 | 0.385 |
| T02 | 0.133 | 0.206 | 0.000 | 0.000 | 0.000 |
| ET01 | 0.000 | 0.000 | 0.000 | 0.000 | 0.000 |
| EE01 | 0.012 | 0.073 | 0.018 | 0.043 | 0.000 |
| EE02 | 0.070 | 0.132 | 0.055 | 0.081 | 0.000 |
| EE03 | 0.000 | 0.001 | 0.000 | 0.000 | 0.000 |
| EC01 | 0.186 | 0.056 | 0.000 | 0.000 | 0.000 |
| SE01 | 0.000 | 0.012 | 0.000 | 0.000 | 0.000 |
| SE02 | 0.000 | 0.000 | 0.000 | 0.000 | 0.142 |
| SE03 | 0.000 | 0.000 | 0.000 | 0.000 | 0.000 |
| P02 | 0.073 | 0.000 | 0.000 | 0.076 | 0.069 |
| P03 | 0.011 | 0.000 | 0.000 | 0.000 | 0.000 |
| PA01 | 0.153 | 0.324 | 0.470 | 0.000 | 0.000 |

\*ES = ester; C = carbonate; T = thiocarbonate; ET = ether; EE = ether-esters; EC = ether-carbonate; SE – silyl-ether; P = phosphate; PA = phosphoramidate; PCB = psilocybin

Contrary to the intended design, a few NPDs, specifically C01 and EE03, which appeared to be efficiently metabolized in some or all fractions tested (Table 1), did not produce any detectable psilocin (Table 2). Interestingly, both these molecules are modified at the indole-NH, which may suggest inappropriate substrate recognition for cleavage of R groups at this site. As the bioanalytical method was developed to only evaluate loss of the parent NPD and emergence of the intended psilocin analyte, intermediate product(s) of these compounds remain unanalyzed. Overall, most NPDs appeared generally stable in aqueous conditions except for C05, T02, and PA01, which show notable T_1/2_ values under these control conditions (Table 1). Interestingly, each of these novel prodrugs maintain a triisopropylsilyl ether (TIPS) protecting group on the indole-NH. Of note, the results of C05 and T02 are in stark contrast to their respective synthetic counterparts, C03 (carbonate-linked cyclopropyl) and T01 (thiocarbonate-linked benzyl), which lack the TIPS moiety on the indole-NH. These NPDs are stable and are readily metabolized into psilocin in liver and serum-based fractions. It is important to note that addition of the TIPS group does not universally generate instability, since compounds ES08 and ES09 both remain stable under aqueous conditions and are metabolized into psilocin in various fractions. However, some distinctions in serum metabolism are observed between ES08 and its synthetic counterpart, ES05, suggesting additional enzymatic steps are required to cleave the TIPS moiety that may not be present in isolated serum fractions.

### Murine Plasma Pharmacokinetics

Metabolic assessment *in vitro* using subcellular fractions can provide an estimation of first-pass metabolism; however, these fractions do not represent the full enzyme complement of their representative tissues. Furthermore, available methods lack appropriate standardization making *in vitro* results only indicative of the true metabolic profile (30). To further evaluate the metabolic characteristics of these NPDs, a selection of molecules observed to produce psilocin *in vitro*, and representing various cleavable motifs described above were chosen for plasma pharmacokinetic (PK) analysis in healthy mice (Figure S7). Of note, a carbonate-based NPD C01, which demonstrated appreciable instability in all biological fractions (Table 1) yet did not produce psilocin (Table 2), was also elected as a non-conforming test article for analysis. Each compound was administered intravenously (i.v.) at 1 mg/kg, and via oral gavage (per os, p.o.) at 1, 3 and 10 mg/kg. Serial blood samples were evaluated for concentrations of each parent NPD and psilocin over a period of 24 h. As a comparator for PK profiling, psilocybin was also analyzed under the same parameters. When dosed at 1 mg/kg i.v., most NPDs produced notable levels of plasma psilocin reaching maximum concentrations (C_max_) of at least 25% of those detected with psilocybin at the same i.v. dose (Figure 2, Table S1). Indeed, six molecules, ES01, ES02, C02, C03, T01, and EE02 produced psilocin levels approaching or exceeding that derived from psilocybin. Of these, the thiocarbonate-based prodrug, T01 produced a C_max_ three to five times greater than psilocybin and maintained an overall systemic exposure of psilocin (AUC_(0-tlast)_) equivalent to the same dose of psilocybin (Table S1). In contrast, psilocin produced from orally administered psilocybin at 1 mg/kg exceeded all NPDs tested by at least 80% (Figure 2, Table S1). This distinction became generally more marginal as dose level increased to 3 mg/kg and 10 mg/kg. Notable compounds proving more amenable to oral administration included two representative carbonate prodrugs, C02 and C03. Both these molecules achieved psilocin C_max_ values and AUC_(0-tlast)_ measures comparable to psilocybin at both 3 mg/kg and 10 mg/kg dose levels. C03 also demonstrated an extended rate of psilocin elimination, or psilocin half-life (T_1/2_) (Table S1), suggesting elevated levels of parent prodrug may have been absorbed and metabolized. Interestingly, C02 and C03 contain non-polar sidechains (benzyl and cyclopropyl, respectively) that may increase lipophilicity and contribute to greater absorption when orally administered (37,38). T01 also contains a benzyl prodrug moiety, which may explain the favourable psilocin C_max_ and exposure at higher oral dose levels. Similarly, ester-linked aryl derivatives, ES03 and ES04, also produce slightly greater psilocin bioavailability upon oral administration. Indeed, a previously reported approach involving the addition of non-polar L-valine, L-isoleucine and L-phenylalanine esters to positively charged guanidino-containing molecules significantly enhanced enteric permeability of these compounds, ultimately enhancing oral delivery (39). In line with this potential for enhanced permeability, many of the NPDs were detected as circulating unmetabolized parent prodrug after oral delivery, a property not observed at any oral dose of psilocybin (Figure S8).

**Figure 2.**
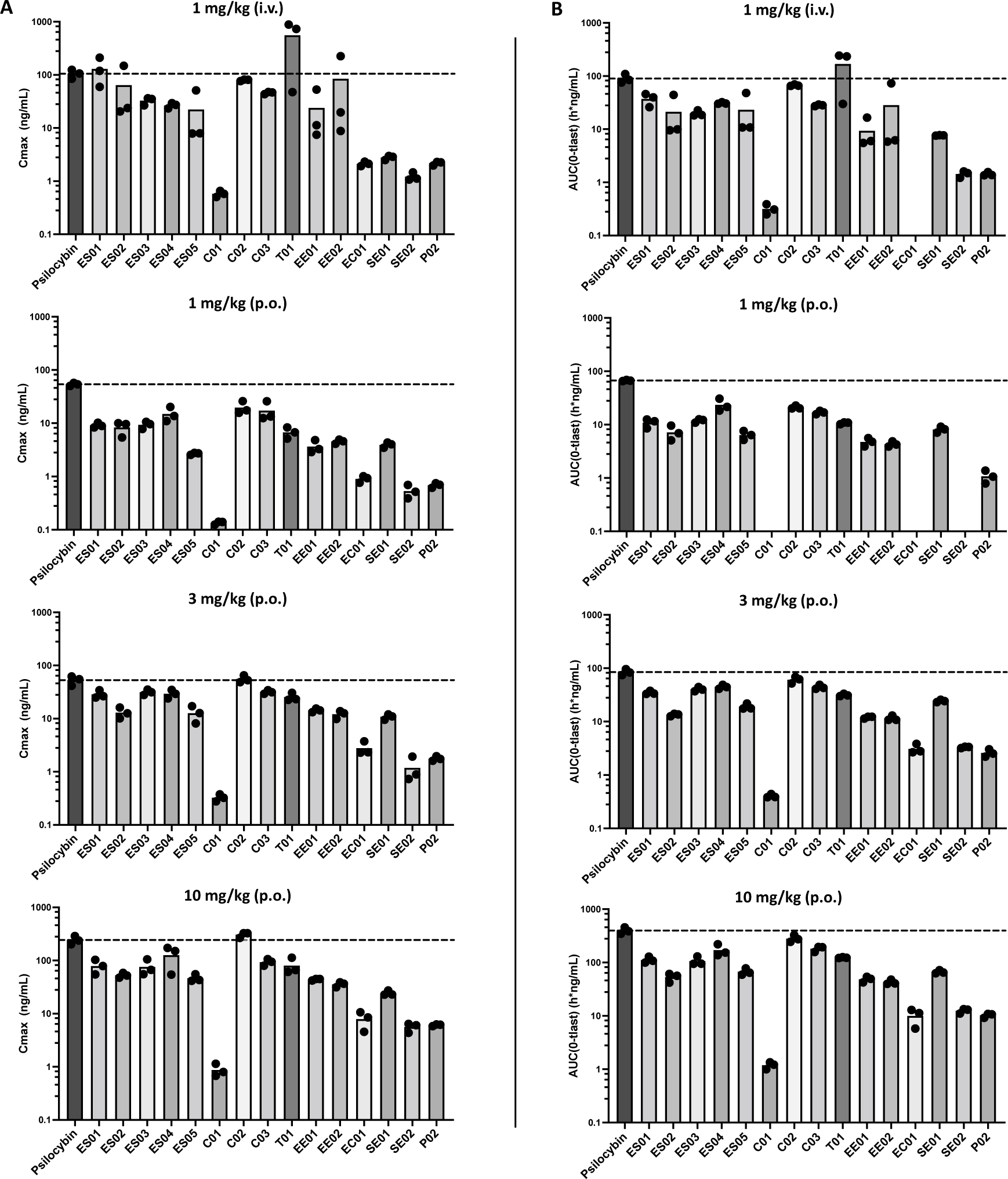
Pharmacokinetic (PK) analysis of plasma psilocin concentration and measurable psilocin exposure was evaluated in healthy C57BL/6 mice dosed with a single administration of psilocybin or an NDP. LC-MS-based quantification of resultant psilocin metabolite was performed at timepoints 0.25, 0.5, 1, 2, 4, 6, 8 and 24 h post dose. An additional serial sample was collected at 5 min for animals dosed i.v. A) Maximum plasma concentration (C_max_) achieved after administration of each prodrug compound is plotted for 1 mg/kg (i.v.), and 1, 3 and 10 mg/kg (p.o). B) Measurable plasma psilocin exposure (AUC_(0-tlast)_) achieved after administration of each prodrug compound is plotted for 1 mg/kg (i.v.), and 1, 3 and 10 mg/kg (p.o). n = 3 mice for all dose groups analyzed.

As discussed above, alteration to the cleavable prodrug moiety of psilocybin is predicted to produce variability in the metabolic profile of each NPD, ultimately modifying their PK and overall bioavailability. When dosed orally at both 3 and 10 mg/kg, psilocybin produces notably elevated psilocin C_max_ (52.9 ±6.03 and 243 ±25.2 ng/mL) and significantly enhanced psilocin exposure (AUC_(0-tlast)_ (84.4 ±6.24 and 397 ±31.9 h*ng/mL)) (Table S1). These metabolic characteristics lead to measurable levels of plasma psilocin detectable up to 24 h post-administration at both these oral dose levels (Figure S7). Two carbonate-based prodrugs C02 and C03 which maintain C_max_ and AUC_(0-tlast)_ values comparable to psilocybin (Table S1, Figure S7) also demonstrate detectable psilocin at 24 h post-dose when administered at 10 mg/kg. However, no other NPD produced detectable levels of psilocin when measured beyond 8 h after oral dosing, even when dosed as high as 10 mg/kg (p.o.). In fact, with the exception of C02 and C03, even when plasma levels of psilocin exceed those derived from psilocybin, the active metabolite is eliminated notably faster (Figure 3). These results strongly suggest that derivatization of the cleavable moiety at the 4-OH position does have a markedly prominent effect on the metabolic profile of these compounds. Variation at this position may potentially alter various pharmacological parameters such as absorption of parent molecules, physiological compartment for metabolic processing, and elimination of the psilocin metabolite.

**Figure 3.**
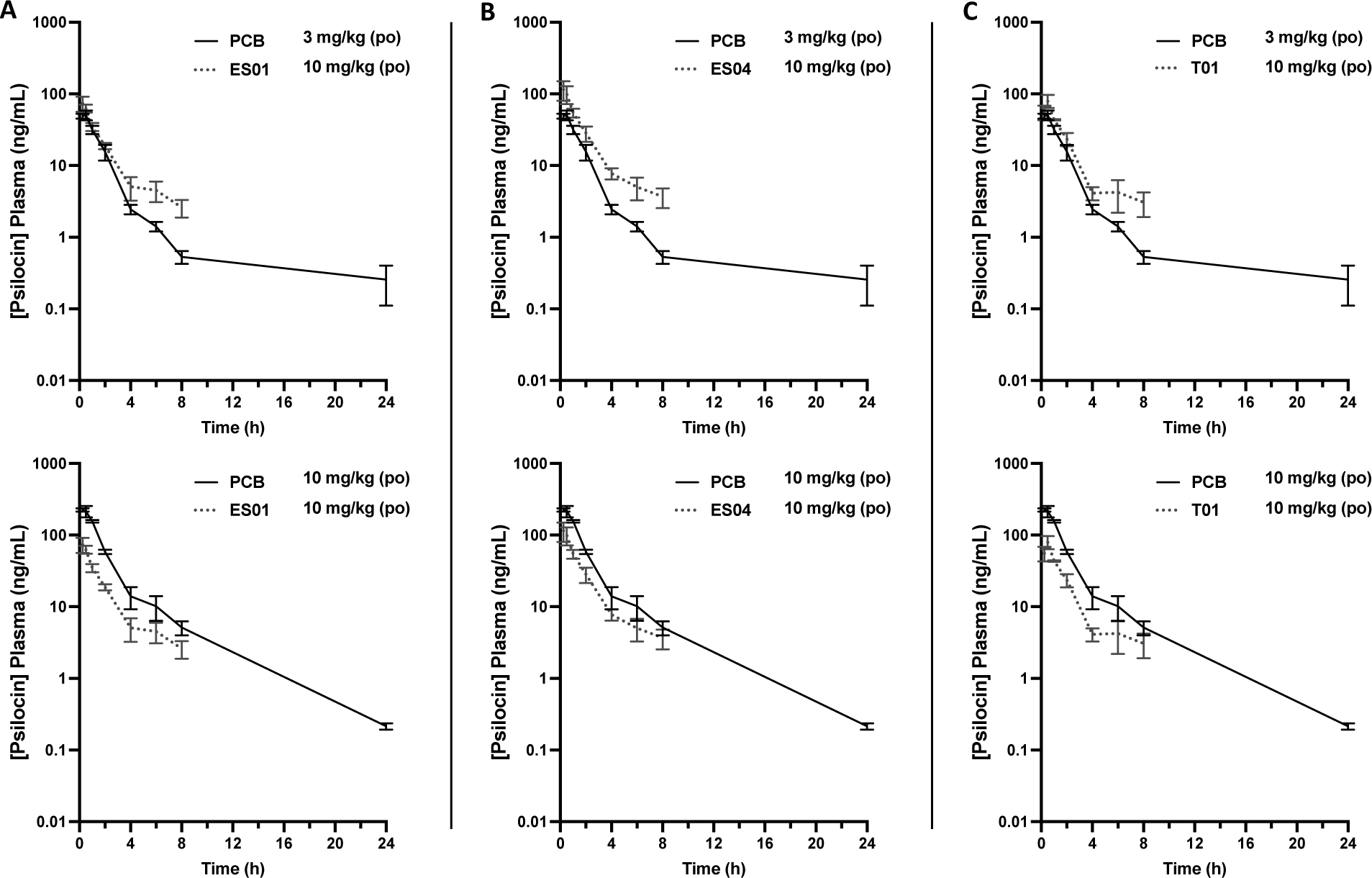
Comparison of plasma psilocin concentration profiles for A) ES01, B) ES04 and C) T01 dosed at 10 mg/kg relative to psilocybin (PCB) dosed at either 3 mg/kg (top plots) or 10 mg/kg (bottom plots). LC-MS-based quantification of psilocin metabolite was performed at timepoints 0.25, 0.5, 1, 2, 4, 6, 8 and 24 h post oral administration.

Overall, *in vitro* metabolic profiles generally conformed to the *in vivo* PK profiles for plasma psilocin, with the notable exception being psilocybin. It has been well documented that alkaline phosphatase (AP), which is considered the main enzyme contributing to the dephosphorylation of psilocybin post-absorption (see Figure S1) (4,17,25,26), is reported to be anchored to cellular plasma membranes in various tissues throughout the body (40,41,42,43,44). This membrane-localized feature of AP could explain the persistent stability of psilocybin when incubated in cell-free biological fractions *in vitro* (see Table 1, 2 and Figure S2), since these fractions are free of plasma membrane fragments. This data may also suggest that soluble esterases, known to be prevalent in these specific biological fractions, may not play a profound role in the metabolism of psilocybin.

### Head Twitch Response Behavioral Analysis

The Head Twitch Response (HTR) test in mice has proven to be a reliable behavioural proxy for the psychedelic activity produced by hallucinogenic drugs in humans (45–48). HTR, a rapid and involuntary side-to-side rotational head movement, is a distinct behavior induced in rodents upon exposure to psychedelic 5-HT_2A_ receptor agonists. Importantly, the degree of HTR has been shown to be strongly correlated to the psychoactive potency of administered hallucinogens, demonstrating significant predictive validity in this model. For our purposes, we employed the HTR mouse model to assess the functional bioavailability of systemic psilocin. There is strong translational precedence for this, as it has been demonstrated in rodents, pigs, and humans that plasma psilocin levels positively correlate with 5-HT_2A_ receptor occupancy and potency of the psychedelic response (29,49,50). In essence, varying levels of NPD-derived psilocin can be relatively determined as measures of HTR intensity, which can translate to overall prodrug metabolic efficiency *in vivo*. The same selection of NPDs assessed for their PK properties were further evaluated for psychedelic potency in healthy naïve mice at a well-tolerated oral dose of 1 mg/kg (p.o.). Psilocybin was also included in this test as an appropriate comparator. Relative to vehicle, psilocybin produced significantly enhanced HTR up to 90 min post-administration, with a peak intensity occurring within the 15-30 min observation window (Figure 4A). This profile closely correlates with the systemic levels of psilocybin-derived psilocin, which reaches a plasma C_max_ within this 15-min window (Figure S7). For screening purposes, this 15-min observation window was used to assess the HTR induced by each NPD dosed at the same level. Overall, ten of the NPDs assayed produced significant HTR above the vehicle-treated baseline (Figure 4B). Interestingly, despite producing peak plasma psilocin concentrations approximately five times lower than oral psilocybin (Table S1), three ester-based prodrugs (ES01, ES02 and ES03) induced HTR at an intensity statistically equivalent to the natural psychedelic compound. Two additional NPDs, T01 (thiocarbonate) and SE01 (silyl-ether) also induced notable HTR, each also producing lower peak psilocin levels. Interestingly, for most of these highlighted NPDs, lead-in *in vitro* and PK analyses produced results in-line with this physiological response, suggesting the potential predictability of these *in vitro* and pharmacokinetic measures.

**Figure 4.**
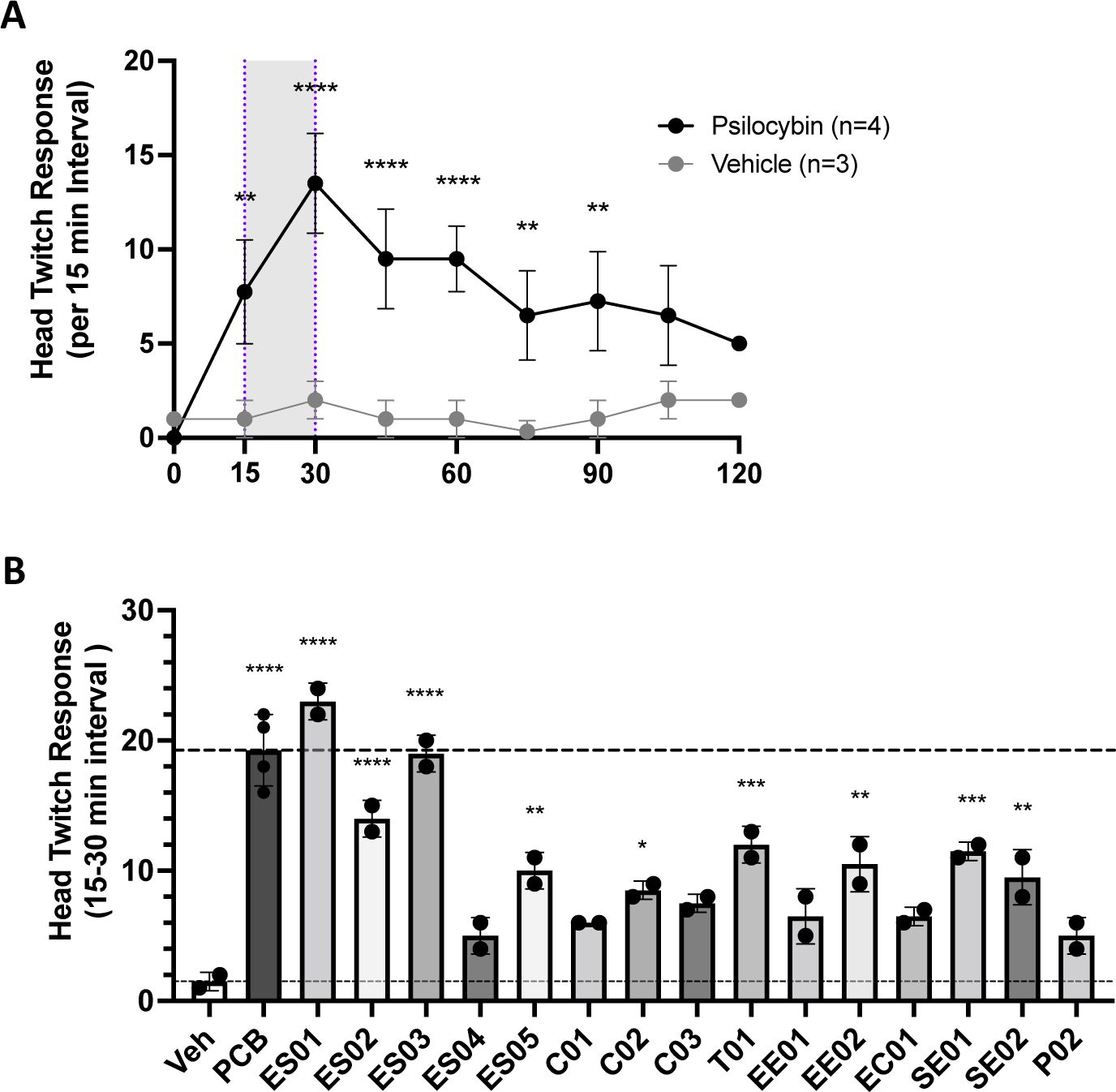
Evaluation of Head Twitch Response (HTR) in healthy C57BL/6 mice dosed with a single administration of psilocybin or an elected set of NPDs at 1 mg/kg (p.o.). A) Plot presents the time-course of HTR in mice orally dosed with 1 mg/kg psilocybin relative to mock-treated mice over a period of 120 min. HTR counts are plotted as number of counts per 15-min intervals and are marked on the plot at the end of the represented measurement period (n = 4 mice in the psilocybin group, n = 3 mice in the vehicle group). B) Plot presents the HTR count for the 15-30 min interval for mice treated with a single oral dose of vehicle, PCB, or an NPD at 1 mg/kg (n = 4 mice in the PCB group, n = 2 mice in all remaining groups). Statistical analysis conducted using ordinary one-way ANOVA: **** p ≤ 0.0001, *** p ≤ 0.001, ** p ≤ 0.01, * p ≤ 0.05.

Additionally, the carbonate-based NPD C01, with unfavourable *in vitro* and PK results, also confirmed to be ineffective in inducing HTR. In contrast, the silyl-ether based NPD, SE01, appears to have performed more favourably than the *in vitro* and PK data would have predicted. Conversely, NPDs ES04, C03 and EE01, each with *in vitro* and/or PK data suggestive of potent HTR outcomes, in fact appeared not to induce a notable hallucinogenic effect. Overall, these outcomes demonstrate that the appropriate assessment of a novel psychoactive entity requires a multi-modal approach to effectively ascertain true biofunctionality.

### Marble Burying model for Anxiolytic Potential

Numerous psychiatric mouse models have been developed to study anxiety and depression disorders (48,51,52). Many of these models have successfully demonstrated translational validity in the study of human neuropathology and therapeutic intervention. As critical as these *in vivo* models are for the development of novel psychotherapeutic treatments, there has also been some skepticism in their application due to the more complicated cognitive and emotional spectrum inherent in human neuropsychology. Additionally, there is the potential for overattributing innate behavioural characteristics, like burrowing and digging, as proxies for anxious behaviour (53,54). For example, the Marble Burying Test (MBT), regularly adopted as a model of anxiety and compulsive behaviours, has been successfully used to predict anxiolytic or anti-compulsive pharmacological potential (55–62). In fact, it has been presented that obsessive burying behaviour directed at specific sources of aversive or threatening stimuli may be a defensive activity driven by a state of anxiety created by these noxious objects (63). However, the activity of burrowing and nest building has also been used as an indicator of well-being in mice (64). Due to this potential ethological caveat, it has been strongly suggested that contextually valid experimental designs are needed to strengthen the translational value of studies like MBT (53). To this end, the application of a chronic stress paradigm, used to evoke extended hedonic deficit and other depressive-like symptoms in rodents, such as reduced motivation, increased aggression, anxiety, and overall reduction in well-being, is widely considered to have significant aetiological relevance and can serve as a valid preclinical model for various human psychiatric disorders (65–78).

To appropriately assess the anxiolytic potential of our NPDs we developed a seven-day mild chronic stress paradigm (MCSP) involving a combined oral corticosterone exposure (25 µg/mL introduced into drinking water for seven days) with a daily 30-min restraint stress session using a ventilated restrainer for the last five days of the protocol. It has been shown by others that the combination of these stress inducing methods leads to more severe depression and anxiety-like behaviours, producing earlier onset of cognitive deficit in conditioned mice (76). Indeed, when evaluated using the MBT, mice subjected to the MCSP exhibited a significant induction in marble burying behaviour over the unstressed (control) group (Figure 5). This stress conditioning did not produce any noticeable gender bias. Interestingly, this elevated level of stress persisted to the same extent 7 days after MCSP conditioning. It is important to note that this sustained cognitive deficiency was established after only seven days of chronic stress exposure, a period of induction considerably shorter than previously reported (75,76). Several studies have reported the successful treatment of severe anxiety after a single oral dose of psilocybin, especially in patients struggling with cancer-related distress (CRD) (13,79,80). As a demonstration of relevant anxiolytic efficacy in our model, mice subjected to the MCSP were treated with a single oral dose of 1 mg/kg psilocybin and assessed for induced marble burying behaviour (Figure 5). Psilocybin treatment resulted in significantly reduced marble burying behaviour in chronically stressed mice, completely returning them to their innate behavioural state, with long term benefit lasting up to seven days post treatment. Consistent with this result, Hesselgrave et. al. (2021) (81) reported robust recovery of anhedonic responses in mice subjected to an analogous chronic stress paradigm and treated with a single dose of psilocybin 24 h prior to behavioural assessment using the sucrose preference test (SPT) and female urine sniffing test (FUST). They also noted that stress-resilient mice, with no induced anhedonic responses, did not display loss of sucrose preference. Interestingly, our data also shows that psilocybin has no significant effect on the innate marble burying behaviour in non-stressed mice (Figure 5). This observation in MBT has also been previously reported by Matsushima et. al. (2009) (82). These combined results, demonstrating anti-depressive and/or anxiolytic effects only occur in behaviourally deficient mice, strongly suggest that the therapeutic value of psilocybin lies in recovery of a defective neurological condition induced by the chronic stress paradigm and not in an acute pharmacological alteration of perception. Indeed, this conclusion has also been reported by Hesselgrave et. al. (2021) (81), who have extended their analyses to reveal that psilocybin works through a specific mechanism of action to restore hippocampal synaptic strength in chronically stressed mice. This neuronal repair is strongly correlated to the recovery of anhedonia seen in these conditioned animals.

**Figure 5.**
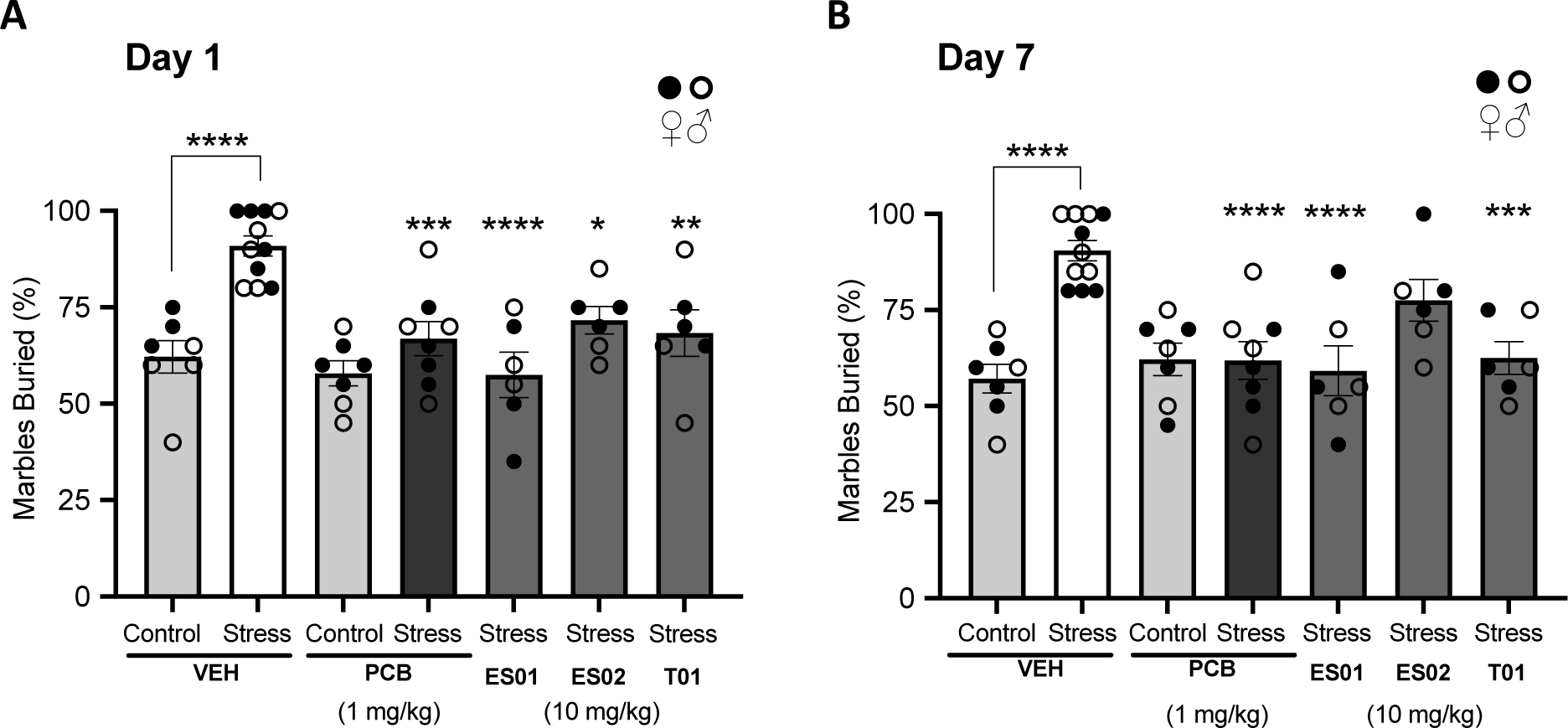
Results for the Marble Burying Test (MBT) in mice subjected to a Mild Chronic Stress Paradigm (MCSP) and treated with a single dose of vehicle, Psilocybin (PCB) at 1 mg/kg (p.o.) and the NPDs ES01, ES02 and T01 each at 10 mg/kg (p.o.). Plots present % marbles buried over a 20-min exposure period measured on A) Day 1 (24 h) and B) Day 7 (168 h) post-administration. Statistical analysis conducted using ordinary one-way ANOVA: **** p ≤ 0.0001, *** p ≤ 0.001, ** p ≤ 0.01, * p ≤ 0.05 (n = 7 mice for vehicle and psilocybin non-stress groups, n = 11 mice for vehicle stress group, n = 8 mice for psilocybin stress group, n = 6 mice for NPD stress groups).

Three NPDs, ES01, ES02 and T01, which demonstrated efficient prodrug properties and notable bioactivity, were elected for evaluation of anxiolytic potential using the MBT in mice subjected to the MCSP. To normalize the acute exposure of psilocin relative to that produced by 1 mg/kg oral psilocybin, each NPD was dosed orally at 10 mg/kg. All three NPDs tested induced a significant reduction in marble burying behaviour in stressed mice at 1-day post-dose (Figure 5). Two of these candidate prodrugs, ES01 (ester-based methoxybenzyl derivative) and T01 (thiocarbonate-based benzyl derivative) produced notable long-term positive outcomes, completely rescuing the elevated anxiety-like condition to the innate behavioural state up to seven days after treatment. Indeed, overall psilocin exposure may explain these outcomes, with these two prodrugs producing a more sustained level of psilocin with AUC_(0-tlast)_ values greater than twice that of ES02 within the first 8 h after administration.

## Discussion

Depression and anxiety disorders affect hundreds of millions of people worldwide and represent a major public health burden (World Health Organization [WHO], 2023). Numerous factors contribute to this global statistic. Current antidepressant and anxiolytic medications remain ineffective for a significant proportion of the afflicted population, demonstrating either eventual relapse or complete treatment resistance (83). Even when effective, antidepressant action of selective serotonin reuptake inhibitors (SSRIs) often results in reduced limbic responsiveness and emotional blunting (84). Alternatively, psychedelics such as psilocybin promote emotional release and have produced increased and sustained improvements in perceptual outlook, optimism and overall well-being in patients struggling with severe depression and anxiety disorders (7,8,10–13,79,80). Undesirably, the extended duration the acute psychoactive effect of psilocybin creates significant hurdles to the widespread therapeutic application of this natural psychedelic. The goal of the present study was to design a series of novel prodrug derivatives (NDPs) of psilocin, able to express therapeutically meaningful levels of this psychoactive agent, while maintaining altered pharmacokinetic profiles of psilocin plasma concentration, with modified overall psilocin exposure yet retained efficacious pharmacological responses.

### Chemical Modification at the 4-Position Alters Plasma Psilocin Exposure Profile

Chemical modifications of pharmacologically active agents into prodrug derivatives to improve their physiochemical, biopharmaceutical and pharmacokinetic properties is a well-established drug development strategy (reviewed in 85). As a natural prodrug of psilocin, psilocybin, by virtue of the phosphate moiety at the indolic 4-OH position, serves as a chemically stabilized form of this psychoactive molecule (24,25,86,87). As a soluble zwitterion, psilocybin maintains limited absorption potential; however, upon oral ingestion, a significant proportion of psilocybin is readily dephosphorylated to the more lipid-soluble psilocin, which is easily absorbed by the large and small intestinal tissues (6,17,27). Additional metabolism by intestinal and liver enzymes further enhances the bioavailability of psilocin (4,17,25,26). The primary objective of the current study was to generate a series of novel prodrugs (NPDs) of psilocin with diverse metabolically amendable substitutions at the 4-OH position. When orally dosed, all NPDs produced measurable levels of systemic psilocin (Figure 2, Table S1). Though these plasma psilocin levels are somewhat reduced relative to the mice dosed with psilocybin, most NPDs generated biofunctionally relevant psilocin concentrations, producing significant and timely stimulation of the 5-HT_2A_ receptor *in vivo* (Figure 4B). Also, with the exception of the carbonate-based prodrugs C02 and C03, which maintained C_max_ and AUC_(0-tlast)_ values comparable to psilocybin (Figure 2, Table S1), no other NPD produced detectable levels of psilocin beyond 8 h post administration even in mice dosed as high as 10 mg/kg (p.o.) (Figure S7). Notably, many NPDs harbouring non-polar, aromatic sidechains produced more systemic psilocin relative to the aliphatic and charged derivatives. This result suggests that these more favourable NPDs may possess potentially greater intestinal absorption properties. Additionally, it can be argued that the reduced level of plasma psilocin relative to that derived by an equivalent dose of psilocybin may suggest greater gastrointestinal (GI) stability of the orally dosed NPDs. This may prove to be a highly advantageous parameter. A commonly reported acute adverse effect of ingesting psilocybin is nausea, vomiting and occasional abdominal pain (7,9,11,88). Although the exact cause for this negative GI effect has not been appropriately elucidated, it is thought to be due to the rapid hydrolysis of psilocybin into the bioactive agent psilocin within the acidic lumen of the GI tract. Interestingly, serotonin signalling within the GI is a physiologically important function, and the majority of serotonin is synthesized, stored and released by enteroendocrine cells within the intestinal mucosa (reviewed in 89). Indeed, signals stimulating the perception of nausea and enteric pain are among the many effects of serotonin signalling within the GI. Hence, it may be reasonable to suggest that the presence of serotonergic psilocin following ingestion of psilocybin may have the potential to stimulate these reactive centres in the gut lining, inducing similar sensations of GI distress. As such, establishing greater GI stability and designing a metabolic mechanism reliant on post-absorption conversion would generate a notable therapeutic advantage by potentially reducing the adverse GI effects and producing a more acute systemic psilocin exposure.

### NPDs Generating Bioavailable Psilocin Stimulate Psychoactive Responses in Mice

Psychedelic-induced Head Twitch Response (HTR) in rodents has been reliably used as a behavioural pharmacodynamic (PD) biomarker of 5-HT_2A_ receptor stimulation and a proxy for psychoactive potential (47,48). In fact, it has been shown that plasma levels of psilocin strongly correlate to the intensity of this psychomimetic behaviour (49). Among the NPDs, orally dosed ester- and thiocarbonate-based derivatives produced the most potent HTR in healthy mice (Figure 4). Overall, these compounds reliably produced favourable *in vitro* and *in vivo* metabolic profiles relative to other NPDs, making these tests highly predictive of the intended behavioural outcome (Tables 1 and 2, Figure S7). It is important to note that, relative to psilocybin, each of these compounds produced less than 20% of the maximum psilocin concentration (C_max_) and exposure (AUC_(0-tlast)_). However, each were able to induce a psychoactive response statistically relevant to the natural prodrug. In contrast to the ester and thiocarbonate results, two silyl-ether based NPDs, SE01 and SE02, also induced notable HTR, although these compounds did not produce favourable metabolic profiles, with plasma psilocin levels at 7% and 1% that of an equivalent dose of psilocybin, respectively. These molecules, in fact, proved to be quite stable *in vivo*, with significant systemic levels of the parent derivatives detected up to 8 h post-dose (Figure S8). These results may implicate a more drawn-out, multistep conversion process for these molecules. The parent molecule in these compounds may have the potential to be used as novel chemical entities with psychoactive properties. Further investigation is required to characterize the biodistribution and functional nature for these NPDs.

Within our library of NPDs, a few analogous compound pairs were generated harbouring the same pendant linked by different cleavable moieties. As an example, the carbonate-based compound C02 and the thiocarbonate-based compound T01 both carry a benzyl sidechain R group (Figure 1). Interestingly, both these prodrugs performed differently when subjected to *in vitro* metabolism. T01 maintained much shorter half-life (T_1/2_) in each of the biological fractions tested compared to C02 (Table 1). Correspondingly, much greater psilocin was generated by T01, with a significant preference for liver-based *in vitro* metabolism (Table 2). In contrast to the variation seen *in vitro*, these two NPDs maintained very similar *in vivo* metabolic profiles, with both compounds producing notable levels of plasma psilocin and maintaining very similar overall psilocin exposure (Figure 2, Table S1). It should be noted that T01 performed much more favourably upon intravenous administration of 1 mg/kg, producing approximately seven times greater psilocin compared to C02. Indeed, T01 proved to be more metabolically amenable than even psilocybin when delivered systemically, producing approximately three to five times greater psilocin relative to the natural prodrug at the same dose. Finally, both compounds produced functional levels of plasma psilocin when dosed orally at 1 mg/kg, inducing statistically meaningful psychomimetic effects in healthy mice.

### NPDs Retain Anxiolytic Benefit in Chronically Stressed Mice

Alteration of the cleavable linker in a prodrug has profound effects on the metabolic profile of the modified compound. Changing the linker to attract a different target enzyme may not only diversify the stability of the molecule delivered by different routes of administration, but also alter the compartment responsible for the metabolic conversion of the prodrug. Psilocybin is known to be readily hydrolyzed in the acidic gastric compartment, releasing a significant level of psilocin prior to initial absorption. Our data has demonstrated that the substitution of an ester-based linker at the same 4-carbon position favours liver-based metabolic processing. Additionally, due to the lower levels of plasma psilocin produced when NPDs are orally dosed, our results imply a potentially greater stability in the gastro-intestinal (GI) lumen relative to psilocybin. The relative levels of plasma psilocin would be expected to increase if more of the orally dosed NPD was converted to psilocin within the intestinal lumen owing to favorable absorption properties. Therefore, greater GI stability creates a greater reliance on post-absorption metabolic processing, and potentially a more mediated acute psilocin exposure relative to psilocybin (Figure 3, Figure S7). Despite significant alteration to the pharmacokinetic profiles of these NPDs, we showed that fast acting and sustained anxiolytic benefit is retained by these candidate prodrugs. Moreover, they achieve efficacy comparable to psilocybin at dose-normalized levels (Figure 5). We note that, even when dosed at high levels, most NPDs maintain reduced overall exposure of psilocin compared to psilocybin, which may suggest a potentially reduced period of psychoactive effect, while maintaining the associated long-term benefits.

Though this needs to be fully explored, and may only be appropriately determined by human trial, the current study demonstrates the potential of using modified prodrug derivatives of psilocin to treat anxiety and depression. These newly described NPDs have the promise to dramatically improve the clinical application of psilocin as a therapeutic agent, offering alternative small-molecule pharmaceutical entities able to address current, treatment-resistant mental disorders.

## Methods

### Chemical Synthesis

All reactions were performed under argon or nitrogen atmosphere in flame-dried glassware and dried solvents at room temperature, unless stated otherwise. Controlled temperature reactions were performed using a mineral oil bath or aluminum heating blocks and a temperature controlled hot plate (IKA Ceramag Midi). Temperatures below room temperature were achieved in an ice/water bath (0 °C), dry ice/ethylene glycol bath (−20 °C), dry ice/ethanol/ethylene glycol bath (−20 °C to −75 °C) and dry ice/isopropanol bath (−78 °C). Solvents were removed under reduced pressure using a Büchi rotary evaporator. Anhydrous solvents were purchased from Sigma-Aldrich or prepared by distillation under nitrogen atmosphere or drying over 3 Å or 4 Å molecular sieves for at least 48 h. All reagents and solvents were purchased from Sigma-Aldrich, Combi-Blocks, Thermo Fisher, A2B Chemicals, Enamine, Oakwood Chemicals or Ambeed, unless stated otherwise. Thin-layer chromatography (TLC) was performed using silica gel 60 F254 precoated glass-backed plates. Detection of TLC spots was performed using a handheld UV lamp at 254 nm, or by staining with potassium permanganate prepared according to literature procedure. Flash column chromatography purifications were performed using silica gel 60 (230-400 mesh, Silicycle, Quebec) on automated Combi-Flash Teledyne NextGen 300 systems. Low-resolution mass spectra (LRMS) in heated electrospray ionization (HESI) mode were obtained from a Thermo Fortis, triple quadrupole (3Q) system. High-resolution mass spectra (HRMS) in heated electrospray ionization (HESI) mode were obtained from a Thermo LC-LTQ-Orbitrap-XL system. Proton (1H-NMR) spectra were obtained using Bruker RDQ-400 or DRY-400 (400 MHz) spectrometers. A detailed description of all chemical syntheses, including nuclear magnetic resonance spectra of all intermediates and products, is provided in Supplementary Information Dataset 1.

### *In vitro* Prodrug Metabolism

Human liver and intestinal microsomes and S9 fractions were purchased from XenoTech (xenotech.com). Human AB serum, purified bovine alkaline phosphatase, and purified porcine esterase were purchased from Sigma-Aldrich. For all *in vitro* metabolic stability assays, psilocybin and each NPD was prepared at an assay concentration of 2.5 µM in 50 mM potassium phosphate buffer (pH 7.4) containing 3 mM MgCl_2_, 1 mM EDTA, and supplemented with NADPH RapidStart Regeneration System (XenoTech) according to the instructions of the manufacturer. For assays involving biological fractions, each synthetic prodrug was incubated at 37 °C in 400 µg/mL of each S9 and microsomal fraction (based on total protein content), or in 10% serum fraction, in a total volume of 500 µL. For assays using purified enzymes, one unit of enzyme was added to prodrug compounds prepared as described above and incubated at 37 °C. Each molecule was also assessed for aqueous instability independent of enzyme activity in assay buffer alone. Metabolism of the parent prodrug compound into the psychoactive metabolite psilocin was monitored for each molecule every 20 min for a duration of 2 h. Briefly, 50 µL aliquots were drawn at the start of the assay (0 min), and then at 20, 40, 60, 80, 100 and 120 min thereafter. Aliquots were precipitated by adding one volume of acetonitrile to quench the reaction before centrifugation at 10,000 x g for 20 min. Supernatants were analyzed for the presence of target compound using LC-LTQ-Orbitrap-XL MS analysis (LC-MS) (ThermoScientific). Changes in analyte concentrations monitored over the time-course were plotted and analyzed using GraphPad Prism. Rate of metabolism was calculated by determining the *in vitro* metabolic half-life (T_1/2_) of the parent prodrug.

### Plasma Pharmacokinetics in Healthy Male C57BL/6 Mice

All plasma PK analyses were conducted under contract by InterVivo Solutions Inc (Toronto, ON) as per their standard protocol and procedures, and performed in accordance with the principles of the Canadian Council on Animal Care (CCAC). Male C57BL/6 mice (20-30 g) were used for all PK analyses. Briefly, psilocybin and 15 NPDs (as indicated in Figure 4) were prepared fresh in physiological saline at appropriate concentrations for dosing. 192 mice were randomly allocated into 64 groups (n = 3 per group). Each group received a single formulated dose of either psilocybin or an NPD administered at either 1 mg/kg intravenously (i.v.), or at 1, 3, or 10 mg/kg orally (p.o.) via gavage. Serial blood samples (∼0.03 mL) were collected in all mice via tail snip at 0.25, 0.5, 1, 2, 4, 6, 8, and 24 h post-dose. An additional serial sample was collected at 5 min for animals dosed i.v. Terminal blood samples were collected under isoflurane anesthesia via cardiac puncture at 24 h. All blood samples were transferred into K_2_EDTA tubes on wet ice and centrifuged within 5 min of collection at 3200 x g for 5 min at 4°C to obtain plasma. Plasma samples were stored at −80 °C until analysis. Parent prodrug and psilocin analyte concentrations were determined via liquid chromatography with tandem mass spectrometry (LC-MS-MS) using an AB Sciex QTRAP 4000 or 6000 MS/MS system as per InterVivo standard protocol.

### Animals Housing and Ethics Statement

All animal experimentations were approved by the University of Calgary Animal Care and Use Committee in accordance with Canadian Council on Animal Care (CCAC) guidelines. C57BL/6-Elite mixed sex mice were obtained from Charles River (8 weeks old). Upon arrival, mice were group-housed for 1 week on a 12:12 h light/dark schedule (lights on at 7:00 am) with *ad libitum* access to food and water. Three days prior to any behavioural protocol mice were handled for 5 to 10 min and exposed to the testing chamber for at least 5 min each day.

### Head Twitch Response Behavioural Assessment

Head twitch response (HTR) in mice, a rapid, involuntary movement of the head with little or no involvement of the trunk, has proven to be a reliable behavioural proxy for the psychedelic activity produced by hallucinogenic drugs in humans (13,45). Before any behavioral screening, mice were singly housed for two days prior to behavioural assessment and habituated to the experimental room 1 h before testing. The testing chamber was cleaned with a 70% (v/v) ethanol solution between experiments to eliminate odor from other mice. Test articles (Psilocybin and NPD compounds) freshly prepared as stock solutions at 100 mM in DMSO, were each diluted in sterile 0.9% (w/v) saline solution to an appropriate concentration for a dose of 1 mg/kg. Vehicle formulation consisted of 1% DMSO in 0.9% saline (w/v). Prior to oral administration, mice were video monitored for 30 min in a plexiglass testing chamber (25.5 x 12.5 x 12.5 cm [length x width x height]) to allow for acclimation to testing environment and to examine pre-drug spontaneous HTR and behavior. Following test article administration, mice were video monitored for 60 min and returned to their home cage. Behavioral analysis was conducted by an individual blinded to subject treatment group using Behavioral Observation Research Interactive Software (BORIS, version 7, DOI: 10.1111/2041-210X.12584). Post-dose behavior for psilocybin was analyzed for every 15-min interval to establish a control HTR profile for the full 60-min evaluation (n = 4 mice for psilocybin group, n = 3 for vehicle group). HTR for each NPD was reported for the 15-to-30-min window following test article administration (n = 4 mice for psilocybin treated groups; n = 2 for all other groups analyzed). Statistical analysis conducted using by one-way ANOVA using GraphPad Prism 9.3.1 statistical analysis software.

### Marble Burying Test coupled with a Combined Mild Chronic Stress Protocol

The Marble Burying Test (MBT) has been successfully used to predict anxiolytic or anti-compulsive pharmacological potential (55–62). This test has been most effective when coupled to a chronic stress paradigm (76). We developed a 7-day mild chronic stress paradigm (MCSP) involving a combined oral corticosterone exposure with a daily 30-min restraint stress session. Two days prior to the start of the MCSP mice were singly housed. For chronic corticosterone-induced stress, mice were exposed to this steroid hormone in their drinking water at 25 µg/mL in 1% ethanol for the full 7-day paradigm (vehicle cohort received water containing 1% ethanol alone). During the last 5 days of the protocol, mice were subjected to a daily 30-min restraint stress session using a ventilated restrainer fashioned from a conical tube. During the last two days of the protocol mice were exposed to the plexiglass test chamber without marbles for 10 min to reduce the impact of cage novelty. Test articles (Psilocybin and NPD compounds), freshly prepared as stock solutions at 100 mM in DMSO, were each diluted in sterile 0.9% (w/v) saline solution at a dose volume equivalent to 2.5 µL/g body weight, to establish an appropriate concentration for a dose of 1 mg/kg (psilocybin) or 10 mg/kg (NPDs). Vehicle formulation consisted of 1% DMSO in 0.9% saline (w/v). All test articles were administered as a single dose (p.o.) 24 h after the MCSP. MBT was conducted on day 1 and day 7 post-administration. Briefly, on each testing day, 10 glass marbles (approximately 15 mm in diameter) were evenly spaced on an even 5 cm layer of wood-chip bedding. Each vehicle and conditioned mouse were placed individually in a prepared test chamber and allowed to explore for 20 min, then returned to their home cage. For analysis, fully buried marbles were scored as 1 marbled buried and partially buried marbles are scored as 0.5 marbles buried. A final assessment of % marbles buried was reported. Statistical analysis was conducted by ordinary one-way ANOVA using GraphPad Prism 9.3.1 statistical analysis software (n = 7 mice for vehicle and psilocybin non-stress groups, n = 11 mice for vehicle stress group, n = 8 mice for psilocybin stress group, n = 6 mice for NPD stress groups).

## Acknowledgements

We are grateful to the Department of Biological Sciences and the Faculty of Science at the University of Calgary for the use of research facilities. Research funding was provided by Enveric Biosciences, Inc. to the University of Calgary with J.S.B. and P.J.F. as principal subproject investigators.

## Author contributions

S.A.R. wrote the manuscript and coordinated the pharmacology work; J.M.H. proposed some of the compounds and coordinated the biochemistry work; K.M. and D.P. proposed some of the compounds, designed the chemical syntheses and performed some synthesis; G.J., J.B.L. and C.C. performed some synthesis; S.G.C. performed the mouse behavioral assays; J.G. performed the LC-MS analyses; G.S., L.Y. and D.D., performed the pharmacological analyses; J.S.B. supervised the mouse behavioral assays; J.E.T. conceived the project; P.J.F. supervised the project; all authors have read and approved the content of the paper.

## Competing interests

All authors receive compensation from, and S.A.R., J.M.H., K.M., L.Y., D.P., G.S., D.D., G.J., J.B.L., C.C., J.G., J.E.T., and P.J.F. hold equity in, Enveric Biosciences, Inc. A United States patent application related to this work has been allowed (17/893,122).

**Correspondence** and requests for materials should be addressed to Peter Facchini.

## References

1. Nutt, D.; Erritzoe, D.; Carhart-Harris, R. Psychedelic Psychiatry’s Brave New World. Cell 2020, 181 (1), 24–28. https://doi.org/10.1016/j.cell.2020.03.020.

2. Rucker, J. J. H.; Iliff, J.; Nutt, D. J. Psychiatry & the Psychedelic Drugs. Past, Present & Future. Neuropharmacology 2018, 142, 200–218. https://doi.org/10.1016/j.neuropharm.2017.12.040.

3. Guzmán, G. Hallucinogenic Mushrooms in Mexico: An Overview. Economic Botany 2008, 62 (3), 404–412. https://doi.org/10.1007/s12231-008-9033-8.

4. Passie, T.; Seifert, J.; Schneider, U.; Emrich, H. M. The Pharmacology of Psilocybin. Addiction Biology 2002, 7 (4), 357–364. https://doi.org/10.1080/1355621021000005937.

5. Wieczorek, P. P.; Witkowska, D.; Jasicka-Misiak, I.; Poliwoda, A.; Oterman, M.; Zielińska, K. Bioactive Alkaloids of Hallucinogenic Mushrooms. Studies in Natural Products Chemistry 2015, 46, 133–168. https://doi.org/10.1016/b978-0-444-63462-7.00005-1.

6. Brown, R. T.; Nicholas, C. R.; Cozzi, N. V.; Gassman, M. C.; Cooper, K. M.; Muller, D.; Thomas, C. D.; Hetzel, S. J.; Henriquez, K. M.; Ribaudo, A. S.; Hutson, P. R. Pharmacokinetics of Escalating Doses of Oral Psilocybin in Healthy Adults. Clinical pharmacokinetics 2017, 56 (12), 1543–1554. https://doi.org/10.1007/s40262-017-0540-6.

7. Carhart-Harris, R. L.; Bolstridge, M.; Rucker, J.; Day, C. M. J.; Erritzoe, D.; Kaelen, M.; Bloomfield, M.; Rickard, J. A.; Forbes, B.; Feilding, A.; Taylor, D.; Pilling, S.; Curran, V. H.; Nutt, D. J. Psilocybin with Psychological Support for Treatment-Resistant Depression: An Open-Label Feasibility Study. The Lancet Psychiatry 2016, 3 (7), 619–627. https://doi.org/10.1016/s2215-0366(16)30065-7.

8. Carhart-Harris, R. L.; Goodwin, G. M. The Therapeutic Potential of Psychedelic Drugs: Past, Present, and Future. Neuropsychopharmacology 2017, 42 (11), 2105–2113. https://doi.org/10.1038/npp.2017.84.

9. Carhart-Harris, R. L.; Bolstridge, M.; Day, C. M. J.; Rucker, J.; Watts, R.; Erritzoe, D. E.; Kaelen, M.; Giribaldi, B.; Bloomfield, M.; Pilling, S.; Rickard, J. A.; Forbes, B.; Feilding, A.; Taylor, D.; Curran, H. V.; Nutt, D. J. Psilocybin with Psychological Support for Treatment-Resistant Depression: Six-Month Follow-Up. Psychopharmacology 2017, 235 (2), 399–408. https://doi.org/10.1007/s00213-017-4771-x.

10. Carhart-Harris, R.; Giribaldi, B.; Watts, R.; Baker-Jones, M.; Murphy-Beiner, A.; Murphy, R.; Martell, J.; Blemings, A.; Erritzoe, D.; Nutt, D. J. Trial of Psilocybin versus Escitalopram for Depression. New England Journal of Medicine 2021, 384 (15), 1402–1411. https://doi.org/10.1056/nejmoa2032994.

11. Davis, A. K.; Barrett, F. S.; May, D. G.; Cosimano, M. P.; Sepeda, N. D.; Johnson, M. W.; Finan, P. H.; Griffiths, R. R. Effects of Psilocybin-Assisted Therapy on Major Depressive Disorder. JAMA Psychiatry 2020, 78 (5). https://doi.org/10.1001/jamapsychiatry.2020.3285.

12. Gukasyan, N.; Davis, A. K.; Barrett, F. S.; Cosimano, M. P.; Sepeda, N. D.; Johnson, M. W.; Griffiths, R. R. Efficacy and Safety of Psilocybin-Assisted Treatment for Major Depressive Disorder: Prospective 12-Month Follow-Up. Journal of Psychopharmacology 2022, 36 (2), 151–158. https://doi.org/10.1177/02698811211073759.

13. Griffiths, R. R.; Johnson, M. W.; Carducci, M. A.; Umbricht, A.; Richards, W. A.; Richards, B. D.; Cosimano, M. P.; Klinedinst, M. A. Psilocybin Produces Substantial and Sustained Decreases in Depression and Anxiety in Patients with Life-Threatening Cancer: A Randomized Double-Blind Trial. Journal of Psychopharmacology 2016, 30 (12), 1181–1197. https://doi.org/10.1177/0269881116675513.

14. Griffiths, R. R.; Richards, W. A.; McCann, U.; Jesse, R. Psilocybin Can Occasion Mystical-Type Experiences Having Substantial and Sustained Personal Meaning and Spiritual Significance. Psychopharmacology 2006, 187 (3), 268–283. https://doi.org/10.1007/s00213-006-0457-5.

15. Hasler, F.; Grimberg, U.; Benz, M. A.; Huber, T.; Vollenweider, F. X. Acute Psychological and Physiological Effects of Psilocybin in Healthy Humans: A Double-Blind, Placebo-Controlled Dose–Effect Study. Psychopharmacology 2004, 172, 145–156. https://doi.org/10.1007/s00213-003-1640-6.

16. Madsen, M. K.; Fisher, P. M.; Burmester, D.; Dyssegaard, A.; Stenbæk, D. S.; Kristiansen, S.; Johansen, S. S.; Lehel, S.; Linnet, K.; Svarer, C.; Erritzoe, D.; Ozenne, B.; Knudsen, G. M. Psychedelic Effects of Psilocybin Correlate with Serotonin 2A Receptor Occupancy and Plasma Psilocin Levels. Neuropsychopharmacology 2019, 44 (7), 1328–1334. https://doi.org/10.1038/s41386-019-0324-9.

17. Hasler, F.; Bourquin, D.; Brenneisen, R.; Bär, T.; Vollenweider, F. X. Determination of Psilocin and 4-Hydroxyindole-3-Acetic Acid in Plasma by HPLC-ECD and Pharmacokinetic Profiles of Oral and Intravenous Psilocybin in Man. Pharmaceutica Acta Helvetiae 1997, 72 (3), 175–184. https://doi.org/10.1016/s0031-6865(97)00014-9.

18. Garcia-Romeu, A.; Barrett, F. S.; Carbonaro, T. M.; Johnson, M. W.; Griffiths, R. R. Optimal Dosing for Psilocybin Pharmacotherapy: Considering Weight-Adjusted and Fixed Dosing Approaches. Journal of Psychopharmacology 2021, 35 (4), 353–361. https://doi.org/10.1177/0269881121991822.

19. Rautio, J.; Meanwell, N. A.; Di, L.; Hageman, M. J. The Expanding Role of Prodrugs in Contemporary Drug Design and Development. Nature Reviews Drug Discovery 2018, 17 (8), 559–587. https://doi.org/10.1038/nrd.2018.46.

20. Shirota, O.; Hakamata, W.; Goda, Y. Concise Large-Scale Synthesis of Psilocin and Psilocybin, Principal Hallucinogenic Constituents of “Magic Mushroom.” Journal of Natural Products 2003, 66 (6), 885–887. https://doi.org/10.1021/np030059u.

21. Matinkhoo, K.; Pryyma, A.; Todorovic, M.; Patrick, B. O.; Perrin, D. M. Synthesis of the Death-Cap Mushroom Toxin α-Amanitin. Journal of the American Chemical Society 2018, 140 (21), 6513–6517. https://doi.org/10.1021/jacs.7b12698.

22. Horita, A.; Weber, L. J. Dephosphorylation of Psilocybin to Psilocin by Alkaline Phosphatase. Experimental Biology and Medicine 1961, 106 (1), 32–34. https://doi.org/10.3181/00379727-106-26228.

23. Horita, A.; Weber, L. J. The Enzymic Dephosphorylation and Oxidation of Psilocybin and Pscilocin by Mammalian Tissue Homogenates. Biochemical Pharmacology 1961, 7 (1), 47–54. https://doi.org/10.1016/0006-2952(61)90124-1.

24. Horita, A.; Weber, L. J. Dephosphorylation of Psilocybin in the Intact Mouse. Toxicology and Applied Pharmacology 1962, 4 (6), 730–737. https://doi.org/10.1016/0041-008x(62)90102-3.

25. Horita, A. SOME BIOCHEMICAL STUDIES on PSILOCYBIN and PSILOCIN. Journal of Neuropsychiatry and Clinical Neurosciences 1962, 4 (6). https://doi.org/10.21236/ad0291057.

26. Dinis-Oliveira, R. J. Metabolism of Psilocybin and Psilocin: Clinical and Forensic Toxicological Relevance. Drug metabolism reviews 2017, 49 (1), 84–91. https://doi.org/10.1080/03602532.2016.1278228.

27. K Eivindvik; Rasmussen, K. Ø.; Sund, R. B. Handling of Psilocybin and Psilocin by Everted Sacs of Rat Jejunum and Colon. Acta pharmaceutica nordica 1989, 1 (5), 295–302.

28. Lenz, C.; Dörner, S.; Trottmann, F.; Hertweck, C.; Sherwood, A.; Hoffmeister, D. Assessment of Bioactivity-Modulating Pseudo-Ring Formation in Psilocin and Related Tryptamines. ChemBioChem 2022, 23 (13). https://doi.org/10.1002/cbic.202200183.

29. Madsen, M. K.; Knudsen, G. M. Plasma Psilocin Critically Determines Behavioral and Neurobiological Effects of Psilocybin. Neuropsychopharmacology 2020, 46, 257–258. https://doi.org/10.1038/s41386-020-00823-4.

30. Yau, E.; Petersson, C.; Dolgos, H.; Peters, S. A. A Comparative Evaluation of Models to Predict Human Intestinal Metabolism from Nonclinical Data. Biopharmaceutics & Drug Disposition 2017, 38 (3), 163–186. https://doi.org/10.1002/bdd.2068.

31. Yan, Z.; Caldwell, G. W. Optimization in Drug Discovery; Springer Science & Business Media, 2008.

32. Zanger, U. M.; Schwab, M. Cytochrome P450 Enzymes in Drug Metabolism: Regulation of Gene Expression, Enzyme Activities, and Impact of Genetic Variation. Pharmacology & Therapeutics 2013, 138 (1), 103–141. https://doi.org/10.1016/j.pharmthera.2012.12.007.

33. Zhang, H.; Gao, N.; Tian, X.; Liu, T.; Fang, Y.; Zhou, J.; Wen, Q.; Xu, B.; Qi, B.; Gao, J.; Li, H.; Jia, L.; Qiao, H. Content and Activity of Human Liver Microsomal Protein and Prediction of Individual Hepatic Clearance in Vivo. Scientific Reports 2015, 5 (1). https://doi.org/10.1038/srep17671.

34. Casey Laizure, S.; Herring, V.; Hu, Z.; Witbrodt, K.; Parker, R. B. The Role of Human Carboxylesterases in Drug Metabolism: Have We Overlooked Their Importance? Pharmacotherapy: The Journal of Human Pharmacology and Drug Therapy 2013, 33 (2), 210–222. https://doi.org/10.1002/phar.1194.

35. J. Richardson, S.; Bai, A. A. Kulkarni, A.F. Moghaddam, M. Efficiency in Drug Discovery: Liver S9 Fraction Assay as a Screen for Metabolic Stability. Drug Metabolism Letters 2016, 10 (2), 83–90. https://doi.org/10.2174/1872312810666160223121836.

36. Peeters, L.; Vervliet, P.; Foubert, K.; Hermans, N.; Pieters, L.; Covaci, A. A Comparative Study on the in Vitro Biotransformation of Medicagenic Acid Using Human Liver Microsomes and S9 Fractions. Chemico-Biological Interactions 2020, 328, 109192. https://doi.org/10.1016/j.cbi.2020.109192.

37. Lipinski, C. A.; Lombardo, F.; Dominy, B. W.; Feeney, P. J. Experimental and Computational Approaches to Estimate Solubility and Permeability in Drug Discovery and Development Settings. Advanced Drug Delivery Reviews 1997, 23 (1-3), 3–25. https://doi.org/10.1016/s0169-409x(96)00423-1.

38. Azman, M.; Sabri, A. H.; Anjani, Q. K.; Mustaffa, M. F.; Hamid, K. A. Intestinal Absorption Study: Challenges and Absorption Enhancement Strategies in Improving Oral Drug Delivery. Pharmaceuticals 2022, 15 (8), 975. https://doi.org/10.3390/ph15080975.

39. Sun, J.; Dahan, A.; Amidon, G. L. Enhancing the Intestinal Absorption of Molecules Containing the Polar Guanidino Functionality: A Double-Targeted Prodrug Approach. Journal of Medicinal Chemistry 2010, 53 (2), 624–632. https://doi.org/10.1021/jm9011559.

40. Sharma, U.; Pal, D.; Prasad, R. Alkaline Phosphatase: An Overview. Indian Journal of Clinical Biochemistry 2013, 29 (3), 269–278. https://doi.org/10.1007/s12291-013-0408-y.

41. Zhang, D.; Surapaneni, S. ADME-Enabling Technologies in Drug Design and Development; John Wiley & Sons, 2012.

42. De Broe, M. E.; Roels, F.; Nouwen, E. J.; Claeys, L.; Wieme, R. J. Liver Plasma Membrane: The Source of High Molecular Weight Alkaline Phosphatase in Human Serum. Hepatology 1985, 5 (1), 118–128. https://doi.org/10.1002/hep.1840050124.

43. Tokumitsu, S.; Tokumitsu, K.; Fishman, W. H. Immunocytochemical Demonstration of Intracytoplasmic Alkaline Phosphatase in HeLa TCRC-1 Cells. Journal of Histochemistry & Cytochemistry 1981, 29 (9), 1080–1087. https://doi.org/10.1177/29.9.7026668.

44. Lin, C. W.; Sasaki, M.; Orcutt, M. L.; Miyayama, H.; Singer, R. M. Plasma Membrane Localization of Alkaline Phosphatase in HeLa Cells. Journal of Histochemistry & Cytochemistry 1976, 24 (5), 659–667. https://doi.org/10.1177/24.5.58927.

45. Gonzalez-Maeso J.; Weisstaub, N. V.; Zhou, M.; Chan, P.; Ivic, L.; Ang, R.; Lira, A.; Bradley-Moore, M.; Ge, Y.; Zhou, Q.; Sealfon, S. C.; Gingrich, J. A. Hallucinogens Recruit Specific Cortical 5-HT2A Receptor-Mediated Signaling Pathways to Affect Behavior. Neuron 2007, 53 (3), 439–452. https://doi.org/10.1016/j.neuron.2007.01.008.

46. Halberstadt, A. L.; Geyer, M. A. Characterization of the Head-Twitch Response Induced by Hallucinogens in Mice. Psychopharmacology 2013, 227 (4), 727–739. https://doi.org/10.1007/s00213-013-3006-z.

47. Halberstadt, A. L.; Chatha, M.; Klein, A. K.; Wallach, J.; Brandt, S. D. Correlation between the Potency of Hallucinogens in the Mouse Head-Twitch Response Assay and Their Behavioral and Subjective Effects in Other Species. Neuropharmacology 2020, 167, 107933. https://doi.org/10.1016/j.neuropharm.2019.107933.

48. Hanks, J. B.; González-Maeso, J. Animal Models of Serotonergic Psychedelics. ACS Chemical Neuroscience 2012, 4 (1), 33–42. https://doi.org/10.1021/cn300138m.

49. Higgins, G. A.; Carroll, N. K.; Brown, M.; MacMillan, C.; Silenieks, L. B.; Thevarkunnel, S.; Izhakova, J.; Magomedova, L.; DeLannoy, I.; Sellers, E. M. Low Doses of Psilocybin and Ketamine Enhance Motivation and Attention in Poor Performing Rats: Evidence for an Antidepressant Property. Frontiers in Pharmacology 2021, 12. https://doi.org/10.3389/fphar.2021.640241.

50. Donovan, L. L.; Johansen, J. V.; Ros, N. F.; Jaberi, E.; Linnet, K.; Johansen, S. S.; Ozenne, B.; Issazadeh-Navikas, S.; Hansen, H. D.; Knudsen, G. M. Effects of a Single Dose of Psilocybin on Behaviour, Brain 5-HT2A Receptor Occupancy and Gene Expression in the Pig. European Neuropsychopharmacology 2021, 42, 1–11. https://doi.org/10.1016/j.euroneuro.2020.11.013.

51. Wang, Q.; Timberlake, M. A.; Prall, K.; Dwivedi, Y. The Recent Progress in Animal Models of Depression. Progress in Neuro-Psychopharmacology and Biological Psychiatry 2017, 77, 99–109. https://doi.org/10.1016/j.pnpbp.2017.04.008.

52. Bouguiyoud, N.; Roullet, F.; Bronchti, G.; Frasnelli, J.; Al Aïn, S. Anxiety and Depression Assessments in a Mouse Model of Congenital Blindness. Frontiers in Neuroscience 2022, 15. https://doi.org/10.3389/fnins.2021.807434.

53. de Brouwer, G.; Fick, A.; Harvey, B. H.; Wolmarans, D. W. A Critical Inquiry into Marble-Burying as a Preclinical Screening Paradigm of Relevance for Anxiety and Obsessive– Compulsive Disorder: Mapping the Way Forward. Cognitive, Affective, & Behavioral Neuroscience 2018, 19 (1), 1–39. https://doi.org/10.3758/s13415-018-00653-4.

54. Thomas, A.; Burant, A.; Bui, N.; Graham, D.; Yuva-Paylor, L. A.; Paylor, R. Marble Burying Reflects a Repetitive and Perseverative Behavior More than Novelty-Induced Anxiety. Psychopharmacology 2009, 204 (2), 361–373. https://doi.org/10.1007/s00213-009-1466-y.

55. Broekkamp, C. L.; Rijk, H. W.; Joly-Gelouin, D.; Lloyd, K. L. Major Tranquillizers Can Be Distinguished from Minor Tranquillizers on the Basis of Effects on Marble Burying and Swim-Induced Grooming in Mice. European Journal of Pharmacology 1986, 126 (3), 223–229. https://doi.org/10.1016/0014-2999(86)90051-8.

56. Nicolas, L. B.; Kolb, Y.; Prinssen, E. P. M. A Combined Marble Burying–Locomotor Activity Test in Mice: A Practical Screening Test with Sensitivity to Different Classes of Anxiolytics and Antidepressants. European Journal of Pharmacology 2006, 547 (1-3), 106–115. https://doi.org/10.1016/j.ejphar.2006.07.015.

57. Egashira, N.; Kubota, N.; Goto, Y.; Watanabe, T.; Kubota, K.; Katsurabayashi, S.; Iwasaki, K. The Antipsychotic Trifluoperazine Reduces Marble-Burying Behavior in Mice via D 2 and 5-HT 2A Receptors: Implications for Obsessive–Compulsive Disorder. Pharmacology Biochemistry and Behavior 2018, 165, 9–13. https://doi.org/10.1016/j.pbb.2017.12.006.

58. Taylor, G. T.; Lerch, S.; Chourbaji, S. Marble Burying as Compulsive Behaviors in Male and Female Mice. Acta Neurobiologiae Experimentalis 2017, 77 (3), 254–260. https://doi.org/10.21307/ane-2017-059.

59. Angoa-Pérez, M.; Kane, M. J.; Briggs, D. I.; Francescutti, D. M.; Kuhn, D. M. Marble Burying and Nestlet Shredding as Tests of Repetitive, Compulsive-like Behaviors in Mice. Journal of Visualized Experiments 2013, No. 82. https://doi.org/10.3791/50978.

60. Dixit, P. V.; Sahu, R.; Mishra, D. K. Marble-Burying Behavior Test as a Murine Model of Compulsive-like Behavior. Journal of Pharmacological and Toxicological Methods 2020, 102, 106676. https://doi.org/10.1016/j.vascn.2020.106676.

61. Lustberg, D.; Iannitelli, A. F.; Tillage, R. P.; Pruitt, M.; Liles, L. C.; Weinshenker, D. Central Norepinephrine Transmission Is Required for Stress-Induced Repetitive Behavior in Two Rodent Models of Obsessive-Compulsive Disorder. Psychopharmacology 2020, 237 (7), 1973–1987. https://doi.org/10.1007/s00213-020-05512-0.

62. Sugimoto, Y.; Tagawa, N.; Kobayashi, Y.; Hotta, Y.; Yamada, J. Effects of the Serotonin and Noradrenaline Reuptake Inhibitor (SNRI) Milnacipran on Marble Burying Behavior in Mice. Biological and Pharmaceutical Bulletin 2007, 30 (12), 2399–2401. https://doi.org/10.1248/bpb.30.2399.

63. De Boer, S. F.; Koolhaas, J. M. Defensive Burying in Rodents: Ethology, Neurobiology and Psychopharmacology. European Journal of Pharmacology 2003, 463 (1-3), 145–161. https://doi.org/10.1016/s0014-2999(03)01278-0.

64. Jirkof, P. Burrowing and Nest Building Behavior as Indicators of Well-Being in Mice. Journal of Neuroscience Methods 2014, 234, 139–146. https://doi.org/10.1016/j.jneumeth.2014.02.001.

65. Strekalova, T.; Couch, Y.; Kholod, N.; Boyks, M.; Malin, D.; Leprince, P.; Steinbusch, H. M. Update in the Methodology of the Chronic Stress Paradigm: Internal Control Matters. Behavioral and Brain Functions 2011, 7 (1), 9. https://doi.org/10.1186/1744-9081-7-9.

66. Willner, P. Validity, Reliability and Utility of the Chronic Mild Stress Model of Depression: A 10-Year Review and Evaluation. Psychopharmacology 1997, 134 (4), 319–329. https://doi.org/10.1007/s002130050456.

67. Willner, P.; Muscat, R.; Papp, M. Chronic Mild Stress-Induced Anhedonia: A Realistic Animal Model of Depression. Neuroscience & Biobehavioral Reviews 1992, 16 (4), 525–534. https://doi.org/10.1016/s0149-7634(05)80194-0.

68. Grønli, J.; Murison, R.; Bjorvatn, B.; Sørensen, E.; Portas, C. M.; Ursin, R. Chronic Mild Stress Affects Sucrose Intake and Sleep in Rats. Behavioural Brain Research 2004, 150 (1-2), 139–147. https://doi.org/10.1016/s0166-4328(03)00252-3.

69. Baker, S. L.; Kentner, A. C.; Konkle, A. T. M.; Santa-Maria Barbagallo, L.; Bielajew, C. Behavioral and Physiological Effects of Chronic Mild Stress in Female Rats. Physiology & Behavior 2006, 87 (2), 314–322. https://doi.org/10.1016/j.physbeh.2005.10.019.

70. Willner, P. Chronic Mild Stress (CMS) Revisited: Consistency and Behavioural-Neurobiological Concordance in the Effects of CMS. Neuropsychobiology 2005, 52 (2), 90–110. https://doi.org/10.1159/000087097.

71. Moreau, J.-L.; Scherschlicht, R.; Jenck, F.; Martin, J. R. Chronic Mild Stress-Induced Anhedonia Model of Depression. Behavioural Pharmacology 1995, 6 (7), 682-687. https://doi.org/10.1097/00008877-199511000-00003.

72. Willner, P.; Towell, A.; Sampson, D.; Sophokleous, S.; Muscat, R. Reduction of Sucrose Preference by Chronic Unpredictable Mild Stress, and Its Restoration by a Tricyclic Antidepressant. Psychopharmacology 1987, 93 (3), 358–364. https://doi.org/10.1007/BF00187257.

73. den Hartog, C. R.; Blandino, K. L.; Nash, M. L.; Sjogren, E. R.; Grampetro, M. A.; Moorman, D. E.; Vazey, E. M. Noradrenergic Tone Mediates Marble Burying Behavior after Chronic Stress and Ethanol. Psychopharmacology 2020, 237 (10), 3021–3031. https://doi.org/10.1007/s00213-020-05589-7.

74. Kedia, S.; Chattarji, S. Marble Burying as a Test of the Delayed Anxiogenic Effects of Acute Immobilisation Stress in Mice. Journal of Neuroscience Methods 2014, 233, 150–154. https://doi.org/10.1016/j.jneumeth.2014.06.012.

75. Son, H.; Yang, J. H.; Kim, H. J.; Lee, D. K. A Chronic Immobilization Stress Protocol for Inducing Depression-like Behavior in Mice. Journal of Visualized Experiments 2019, No. 147. https://doi.org/10.3791/59546.

76. Ngoupaye, G. T.; Yassi, F. B.; Bahane, D. A. N.; Bum, E. N. Combined Corticosterone Treatment and Chronic Restraint Stress Lead to Depression Associated with Early Cognitive Deficits in Mice. Metabolic Brain Disease 2017, 33 (2), 421–431. https://doi.org/10.1007/s11011-017-0148-4.

77. Hill, M. N.; Hellemans, K. G. C.; Verma, P.; Gorzalka, B. B.; Weinberg, J. Neurobiology of Chronic Mild Stress: Parallels to Major Depression. Neuroscience & Biobehavioral Reviews 2012, 36 (9), 2085–2117. https://doi.org/10.1016/j.neubiorev.2012.07.001.

78. Antoniuk, S.; Bijata, M.; Ponimaskin, E.; Wlodarczyk, J. Chronic Unpredictable Mild Stress for Modeling Depression in Rodents: Meta-Analysis of Model Reliability. Neuroscience & Biobehavioral Reviews 2019, 99, 101–116. https://doi.org/10.1016/j.neubiorev.2018.12.002.

79. Ross, S.; Bossis, A.; Guss, J.; Agin-Liebes, G.; Malone, T.; Cohen, B.; Mennenga, S. E.; Belser, A.; Kalliontzi, K.; Babb, J.; Su, Z.; Corby, P.; Schmidt, B. L. Rapid and Sustained Symptom Reduction Following Psilocybin Treatment for Anxiety and Depression in Patients with Life-Threatening Cancer: A Randomized Controlled Trial. Journal of Psychopharmacology 2016, 30 (12), 1165–1180. https://doi.org/10.1177/0269881116675512.

80. Grob, C. S.; Danforth, A. L.; Chopra, G. S.; Hagerty, M.; McKay, C. R.; Halberstadt, A. L.; Greer, G. R. Pilot Study of Psilocybin Treatment for Anxiety in Patients with Advanced-Stage Cancer. Archives of General Psychiatry 2011, 68 (1), 71. https://doi.org/10.1001/archgenpsychiatry.2010.116.

81. Hesselgrave, N.; Troppoli, T. A.; Wulff, A. B.; Cole, A. B.; Thompson, S. M. Harnessing Psilocybin: Antidepressant-like Behavioral and Synaptic Actions of Psilocybin Are Independent of 5-HT2R Activation in Mice. Proceedings of the National Academy of Sciences 2021, 118 (17), e2022489118. https://doi.org/10.1073/pnas.2022489118.

82. Matsushima, Y.; Shirota, O.; Kikura-Hanajiri, R.; Goda, Y.; Eguchi, F. Effects Of Psilocybe Argentipeson Marble-Burying Behavior in Mice. Bioscience, Biotechnology, and Biochemistry 2009, 73 (8), 1866–1868. https://doi.org/10.1271/bbb.90095.

83. Gaynes, B. N. Identifying Difficult-To-Treat Depression. The Journal of Clinical Psychiatry 2009, 70 (suppl 6), 10–15. https://doi.org/10.4088/jcp.8133su1c.02.

84. McCabe, C.; Mishor, Z.; Cowen, P. J.; Harmer, C. J. Diminished Neural Processing of Aversive and Rewarding Stimuli during Selective Serotonin Reuptake Inhibitor Treatment. Biological Psychiatry 2010, 67 (5), 439–445. https://doi.org/10.1016/j.biopsych.2009.11.001.

85. Rautio, J.; Kumpulainen, H.; Heimbach, T.; Oliyai, R.; Oh, D.; Järvinen, T.; Savolainen, J. Prodrugs: Design and Clinical Applications. Nature Reviews Drug Discovery 2008, 7 (3), 255–270. https://doi.org/10.1038/nrd2468.

86. Shaba, R. Development of an Improved Psilocybin Synthesis; Uppsala University Innovation, Uppsala University, 2020.

87. Lenz, C.; Wick, J.; Braga, D.; García-Altares, M.; Lackner, G.; Hertweck, C.; Gressler, M.; Hoffmeister, D. Injury-Triggered Blueing Reactions of Psilocybe “Magic” Mushrooms. Angewandte Chemie International Edition 2020, 59 (4), 1450–1454. https://doi.org/10.1002/anie.201910175.

88. Peden, N. R.; Pringle, S. D.; Crooks, J. The Problem of Psilocybin Mushroom Abuse. Human Toxicology 1982, 1 (4), 417–424. https://doi.org/10.1177/096032718200100408.

89. Mawe, G. M.; Hoffman, J. M. Serotonin Signalling in the Gut—Functions, Dysfunctions and Therapeutic Targets. Nature Reviews Gastroenterology & Hepatology 2013, 10 (8), 473–486. https://doi.org/10.1038/nrgastro.2013.105.

