## Supplementary Information for "Novel psilocin prodrugs with altered pharmacological properties as candidate therapies for treatment-resistant anxiety disorders"

#### Supplementary Tables and Figures

**Table S1:** Pharmacokinetic (PK) analysis of maximum plasma psilocin concentration ($C_{max}$), measurable psilocin exposure ($AUC_{(0\text{-}tlast)}$) and apparent half-life ($T_{1/2}$) of systemic psilocin levels are listed. PK was evaluated in healthy C57BL/6 mice dosed with a single administration of psilocybin or an NDP dosed at 1 mg/kg (i.v.), and 1, 3, and 10 mg/kg (p.o.). LC-MS-based quantification of resultant psilocin metabolite was performed at timepoints 0.25, 0.5, 1, 2, 4, 6, 8 and 24 h post dose. An additional serial sample was collected at 5 min for animals dosed i.v. n = 3 for all dose groups; values presented as ± standard error.

| Prodrug* | Psilocin Cmax [ng/mL] | | | | Psilocin AUC (0-tlast) (h*ng/mL) | | | | Psilocin Apparent T1/2 (h) | | | |
|---|---|---|---|---|---|---|---|---|---|---|---|---|
| | 1 mg/kg (iv) | 1 mg/kg (po) | 3 mg/kg (po) | 10 mg/kg (po) | 1 mg/kg (iv) | 1 mg/kg (po) | 3 mg/kg (po) | 10 mg/kg (po) | 1 mg/kg (iv) | 1 mg/kg (po) | 3 mg/kg (po) | 10 mg/kg (po) |
| PCB | 104.8 ±11.4 | 53.7 ±2.02 | 52.9 ±6.03 | 243.0 ±25.2 | 89.7 ±10.0 | 67.1 ±0.93 | 84.4 ±6.24 | 397.3 ±31.9 | 0.91 ±n/a | 2.15 ±n/a | 4.52 ±2.07 | 3.58 ±0.53 |
| ES01 | 129.5 ±43.8 | 9.15 ±0.55 | 28.5 ±2.86 | 78.9 ±13.9 | 37.1 ±5.83 | 10.9 ±1.21 | 34.8 ±1.47 | 112.3 ±8.65 | 1.34 ±0.28 | 1.55 ±0.01 | 2.32 ±0.48 | 1.52 ±n/a |
| ES02 | 64.1 ±42.0 | 8.34 ±1.48 | 13.0 ±1.77 | 52.8 ±3.2 | 21.2 ±11.4 | 7.19 ±1.31 | 13.4 ±0.42 | 53.3 ±5.71 | 1.42 ±0.15 | 1.47 ±0.37 | 1.44 ±0.13 | 1.82 ±0.17 |
| ES03 | 32.7 ±3.01 | 9.34 ±0.82 | 31.4 ±2.14 | 75.6 ±15.1 | 19.8 ±1.45 | 11.9 ±0.51 | 40.4 ±2.13 | 106.7 ±11.7 | 1.46 ±0.16 | 1.90 ±0.63 | 1.44 ±0.02 | 1.20 ±0.04 |
| ES04 | 26.7 ±1.76 | 15.0 ±2.73 | 29.3 ±3.09 | 126.1 ±36.5 | 30.9 ±0.64 | 23.5 ±3.70 | 44.8 ±2.26 | 170.7 ±25.1 | 0.99 ±0.07 | 1.74 ±0.06 | 2.38 ±0.92 | 3.68 ±0.81 |
| ES05 | 22.3 ±14.4 | 2.68 ±0.07 | 12.6 ±2.59 | 46.5 ±4.31 | 23.2 ±12.4 | 6.38 ±0.70 | 19.0 ±1.38 | 67.2 ±5.56 | 1.23 ±0.20 | 2.09 ±0.25 | 1.60 ±0.08 | 2.15 ±0.66 |
| C01 | 0.58 ±0.04 | 0.14 ±0.01 | 0.33 ±0.03 | 0.87 ±0.14 | 0.32 ±0.04 | n/d | 0.40±0.02 | 1.19 ±0.10 | 0.62 ±0.14 | n/d | n/d | 1.23 ±0.35 |
| C02 | 79.6 ±1.54 | 19.7 ±3.18 | 55.6 ±5.01 | 306 ±20.0 | 67.0 ±1.37 | 21.0 ±0.88 | 60.8 ±4.84 | 282.3 ±24.0 | 1.95 ±0.18 | 1.69 ±0.06 | 1.23 ±0.12 | 1.83 ±0.31 |
| C03 | 45.9 ±1.29 | 17.3 ±4.32 | 31.6 ±1.42 | 93.5 ±7.59 | 28.2 ±0.55 | 16.9 ±0.69 | 44.2 ±2.34 | 182.7 ±12.3 | 1.34 ±0.48 | 3.45 ±0.11 | 2.33 ±0.60 | 3.99 ±0.46 |
| T01 | 553.8 ±256.9 | 6.74 ±0.92 | 25.5 ±2.52 | 80.1 ±16.5 | 169.0 ±69.6 | 10.7 ±0.25 | 31.4 ±0.95 | 123.7 ±1.20 | 1.33 ±0.48 | 1.66 ±0.33 | 2.88 ±0.74 | 1.39 ±n/a |
| EE01 | 23.9 ±14.5 | 3.65 ±0.59 | 14.3 ±0.66 | 44.0 ±0.98 | 9.34 ±3.88 | 4.75 ±0.49 | 12.0 ±0.30 | 48.7 ±3.37 | 0.80 ±0.35 | 0.84 ±0.04 | 1.17 ±0.05 | 2.59 ±0.12 |
| EE02 | 84.5 ±70.3 | 4.55 ±0.23 | 12.1 ±1.17 | 35.6 ±2.19 | 28.3 ±22.3 | 4.32 ±0.28 | 11.5 ±0.73 | 42.8 ±2.51 | 1.01 ±0.13 | 0.75 ±0.27 | 0.87 ±0.06 | 1.65 ±0.02 |
| EC01 | 2.12 ±0.12 | 0.90 ±0.07 | 2.78 ±0.47 | 7.9 ±1.77 | n/d | n/d | 3.09 ±0.37 | 10.0 ±2.15 | n/d | n/d | n/d | n/d |
| SE01 | 2.78 ±0.15 | 3.94 ±0.25 | 10.8 ±0.73 | 24.3 ±1.43 | 7.67 ±0.05 | 8.13 ±0.59 | 24.3 ±0.70 | 66.6 ±2.65 | 2.78 ±0.88 | 1.51 ±0.40 | 1.78 ±0.31 | 3.04 ±1.13 |
| SE02 | 1.22 ±012 | 0.53 ±0.09 | 1.18 ±0.38 | 5.6 ±0.63 | 1.44 ±0.12 | n/d | 3.31 ±0.06 | 12.5 ±0.72 | 0.98 ±n/a | n/d | 1.21 ±n/a | 2.33 ±0.26 |
| P02 | 2.18 ±0.10 | 0.69 ±0.04 | 1.77 ±0.10 | 6.1 ±0.14 | 1.43 ±0.07 | 1.07 ±0.17 | 2.58 ±0.24 | 10.3 ±0.52 | 0.96 ±0.31 | 3.38 ±2.1 | 1.69 ±0.28 | 1.85 ±0.24 |

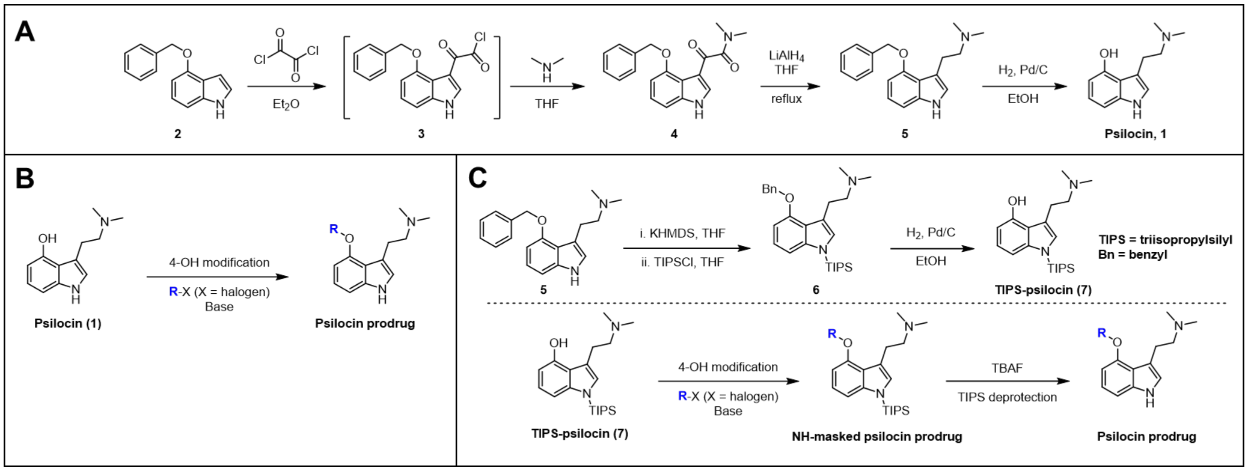


**Figure S1.** A) Synthesis of psilocin (**1**) from 4-benzyloxy-indole (**2**). B) General scheme for the synthesis of 4-OH-derived prodrugs. C) General scheme for the synthesis of 4-OH-derived prodrugs with masked indole-NH to achieve orthogonal derivatization. See below for reagents, detailed synthetic procedures, and reaction yields.

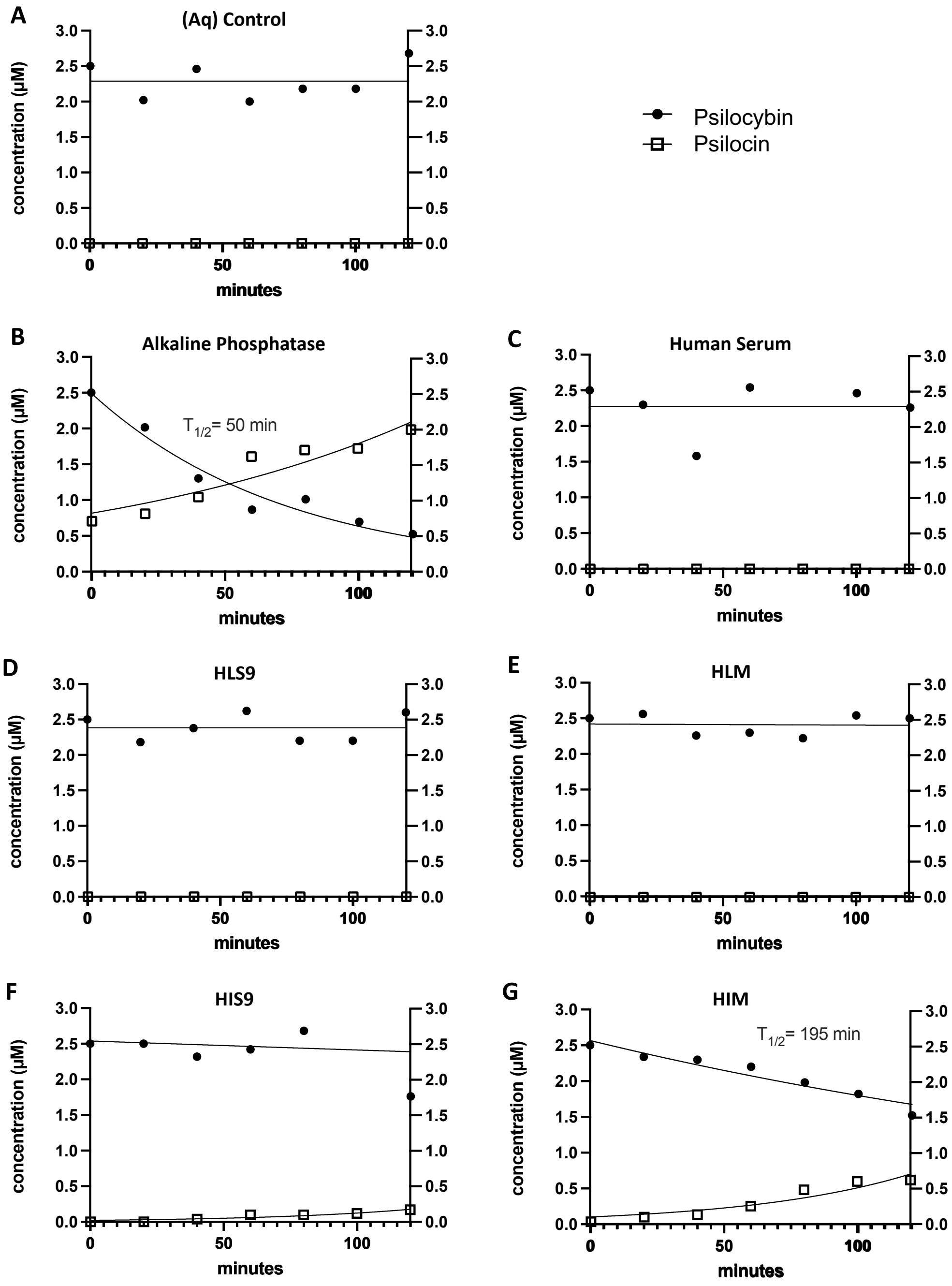


**Figure S2.** *In vitro* metabolism of psilocybin incubated in assay buffer (Aq), bovine alkaline phosphatase, human serum, and human S9 and microsomal (M) fractions derived from donor liver (HL) and intestinal (HI) tissues. Decay of parent psilocybin and biotransformation into psilocin metabolite over 120-min incubation period is plotted in µM. Psilocybin was prepared at an assay concentration of 2.5 µM and incubated in each biological fraction independently for a total of 120 min. LC-MS-based quantification of resultant psilocin metabolite was performed at timepoints 0, 20, 40, 60, 80, 100 and 120 min.

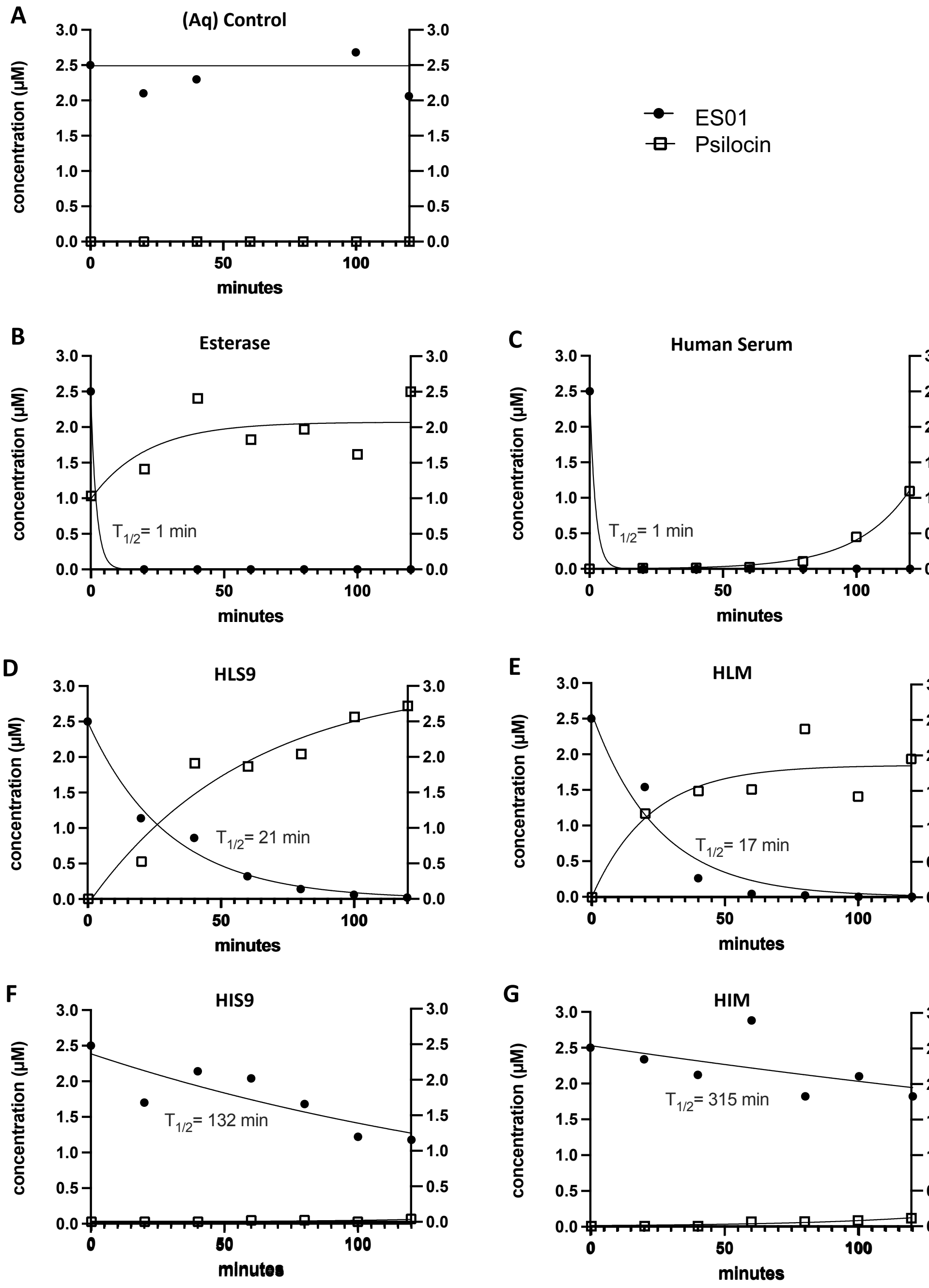


**Figure S3.** *In vitro* metabolism of ES01 incubated in assay buffer (Aq), porcine esterase, human serum, and human S9 and microsomal (M) fractions derived from donor liver (HL) and intestinal (HI) tissues. Decay of parent ES01 and biotransformation into psilocin metabolite over 120-min incubation period is plotted in µM. ES01 was prepared at an assay concentration of 2.5 µM and incubated in each biological fraction independently for a total of 120 min. LC-MS-based quantification of resultant psilocin metabolite was performed at timepoints 0, 20, 40, 60, 80, 100 and 120 min.

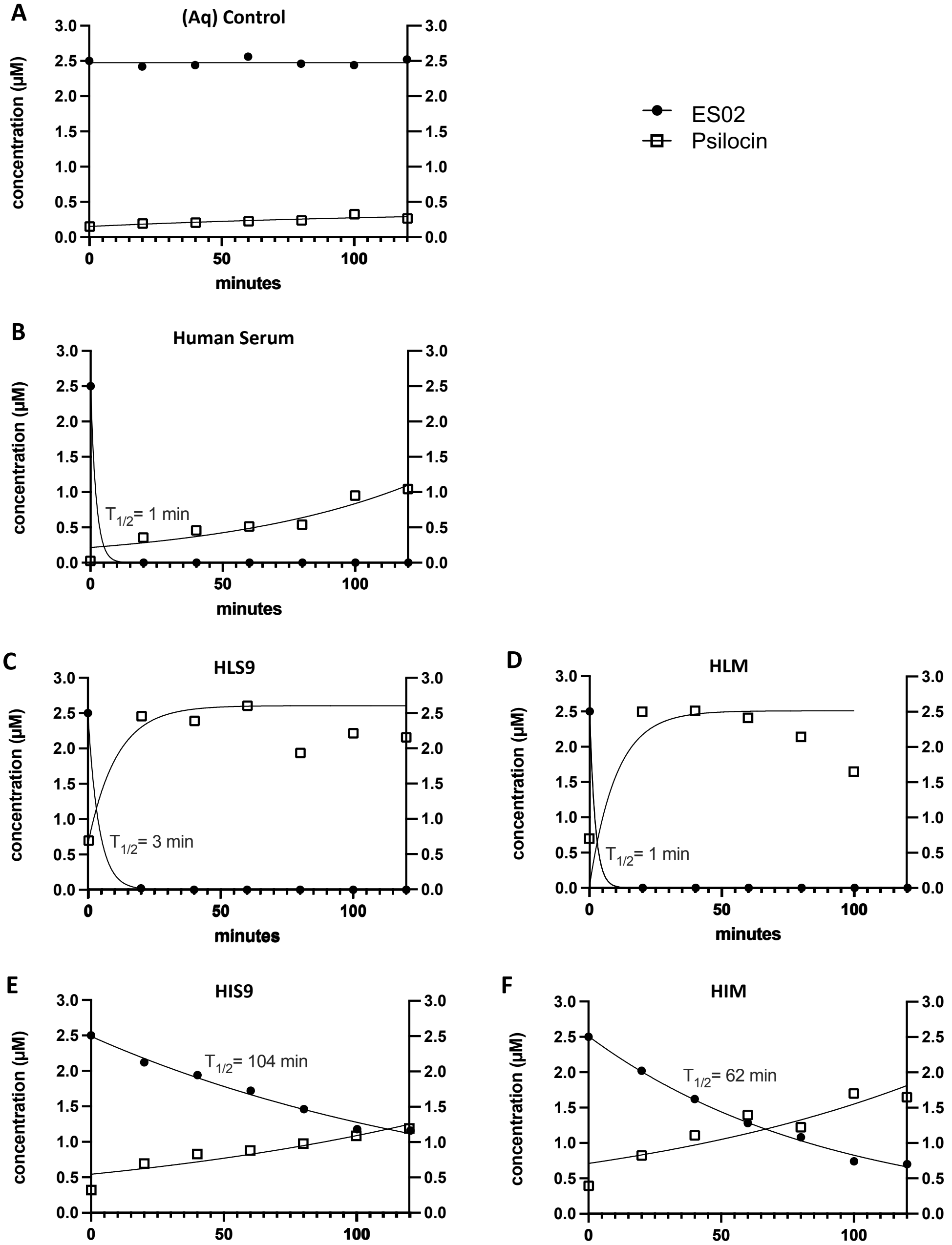


**Figure S4.** *In vitro* metabolism of ES02 incubated in assay buffer (Aq), human serum, and human S9 and microsomal (M) fractions derived from donor liver (HL) and intestinal (HI) tissues. Decay of parent ES02 and biotransformation into psilocin metabolite over 120-min incubation period is plotted in μM. ES02 was prepared at an assay concentration of 2.5 μM and incubated in each biological fraction independently for a total of 120 min. LC-MS-based quantification of resultant psilocin metabolite was performed at timepoints 0, 20, 40, 60, 80, 100 and 120 min.

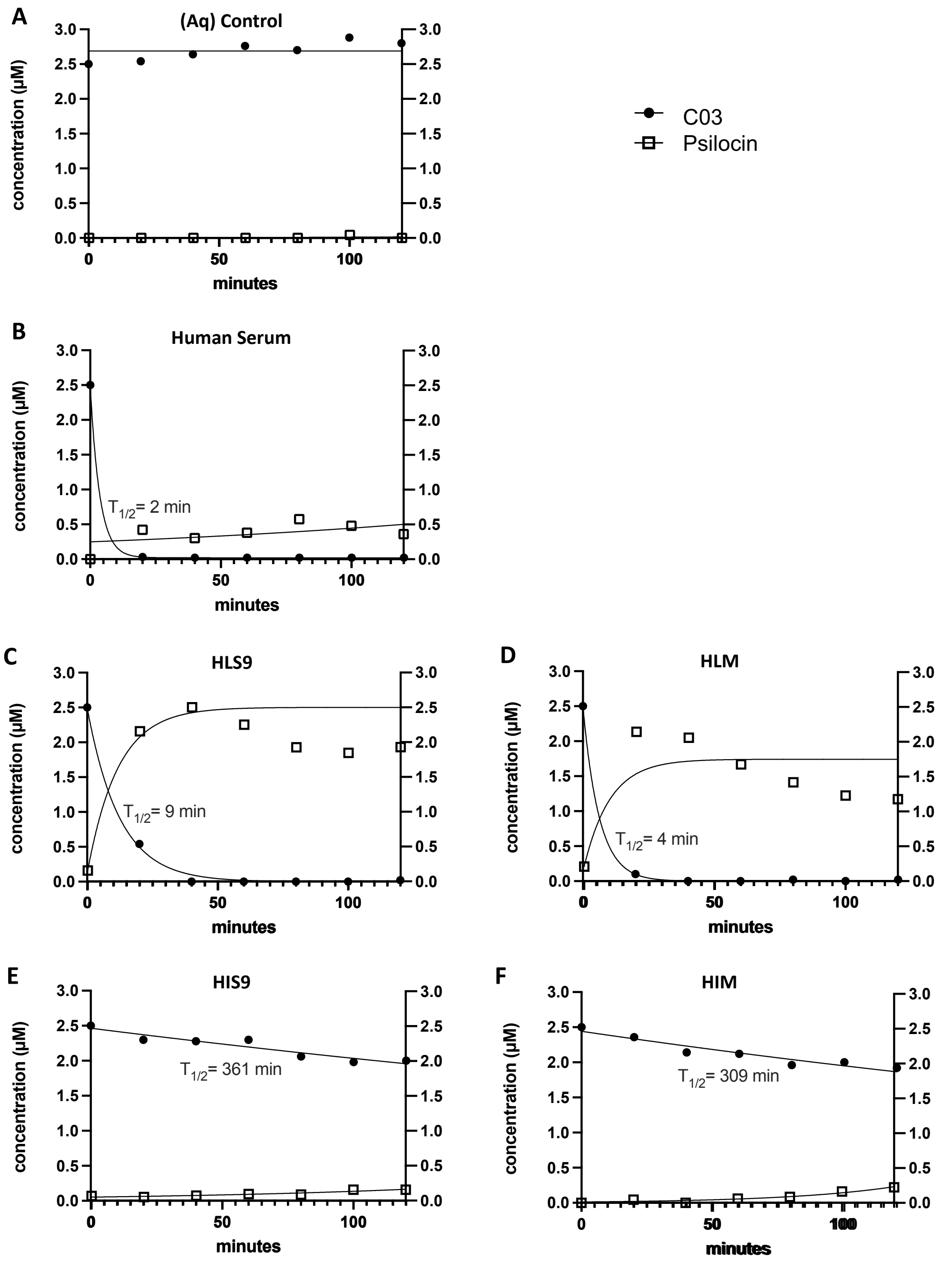


**Figure S5.** *In vitro* metabolism of C03 incubated in assay buffer (Aq), human serum, and human S9 and microsomal (M) fractions derived from donor liver (HL) and intestinal (HI) tissues. Decay of parent C03 and biotransformation into psilocin metabolite over 120-min incubation period is plotted in µM. C03 was prepared at an assay concentration of 2.5 µM and incubated in each biological fraction independently for a total of 120 min. LC-MS-based quantification of resultant psilocin metabolite was performed at timepoints 0, 20, 40, 60, 80, 100 and 120 min.

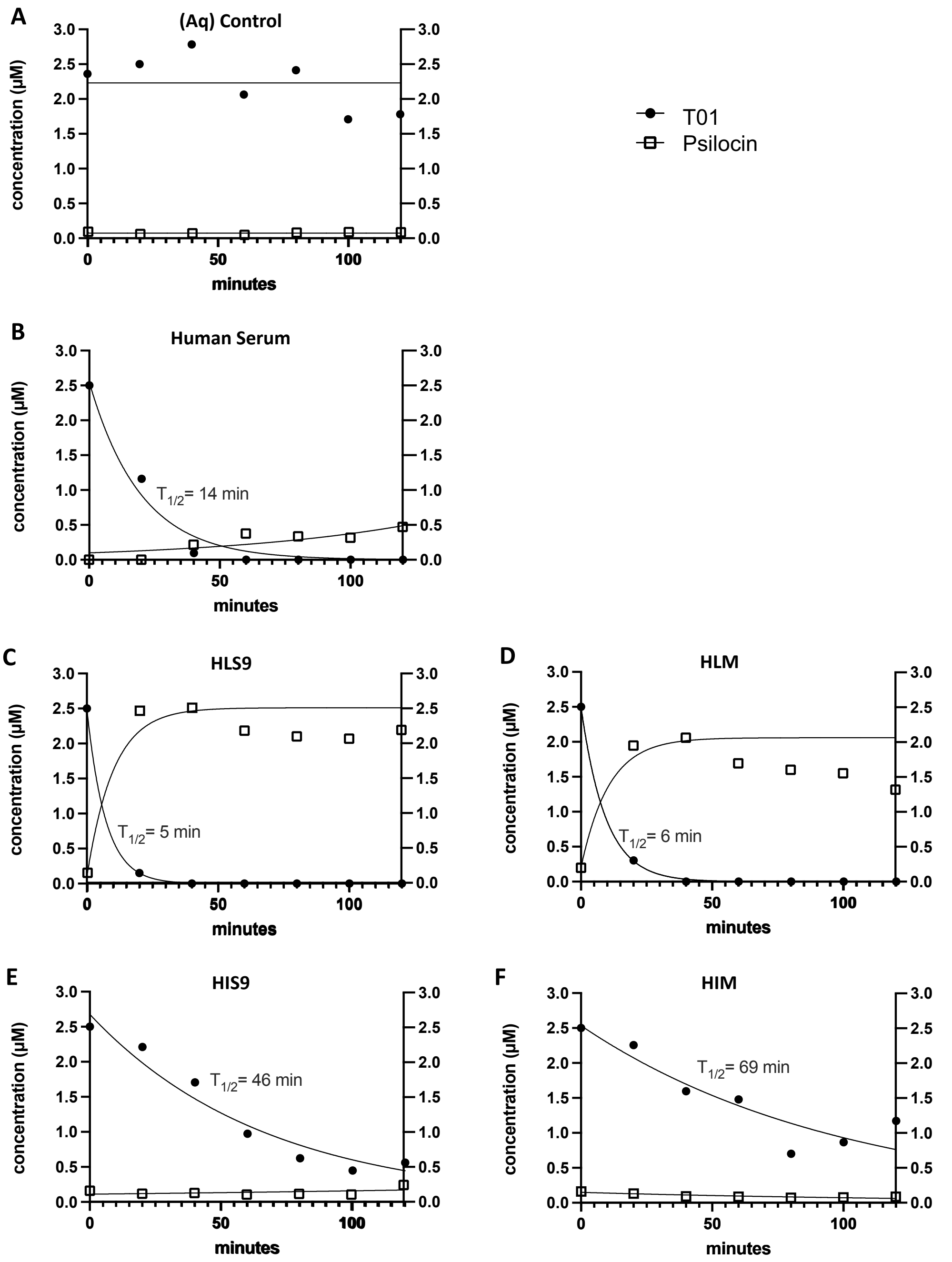


**Figure S6.** *In vitro* metabolism of T01 incubated in assay buffer (Aq), human serum, and human S9 and microsomal (M) fractions derived from donor liver (HL) and intestinal (HI) tissues. Decay of parent T01 and biotransformation into psilocin metabolite over 120-min incubation period is plotted in µM. T01 was prepared at an assay concentration of 2.5 µM and incubated in each biological fraction independently for a total of 120 min. LC-MS-based quantification of resultant psilocin metabolite was performed at timepoints 0, 20, 40, 60, 80, 100 and 120 min.

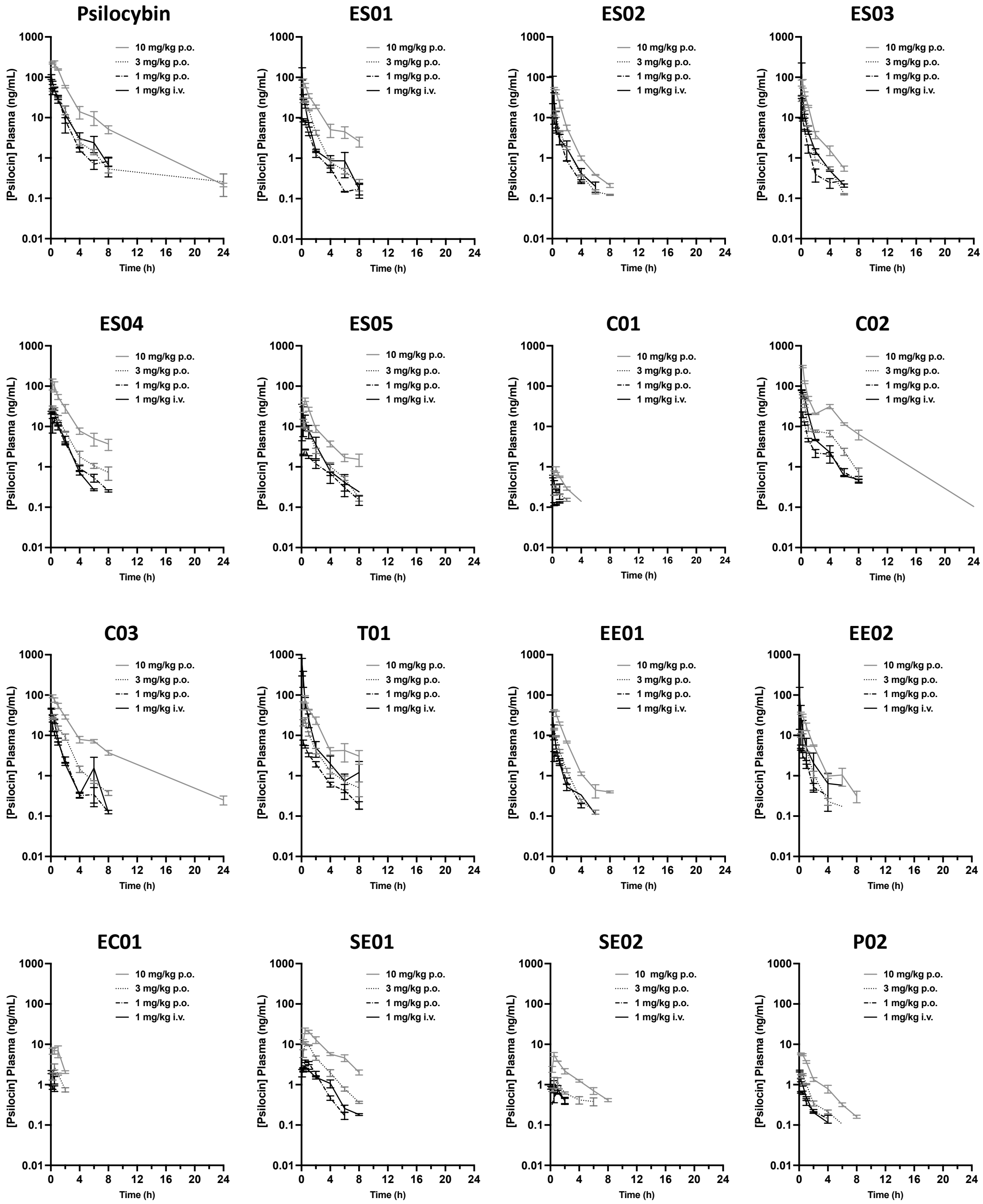


**Figure S7.** Pharmacokinetic (PK) analysis of plasma psilocin concentration. Plasma PK was evaluated in healthy C57BL/6 mice dosed with a single administration of psilocybin or an NDP dosed at 1 mg/kg (i.v.), and 1, 3, and 10 mg/kg (p.o.). LC-MS-based quantification of resultant psilocin metabolite was performed at timepoints 0.25, 0.5, 1, 2, 4, 6, 8 and 24 h post dose. An additional serial sample was collected at 0.0833 hours for animals dosed i.v. n = 3 for all dose groups. Values plotted as mean ng/mL plasma concentration ± standard error.

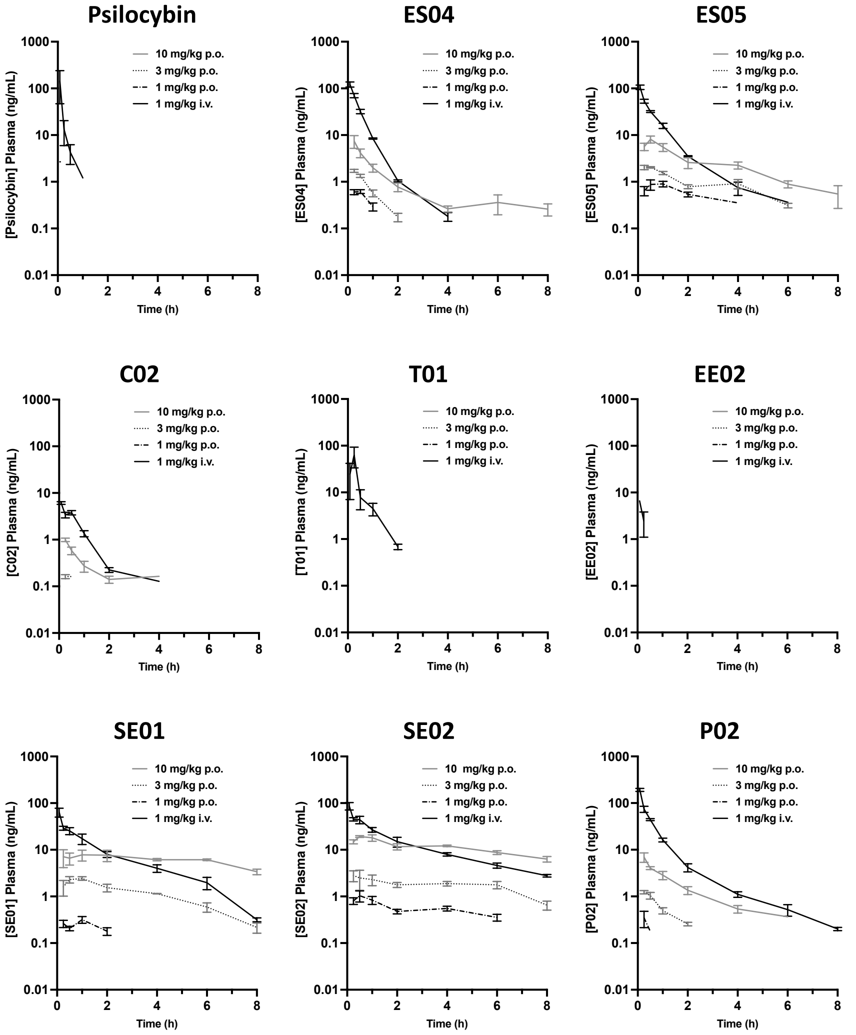


**Figure S8.** Pharmacokinetic (PK) analysis of plasma prodrug concentration. Plasma PK was evaluated in healthy C57BL/6 mice dosed with a single administration of psilocybin or an NDP dosed at 1 mg/kg (i.v.), and 1, 3, and 10 mg/kg (p.o.). LC-MS-based quantification of residual parent prodrug analyte was performed at timepoints 0.25, 0.5, 1, 2, 4, 6, 8 and 24 hours post dose. An additional serial sample was collected at 0.0833 hours for animals dosed i.v. n = 3 for all dose groups. Values plotted as mean ng/mL prodrug concentration ± standard error. Note, only those NPDs with detectable plasma parent prodrug levels are presented.

### Supplementary Dataset 1

#### Synthesis of Psilocin

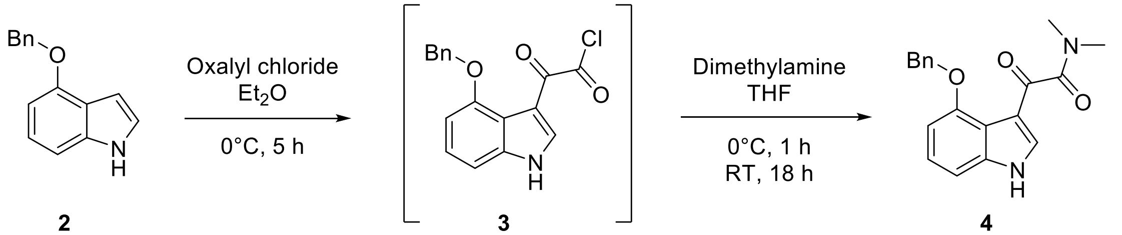


2 3 4

**2-(4-(Benzyloxy)-1*H*-indol-3-yl)-*N*,*N*-dimethyl-2-oxoacetamide (4)**

A dry, 3-neck RBF was charged with 4-benzyloxyindole **2** (14.0 g, 62.7 mmol) and $Et_2O$ (327 mL) under Ar. The mixture was cooled down to 0 °C in an ice bath. An argon sparge was placed on the RBF and into the reaction mixture to purge out the HCl gas released from the reaction. Oxalyl chloride (10.9 mL, 129 mmol) was added dropwise over 40 min, while maintaining the cold temperature. The mixture was stirred at 0°C for 4 h. The argon sparge was removed, and dimethylamine (157 mL, 314 mmol) (2 M in THF) was added dropwise at 0 °C over 1 h using an addition funnel. The mixture was allowed to warm up to RT and stir overnight. Diethyl ether (200 mL) was added, and the mixture was cooled down to 0 °C. The resulting precipitate (crude 3) was filtered and transferred to an Erlenmeyer flask. The solid was suspended in water (300 mL) and stirred for 30 min. Then, it was filtered and washed with more $H_2O$ to remove residual salts. The crude solid was further dried *in vacuo* to afford **4** which was used in the next step without further purification.

LRMS-HESI: calculated: 323.14; observed: 323.18 m/z $[M+H]^+$

$^1H$ NMR (400 MHz, $CDCl_3$) δ 10.44 (s, 1H), 7.54 (ddt, J = 7.4, 1.3, 0.7 Hz, 2H), 7.50 (d, J = 3.2 Hz, 1H), 7.42 – 7.35 (m, 2H), 7.34 – 7.29 (m, 1H), 7.03 (t, J = 8.0 Hz, 1H), 6.89 (dd, J = 8.2, 0.8 Hz, 1H), 6.64 (dd, J = 7.9, 0.8 Hz, 1H), 5.25 (s, 2H), 2.96 (s, 3H), 2.91 (s, 3H).

$^{1}$H-NMR of **4**, 400 MHz, $CDCl_3$

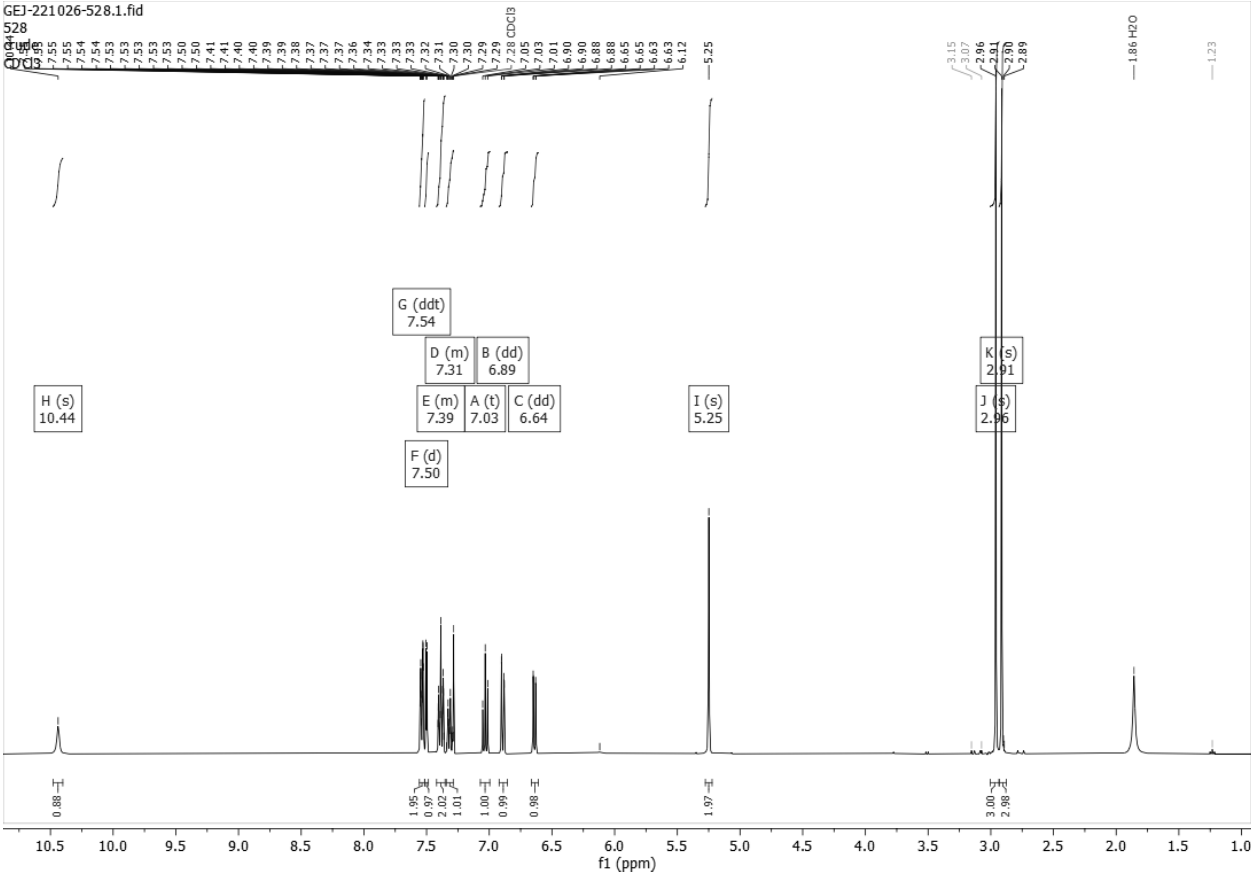

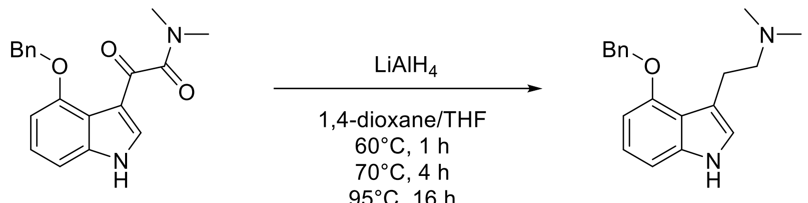

4 5

**2-(4-(Benzyloxy)-1*H*-indol-3-yl)-*N*,*N*-dimethylethanamine, 4-BnO-DMT (5)**

Lithium aluminum hydride (2M in THF, 60.2 mL, 120 mmol) was added to a dry 3-neck flask under argon. The flask was fitted with a reflux condenser and an addition funnel. Dry 1,4-dioxane (100 mL) was added, and the mixture was heated to 60 °C in an oil bath. In a separate flask, compound **4** (7.46 g, 23.1 mmol) was dissolved in a mixture of THF (60 mL) and 1,4-dioxane (120 mL). This solution was added in a dropwise-manner to the rapidly stirring $LiAlH_4$ solution over 1 h using an addition funnel. The oil bath temperature was held at 70 °C for 4 h, followed by vigorous reflux overnight (16 h) in an oil bath temperature of 95 °C. The reaction was placed in an ice bath, and a solution of distilled $H_2O$ (25 mL) in THF (65 mL) was added dropwise to quench $LiAlH_4$, resulting in a gray flocculent precipitate. $Et_2O$ (160 mL) was added to assist breakup of the complex and improve filtration. This slurry was stirred for 1 h and the mixture was then filtered using a Buchner funnel. The filter cake was washed on the filter with warm $Et_2O$ (2 x 200 mL) and was broken up, transferred back into the reaction flask and vigorously stirred with additional warm $Et_2O$ (300 mL). This slurry was filtered, and the cake was washed on the filter with $Et_2O$ (120 mL) and hexane (2 x 120 mL). All the organic filtrates were combined and dried ($MgSO_4$). After the drying agent was removed by filtration, the filtrate was concentrated under vacuum and dried under high vacuum. The crude residue was triturated with EtOAc/hex (1:9, 25 mL) to afford the crude product (**5**) which was used in the next step without further purification.

LRMS-HESI: calculated: 295.18; observed: 295.19 m/z $[M+H]^+$

$^1H$ NMR (400 MHz, $CDCl_3$) δ 8.04 (s, 1H), 7.52 (dtd, J = 7.2, 1.4, 0.6 Hz, 2H), 7.39 (tt, J = 6.5, 1.0 Hz, 2H), 7.35 – 7.31 (m, 1H), 7.11 – 7.02 (m, 1H), 6.97 (dd, J = 8.2, 0.8 Hz, 1H), 6.90 (dd, J = 2.0, 1.0 Hz, 1H), 6.56 (dd, J = 7.7, 0.8 Hz, 1H), 5.21 (s, 2H), 3.11 – 3.02 (m, 2H), 2.65 – 2.56 (m, 2H), 2.16 (s, 6H).

$^1$H-NMR of **5**, 400 MHz, $CDCl_3$

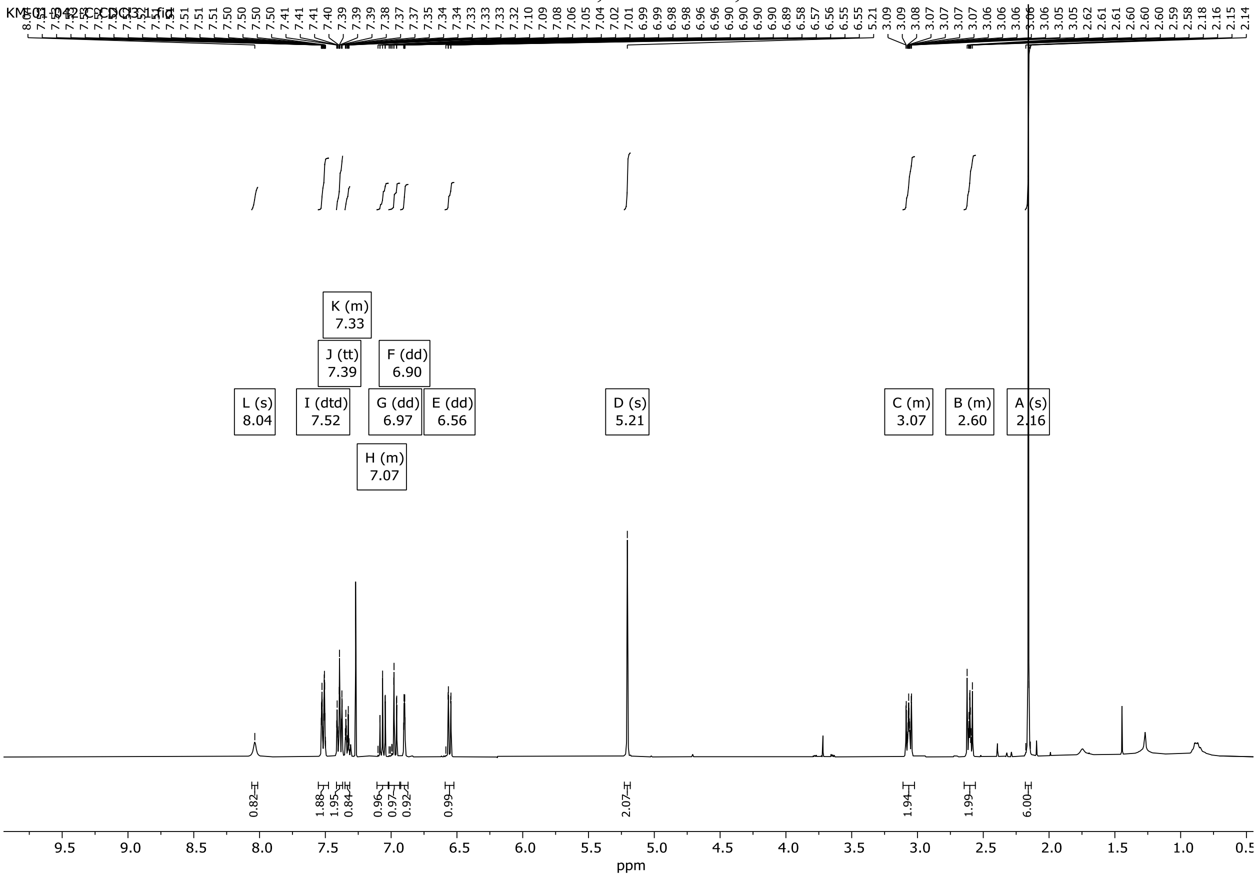

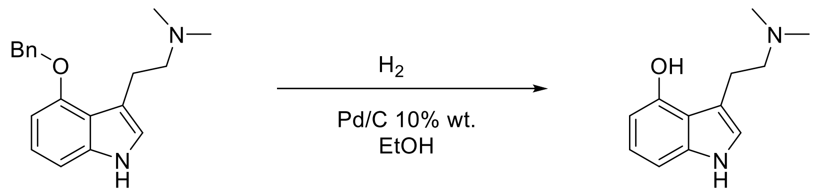


**5** **Psilocin (1)**

**3-(2-(Dimethylamino)ethyl)-1*H*-indol-4-ol, Psilocin (1)**

To a stirring solution of **5** (2.00 g, 6.79 mmol) in ethanol (106 mL) under argon was added palladium on carbon (10 wt%, 723 mg, 0.679 µmol). The vessel was evacuated and backfilled with $H_2$ gas three times, and then a balloon of $H_2$ was left attached to the vessel. The reaction mixture was allowed to stir at room temperature for 75 minutes. Anhydrous magnesium sulphate was added to remove water, and the mixture was filtered over Celite. The filtrate was concentrated under reduced pressure to yield a brown solid. Purification by column chromatography on 25 g silica using a 0-18% methanol-dichloromethane gradient yielded the pure product **1** (psilocin) as a tan solid (530 mg, 38%). Alternatively, the crude product may be used in the next step without further purification (80-85% purity).

LRMS-HESI: calculated: 205.13; observed: 205.15 m/z $[M+H]^+$

$^1$H NMR (400 MHz, MeOD) δ 6.93 – 6.81 (m, 3H), 6.36 (dd, J = 7.3, 1.1 Hz, 1H), 3.12 – 3.03 (m, 2H), 2.88 (t, J = 6.9 Hz, 2H), 2.46 (s, 6H).

$^1$H-NMR of **Psilocin (1)**, 400 MHz, MeOD

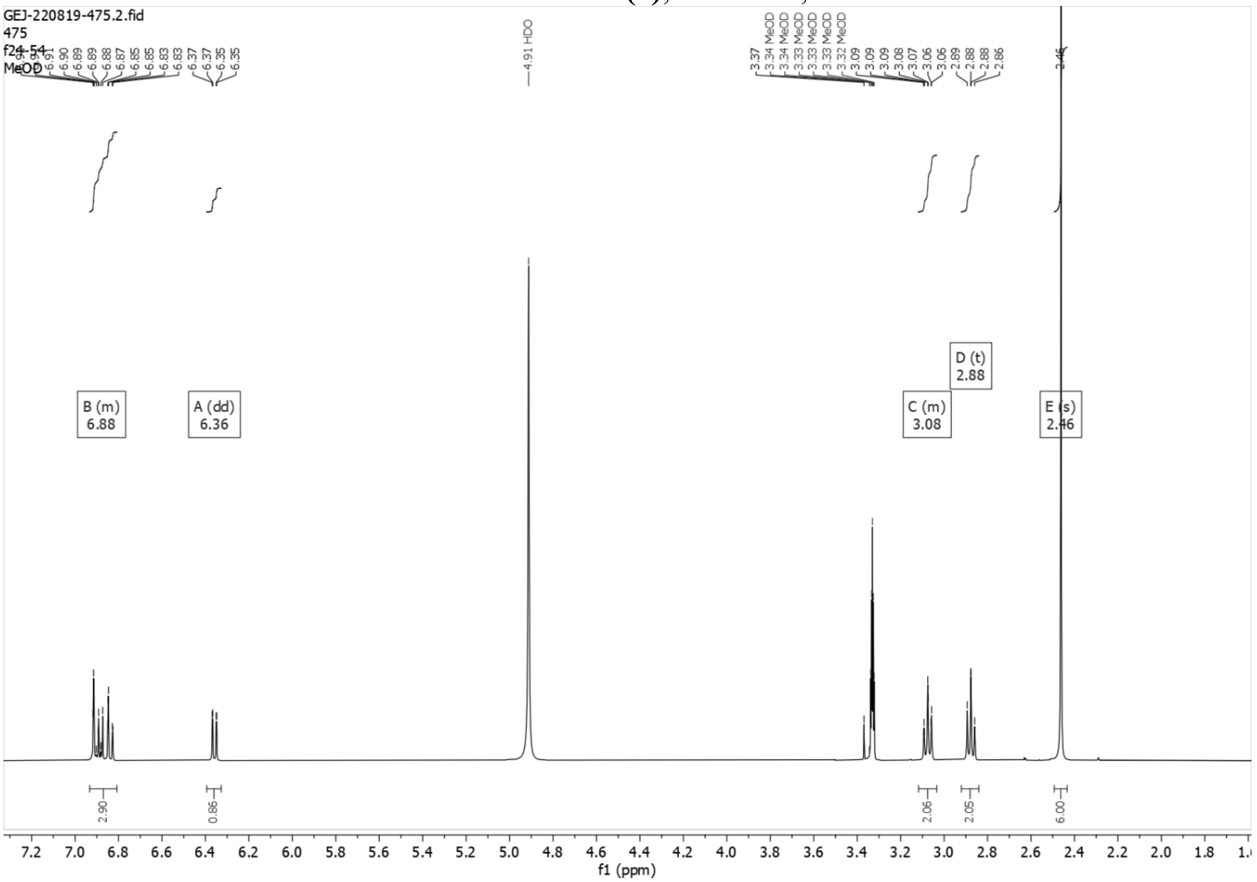

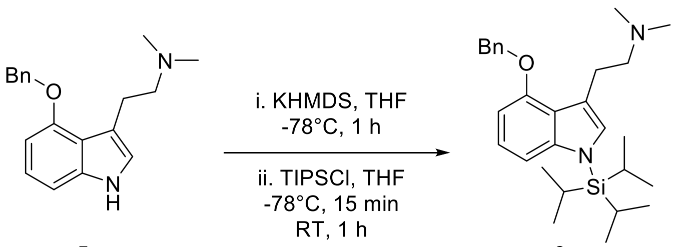


**5** **6**

**2-(4-(Benzyloxy)-1-(triisopropylsilyl)-1*H*-indol-3-yl)-*N*,*N*-dimethylethanamine, 4-BnO-TIPS-DMT (6)**

To a solution of **5** (5.00 g, 17.0 mmol) in dry THF (100 mL) cooled to -78 °C under argon was added dropwise a 1 M solution of KHMDS (18.7 mL, 18.7 mmol) in THF. After stirring at -78 °C for 1 h, a solution of TIPSCl (3.82 mL, 17.8 mmol) in THF (19.0 mL) was added dropwise over 15 minutes, and the reaction mixture was allowed to warm up to RT. After stirring at RT for 1 h, the reaction was quenched with $H_2O$ (40 mL), THF was evaporated under reduced pressure, and the aqueous solution was further diluted with $H_2O$ (75 mL) and extracted with DCM (3 x 100 mL). The organic layers were combined and washed with brine, dried over $Na_2SO_4$, filtered, and concentrated under reduced pressure. The crude product was purified by flash column chromatography (MeOH/DCM 5:95 to 10:90) to afford the pure product **6** as a light brown oil (6.99 g, 91%).

LRMS-HESI: calculated: 451.31; observed: 451.37 m/z $[M+H]^+$

$^1H$ NMR (400 MHz, $CDCl_3$) δ 7.58 – 7.51 (m, 2H), 7.44 – 7.39 (m, 2H), 7.38 – 7.33 (m, 1H), 7.12 (dd, J = 8.4, 0.8 Hz, 1H), 7.08 – 6.99 (m, 1H), 6.94 (s, 1H), 6.60 (dd, J = 7.7, 0.7 Hz, 1H), 5.20 (s, 2H), 3.12 – 3.04 (m, 2H), 2.67 – 2.58 (m, 2H), 2.16 (s, 6H), 1.69 (h, J = 7.5 Hz, 3H), 1.16 (d, J = 7.5 Hz, 18H).

[1]H-NMR of **6**, 400 MHz, $CDCl_3$

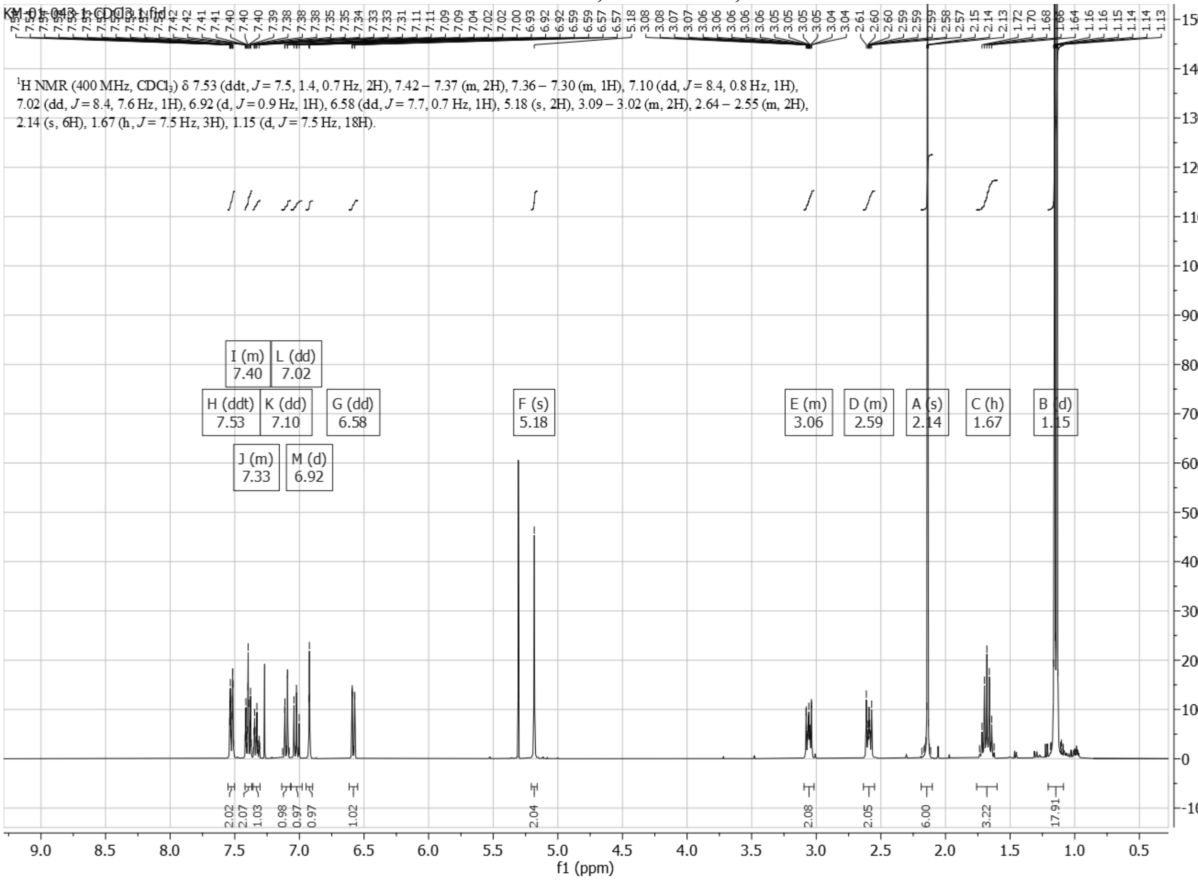

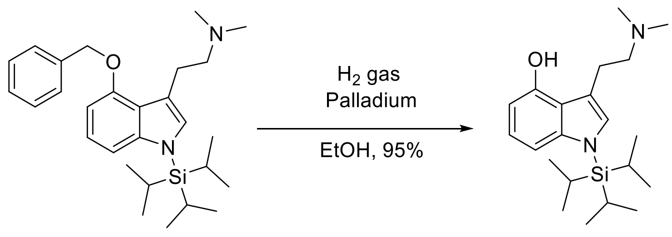


**6** **7**

**3-(2-(dimethylamino)ethyl)-1-(triisopropylsilyl)-1*H*-indol-4-ol, *N*-TIPS-Psilocin (7)**

To a stirring solution of 6 (6.99 g, 15.5 mmol) dissolved in EtOH, 95% (310 mL), was added palladium on carbon (10 wt%, 1.65 g, 1.55 mmol). This mixture was put under vacuum for five minutes, then alternately purged with $H_2$ gas until pressurized hydrogen atmosphere was established, then allowed to stir for 75 minutes at room temperature. The palladium on carbon was removed by filtration through Celite, the filtrate dried with anhydrous magnesium sulphate, and concentrated under reduced pressure to yield **7** (4.67 g, 84%) as an off-white solid.

HRMS-HESI: calculated: 361.2670; observed: 361. 2668 m/z $[M+H]^+$

$^1$H NMR (400 MHz, MeOD) δ 6.98 (d, J = 8.6 Hz, 2H), 6.91 (dd, J = 8.4, 7.5 Hz, 1H), 6.42 (dd, J = 7.5, 0.8 Hz, 1H), 3.06 (t, J = 6.9 Hz, 2H), 2.77 (t, J = 6.9 Hz, 2H), 2.39 (s, 6H), 1.72 (p, J = 7.5 Hz, 3H), 1.16 (d, J = 7.5 Hz, 18H).

$^1$H-NMR of **7**, 400 MHz, MeOD

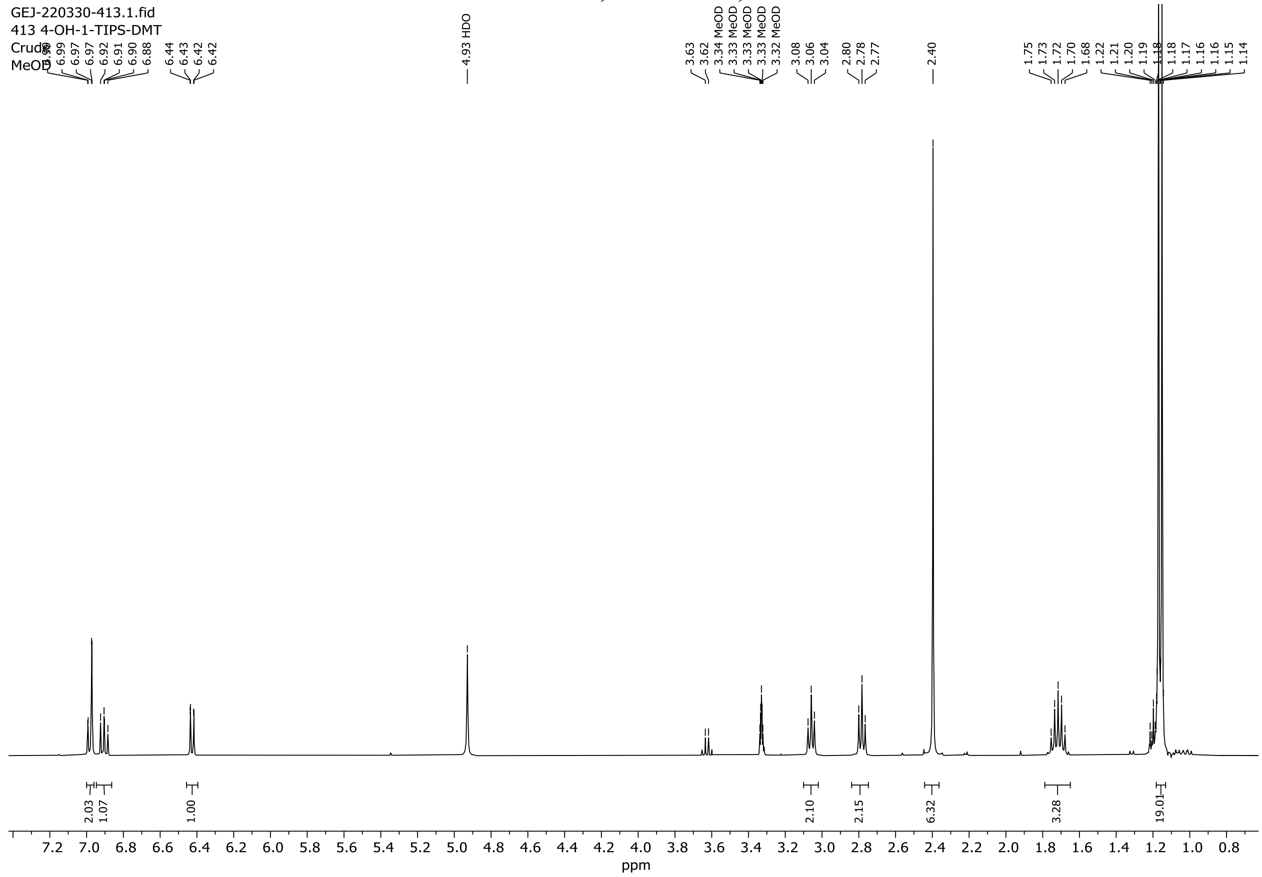

**Synthesis of psilocin prodrugs**

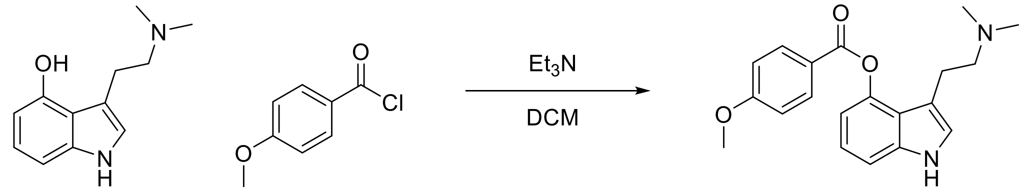


1 ES01

**3-(2-(Dimethylamino)ethyl)-1*H*-indol-4-yl 4-methoxybenzoate (ES01)**

A solution of psilocin **1** (200 mg, 979 µmol) and triethylamine (274 µL, 1.96 mmol) in DCM (8.0 mL) was cooled down to 0 °C. To it was added 4-methoxybenzoyl chloride (192 mg, 1.12 mmol) in DCM (0.5 mL). The resulting mixture was warmed up to RT and stirred for 2 h. At this time, another portion of 4-methoxybenzoyl chloride **2** (192 mg, 1.12 mmol) in DCM (0.5 mL) was added, and the reaction was stirred at RT for another hour. Methanol (2 mL) was added to the reaction, and the volatiles were removed in vacuo. The crude residue was directly purified by FC on silica gel (12 g, MeOH/DCM 0:100 to 20:80, 10 CV, product eluting at 11% MeOH) to afford the product as a brown oil. TLC showed co-elution of p-methoxybenzoic acid with the product. This isolated material was re-dissolved in DCM (50 mL), and extracted with sat'd aq. NaHCO3 (2 x 30 mL). The aq. layer was back-extracted with DCM (x1), washed with brine, dried over anhydrous $Na_2SO_4$, filtered and concentrated to afford the pure product (**ES01**) as an off-white foamy solid (245 mg, 74%).

LRMS-HESI: calculated: 339.17; observed: 339.16 m/z $[M+H]^+$

$^1$H NMR (400 MHz, $CDCl_3$) δ 8.27 – 8.21 (m, 3H), 7.23 (dd, J = 8.2, 1.0 Hz, 1H), 7.17 (t, J = 7.8 Hz, 1H), 7.03 – 6.98 (m, 2H), 6.95 (dd, J = 2.3, 1.1 Hz, 1H), 6.88 (dd, J = 7.6, 1.0 Hz, 1H), 3.90 (s, 3H), 2.92 – 2.84 (m, 2H), 2.60 – 2.52 (m, 2H), 2.09 (s, 6H).

### $^{1}$H-NMR of **ES01**, 400 MHz, $CDCl_3$

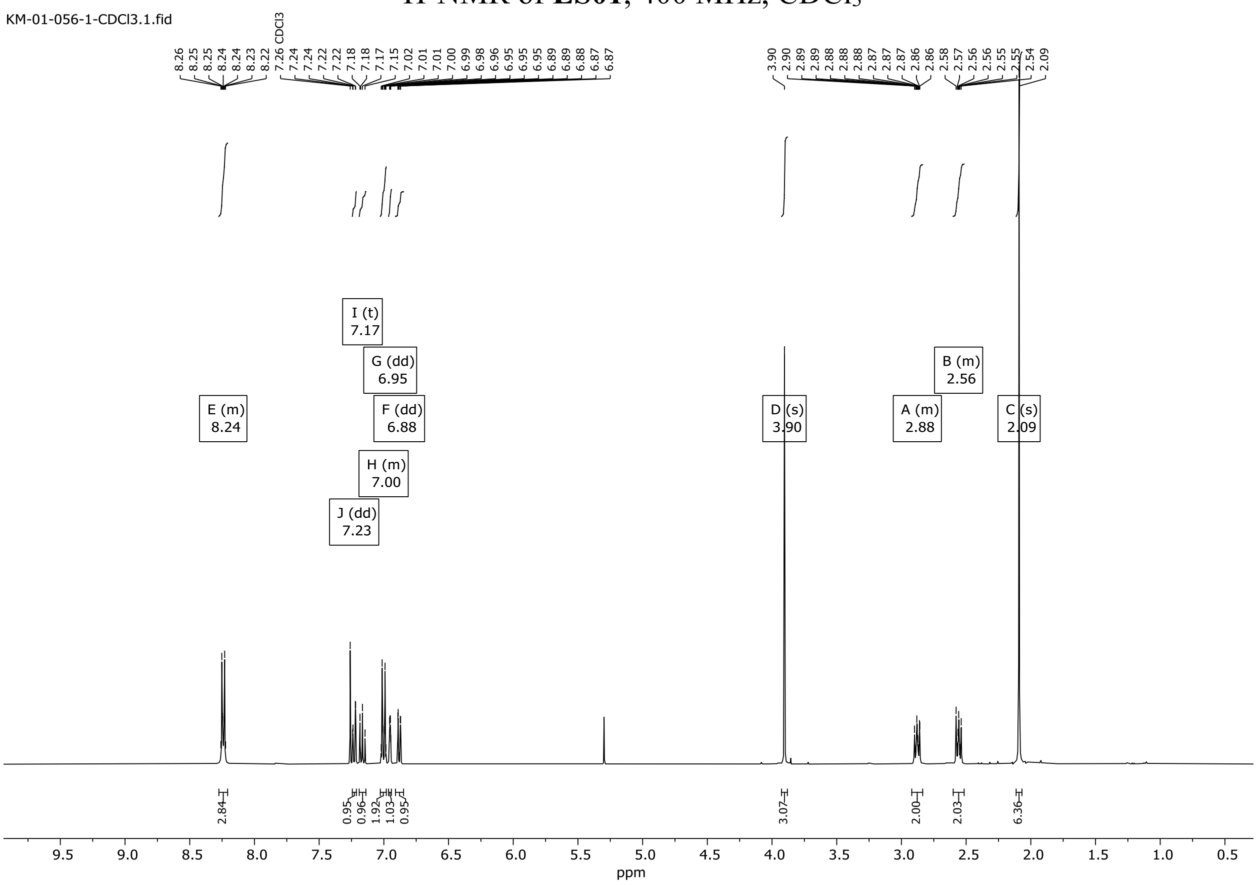

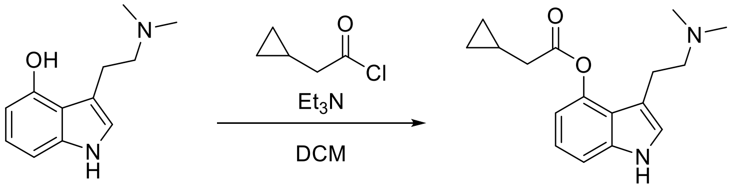

1 ES02

**3-(2-(Dimethylamino)ethyl)-1*H*-indol-4-yl 2-cyclopropylacetate (ES02)**

A solution of psilocin **1** (600 mg, 2.94 mmol) and triethylamine (823 µL, 5.87 mmol) in anhydrous DCM (29.4 mL) was cooled down to 0 °C. To it was added cyclopropylacetyl chloride (813 µL, 8.22 mmol, 2.8 eq) in 3 portions (1.2 eq + 1.2 eq + 0.4 eq; with 2-hour intervals) and the resulting mixture was warmed up to RT and stirred overnight. After 18 h, TLC (MeOH/DCM 20:80) showed full consumption of psilocin. The reaction was quenched with methanol (1 mL), and the volatiles were removed in vacuo. The crude residue was directly purified by FC chromatography on silica gel (24 g, MeOH/DCM 0:100 to 20:80, product eluting at 15% MeOH) to afford the product as a brown oil. $^1$H-NMR showed co-elution of cyclopropylacetic acid with the product. This isolated material was re-dissolved in DCM (25 mL) and extracted with sat'd aq. $NaHCO_3$ (1 x 25 mL). The aq. layer was back extracted with DCM (2 x 30 mL), washed with brine, dried over anhydrous $Na_2SO_4$, filtered and concentrated to afford the pure product **ES02** as a tan solid (510 mg, 61%).

HRMS-HESI: calculated: 287.1760; observed: 287.1753 m/z $[M+H]^+$

$^1$H NMR (400 MHz, $CDCl_3$) δ 8.20 (s, 1H), 7.18 (dd, J = 8.2, 1.0 Hz, 1H), 7.12 (dd, J = 8.2, 7.5 Hz, 1H), 6.94 (dt, J = 2.2, 1.0 Hz, 1H), 6.81 (dd, J = 7.5, 1.0 Hz, 1H), 2.98 – 2.86 (m, 2H), 2.64 – 2.55 (m, 4H), 2.30 (s, 6H), 1.32 – 1.17 (m, 1H), 0.71 – 0.59 (m, 2H), 0.31 (dt, J = 6.0, 4.7 Hz, 2H).

$^{1}$H-NMR of **ES02**, 400 MHz, $CDCl_3$

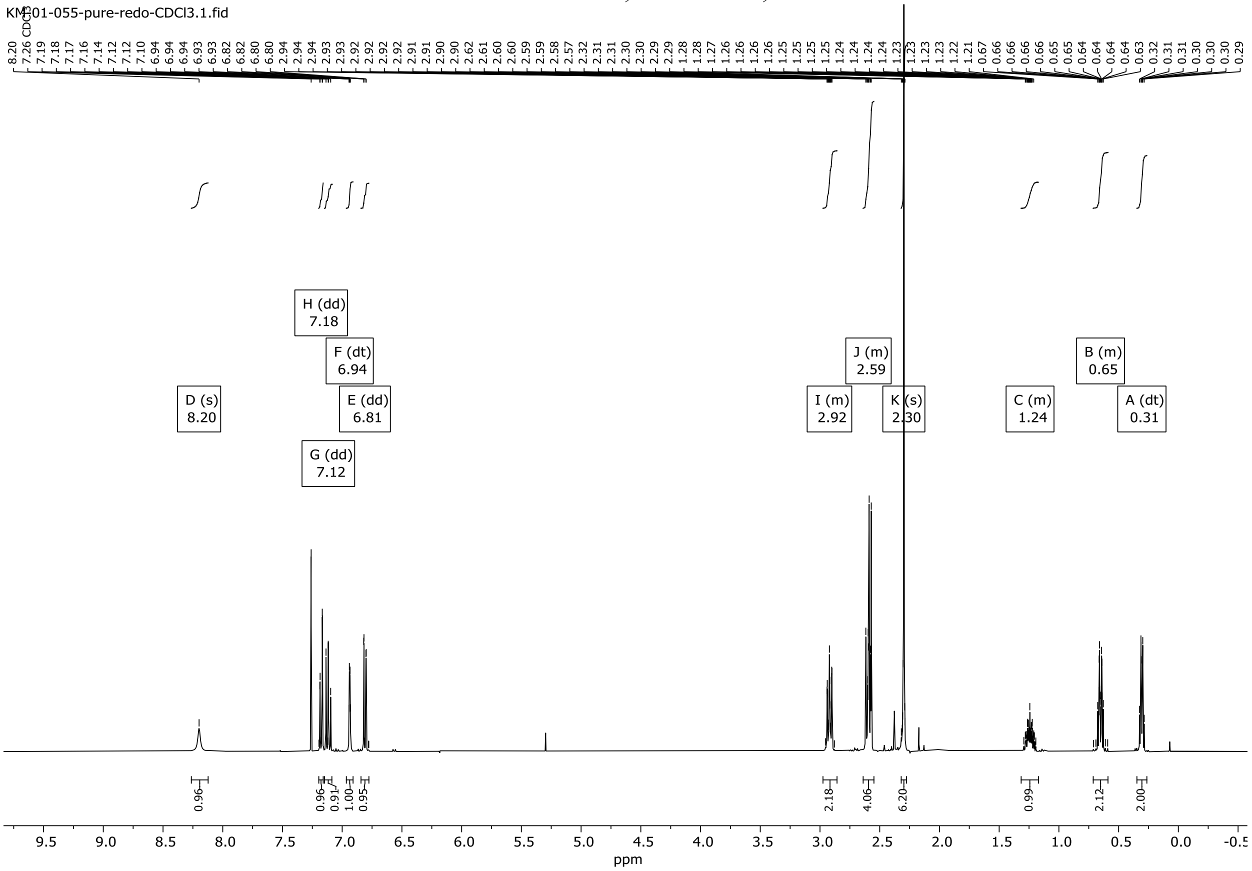

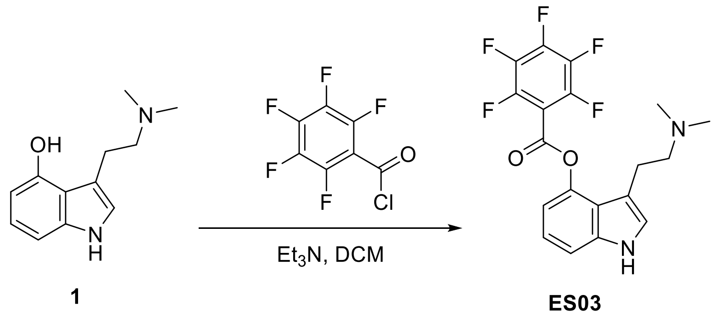


**3-(2-(Dimethylamino)ethyl)-1*H*-indol-4-yl 2,3,4,5,6-pentafluorobenzoate (ES03)**
To a suspension of psilocin **1** (30 mg, 147 μmol) in dry DCM (8 ml) was added of triethylamine (22 μL, 157 μmol) and pentafluorobenzoyl chloride (34.6 μL, 240 μmol). The mixture was stirred at room temperature for 2 hours and the reaction was monitored by TLC. The reaction mixture was then quenched by 0.5 M aq. HCl and extracted with DCM (x3). The combined organic extracts were washed with water, brine, dried over anhydrous $MgSO_4$ and filtered. The filtrate was concentrated under reduced pressure and purified by flash chromatography on silica gel (5% methanol in DCM) to provide the product **ES03** as an off-white solid (21.2 mg, 36% yield).
HRMS-HESI: calculated: 399.1126; observed: 399.1120 m/z $[M+H]^+$
$^1$H NMR (400 MHz, MeOD) δ 7.39 (dd, *J* = 8.2, 0.8 Hz, 1H), 7.30 (d, *J* = 1.0 Hz, 1H), 7.20 (t, *J* = 8.0 Hz, 1H), 6.93 (dd, *J* = 7.7, 0.8 Hz, 1H), 3.41 (dd, *J* = 7.9, 6.9 Hz, 2H), 3.25 – 3.13 (m, 2H), 2.84 (s, 6H).

[1]H-NMR of **ES03**, 400 MHz, MeOD

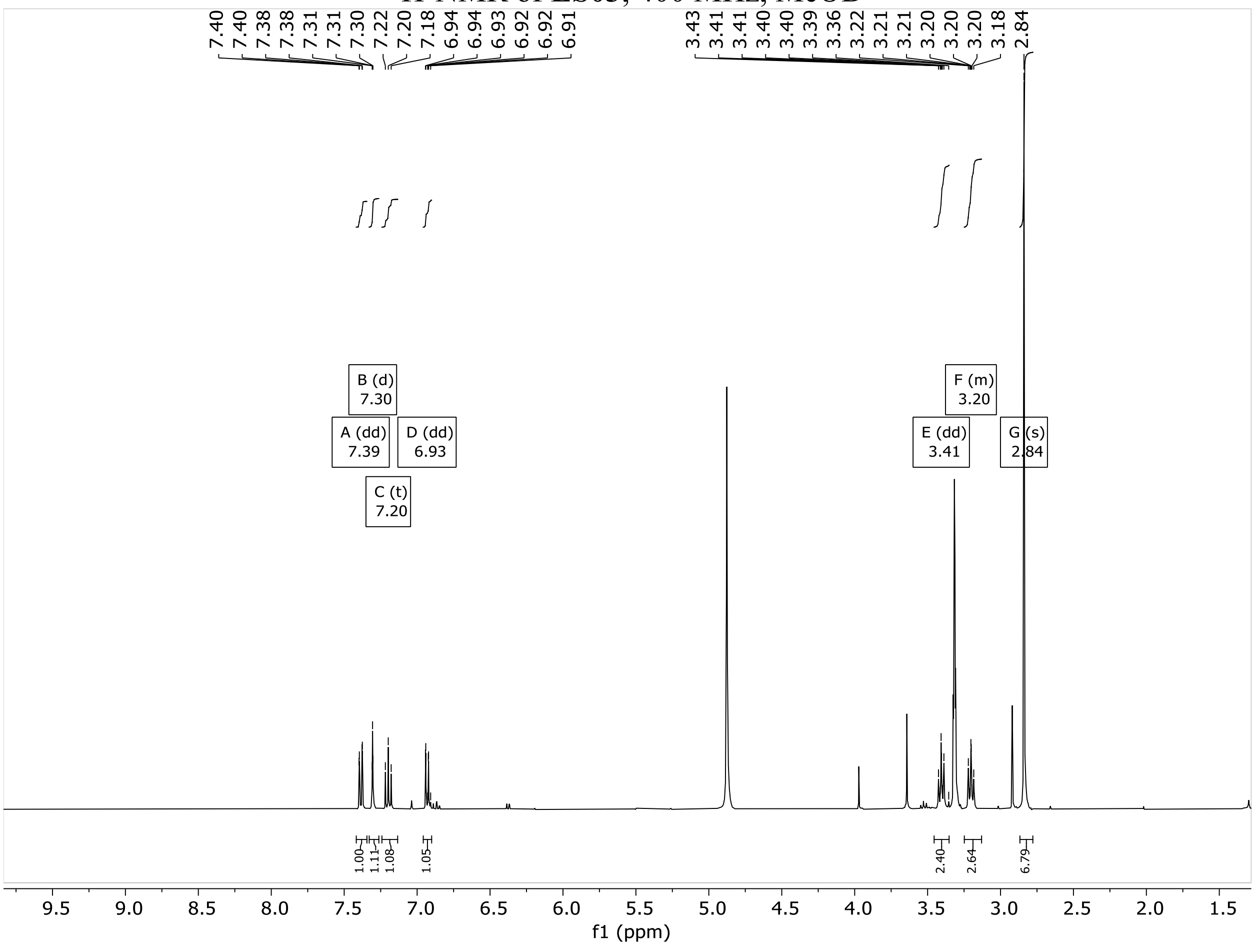


YC-95-220408/1 — Ethyl Crotonate 5% / CDCl3 HW=.27Hz/power -30/gain 122

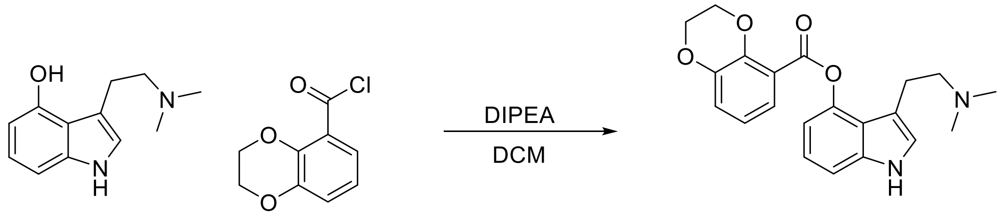

1 ES04

**3-(2-(Dimethylamino)ethyl)-1*H*-indol-4-yl 2,3-dihydrobenzo[b][1,4]dioxine-5-carboxylate (ES04)**

To a solution of **1** (74.7 mg, 366 µmol) and N,N-Diisopropylethylamine (100 µL, 574 µmol) in DCM (2 mL) was added, in a dropwise manner, a solution of 2,3-dihydrobenzo[b][1,4]dioxine-5-carbonyl chloride (77.2 mg, 373 µmol) in DCM (1 mL). The mixture was left to react at RT. After 3 hours the mixture contained very little starting material (TLC) and the reaction mixture was poured into a separatory funnel containing 10 mL of water and 10 mL DCM. The aqueous phase was extracted with DCM (3x10 mL), all organic phases were combined, washed with brine and dried over anhydrous $MgSO_4$. After filtration the solvent was removed under reduced pressure leaving a beige solid. The crude product was purified by flash chromatography on silica gel (1:9 MeOH/DCM) to yield the desired product **ES04** (68 mg, 51%) as a white solid.

HRMS-HESI: calculated: 367.1652; observed: 367.1651 m/z $[M+H]^+$

$^1$H NMR (400 MHz, DMSO-$d_6$) δ 11.06 (s, 1H), 7.57 (dd, *J* = 7.8, 1.6 Hz, 1H), 7.26 (dd, *J* = 8.1, 0.8 Hz, 1H), 7.20 – 7.13 (m, 2H), 7.07 (t, *J* = 7.9 Hz, 1H), 6.98 (t, *J* = 7.9 Hz, 1H), 6.74 (dd, *J* = 7.6, 0.8 Hz, 1H), 4.38 – 4.28 (m, 4H), 2.79 – 2.71 (m, 2H), 2.44 (dd, *J* = 9.2, 6.7 Hz, 2H), 2.01 (s, 6H).

[1]H-NMR of **ES04**, 400 MHz, DMSO-$d_6$

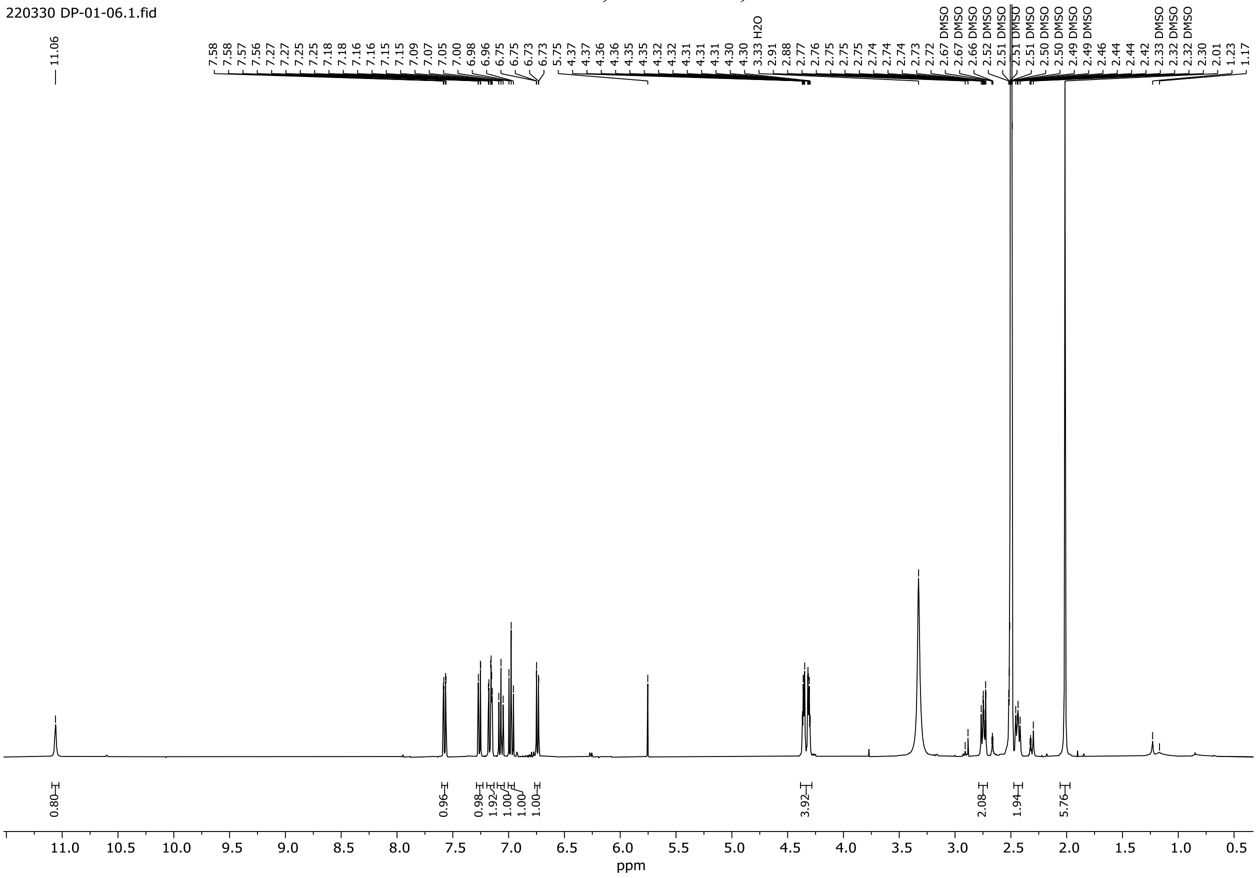

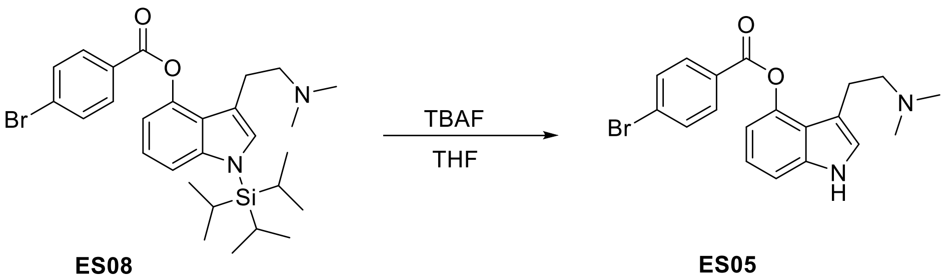


**ES08** **ES05**

**3-(2-(Dimethylamino)ethyl)-1*H*-indol-4-yl 4-bromobenzoate (ES05)**

To a solution of **ES08** (175 mg, 322 µmol) in THF (2.0 mL) was added TBAF (tetrabutylammonium fluoride) (1 M in THF, 483 µL, 483 µmol) dropwise. After 30 min, the mixture was poured into a separatory funnel containing water (15 mL) and DCM (15 mL), the aqueous phase was extracted with DCM (3x15 mL), the combined organic layers were washed with brine, dried over anhydrous $MgSO_4$, filtered and concentrated. The crude residue was purified by flash chromatography on silica gel (1:9 MeOH/DCM) to afford the product (**ES05**) as a white powder (80.0 mg, 64%).

HRMS-HESI: calculated: 387.0703; observed: 387.0704 m/z $[M+H]^+$

$^1$H NMR (400 MHz, $CDCl_3$) δ 7.87 (dd, *J* = 8.2, 0.9 Hz, 1H), 7.66 (d, *J* = 8.5 Hz, 2H), 7.58 (d, *J* = 8.4 Hz, 2H), 7.23 (d, *J* = 8.1 Hz, 1H), 6.89 (s, 1H), 6.81 (dd, *J* = 8.0, 0.9 Hz, 1H), 2.93 – 2.86 (m, 2H), 2.79 – 2.73 (m, 2H), 2.42 (s, 6H).

$^1$H-NMR of **ES05**, 400 MHz, $CDCl_3$

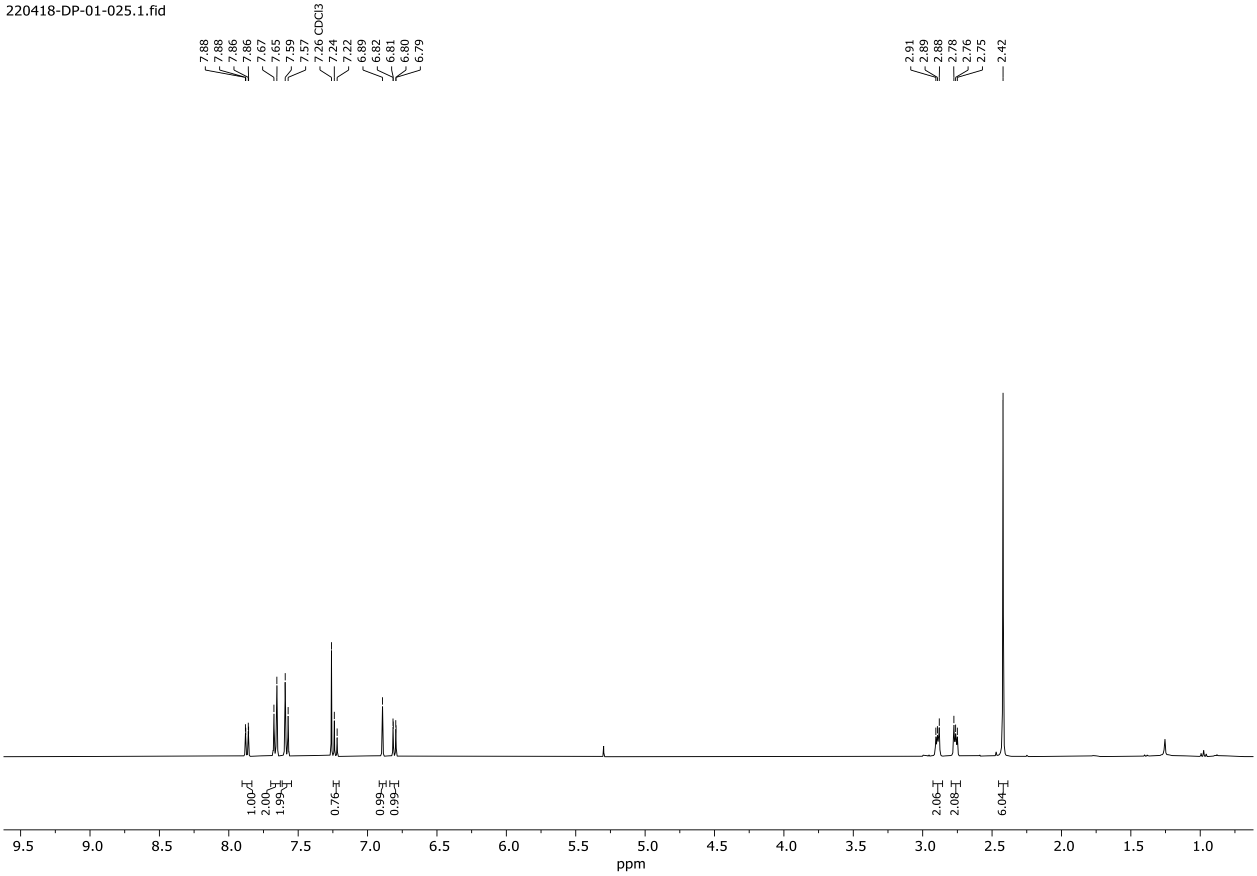

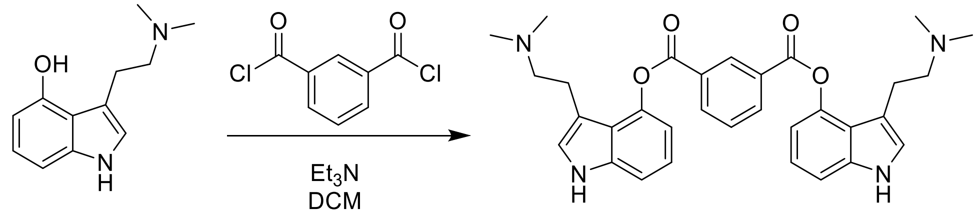


**1** **ES06**

**Bis(3-(2-(dimethylamino)ethyl)-1*H*-indol-4-yl) isophthalate (ES06)**

To a flame-dried flask was added compound **1** (50 mg, 0.25 mmol), and anhydrous dichloromethane (1 mL) under argon. Triethylamine (34 µL, 0.25 mmol) was added, followed by isophthaloyl chloride (25 mg, 0.12 mmol) dissolved in anhydrous dichloromethane (1 mL). The mixture was refluxed overnight, then dried and directly purified by flash chromatography on silica gel (4 g, 10 to 20% MeOH/DCM) to yield **ES06** (19.6 mg, 30%) as a tan oil.

HRMS-HESI: calculated: 539.2653; observed: 539.2645 m/z $[M+H]^+$

$^1$H NMR (400 MHz, MeOD) δ 9.13 (td, J = 1.8, 0.6 Hz, 1H), 8.70 (dd, J = 7.8, 1.8 Hz, 2H), 7.95 (td, J = 7.8, 0.6 Hz, 1H), 7.42 – 7.37 (m, 2H), 7.29 (t, J = 0.9 Hz, 2H), 7.24 – 7.18 (m, 2H), 6.93 (dd, J = 7.7, 0.8 Hz, 2H), 3.41 – 3.36 (m, 4H), 3.20 – 3.14 (m, 4H), 2.73 (s, 12H).

$^1$H-NMR of **ES06**, 400 MHz, MeOD

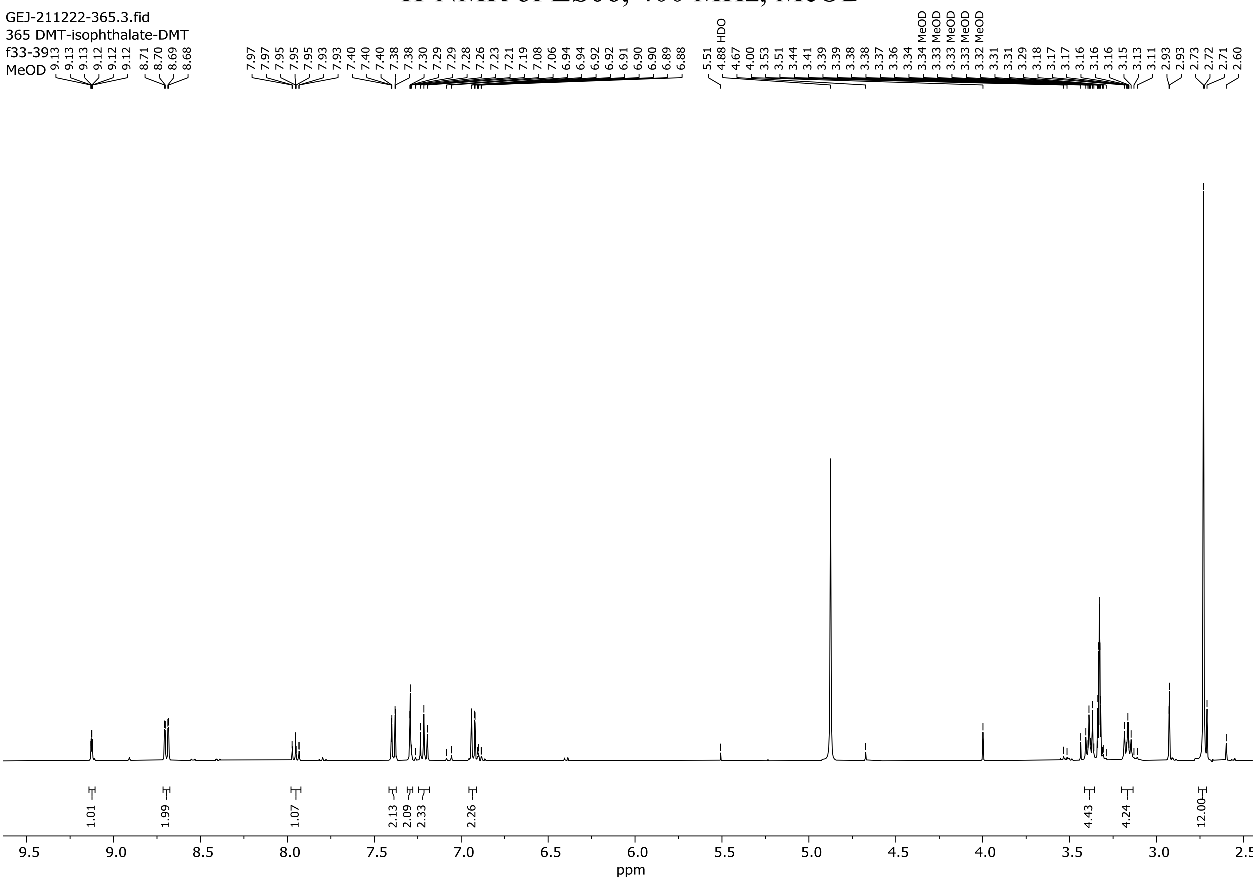

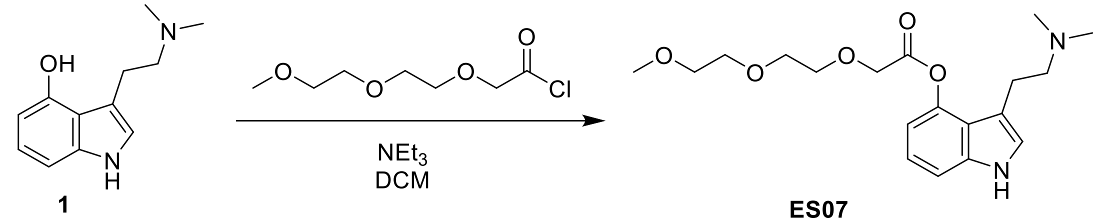


1 ES07

**3-(2-(Dimethylamino)ethyl)-1*H*-indol-4-yl 2-(2-(2-methoxyethoxy)ethoxy)acetate (ES07)**
Compound **1** (100 mg, 0.49 mmol) was suspended in anhydrous DCM (0.5 mL) under argon atmosphere. Triethylamine (0.10 mL, 0.73 mmol) was added, followed by m-PEG2-$CH_2$ acid chloride (96 mg, 0.49 mmol) diluted in DCM (0.2 mL). The resulting mixture was stirred at room temperature for 23 hours and monitored by TLC (20% MeOH/DCM). The volatiles were removed under reduced pressure, and the crude residue was directly purified by flash column chromatography on silica gel (12 g, 10% MeOH/DCM). The resulting semi-pure product was further purified by a second flash column chromatography on silica gel (4 g, 8 to 10% MeOH/DCM) to yield **ES07** (3.9 mg, 2.2%) as a colorless oil.

HRMS-HESI: calculated: 365.2071; observed: 365.2068 m/z $[M+H]^+$
$^1$H NMR (400 MHz, MeOD) δ 7.32 (dd, J = 8.2, 0.8 Hz, 1H), 7.27 (s, 1H), 7.15 (t, J = 8.0 Hz, 1H), 6.88 (dd, J = 7.8, 0.8 Hz, 1H), 4.58 (s, 2H), 3.92 – 3.85 (m, 2H), 3.78 – 3.72 (m, 2H), 3.70 – 3.64 (m, 2H), 3.59 – 3.52 (m, 2H), 3.48 – 3.42 (m, 2H), 3.37 (s, 3H), 3.26 – 3.17 (m, 2H), 2.94 (s, 6H).

$^1$H-NMR of **ES07**, 400 MHz, MeOD

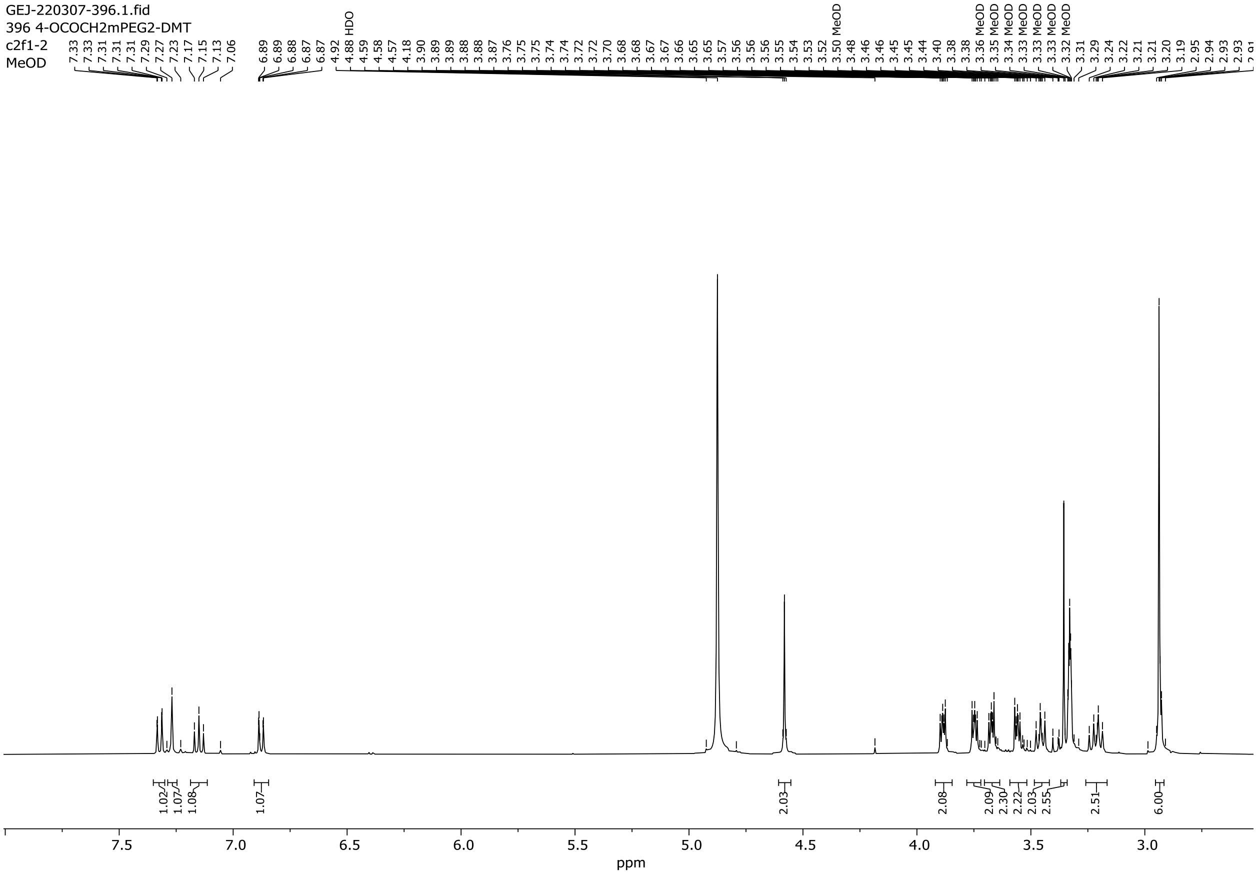

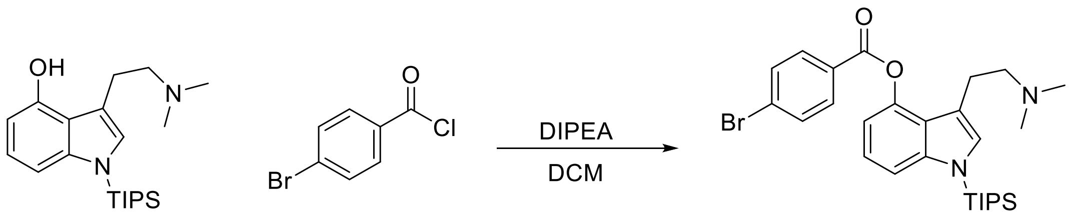


**7** **ES08**

**3-(2-(Dimethylamino)ethyl)-1-(triisopropylsilyl)-1*H*-indol-4-yl 4-bromobenzoate (ES08)**

To a solution of N,N-diisopropylethylamine (114 µL, 653 µmol) and **7** (150 mg, 416 µmol) in DCM (2.50 mL) was added, in a dropwise manner, a solution of 4-bromobenzoyl chloride (93.2 mg, 416 µmol) in DCM (1.25 mL). The mixture was left to react at RT. After 3 hours the mixture contained very little starting material and the reaction mixture was poured into a separatory funnel containing 10 mL of water and 10 mL DCM. The aqueous phase was extracted with DCM (3x10 mL), all organic phases were combined, washed with brine, and dried over magnesium sulfate. Purification was carried out by flash column chromatography on silica gel (1:9 MeOH/DCM) to afford **ES08** (193 mg, 85%) as a colorless oil.

HRMS-HESI: calculated: 543.2037; observed: 543.2036 m/z $[M+H]^+$

$^1$H NMR (400 MHz, $CDCl_3$) δ 8.15 (d, J = 8.6 Hz, 2H), 7.68 (d, J = 8.6 Hz, 2H), 7.38 (dd, J = 8.3, 0.8 Hz, 1H), 7.17 – 7.10 (m, 1H), 7.02 (s, 1H), 6.90 (dd, J = 7.7, 0.7 Hz, 1H), 2.96 – 2.85 (m, 2H), 2.59 (t, J = 8.1 Hz, 2H), 2.12 (s, 6H), 1.68 (m, J = 7.5 Hz, 3H), 1.14 (d, J = 7.5 Hz, 18H).

$^1$H-NMR of **ES08**, 400 MHz, $CDCl_3$

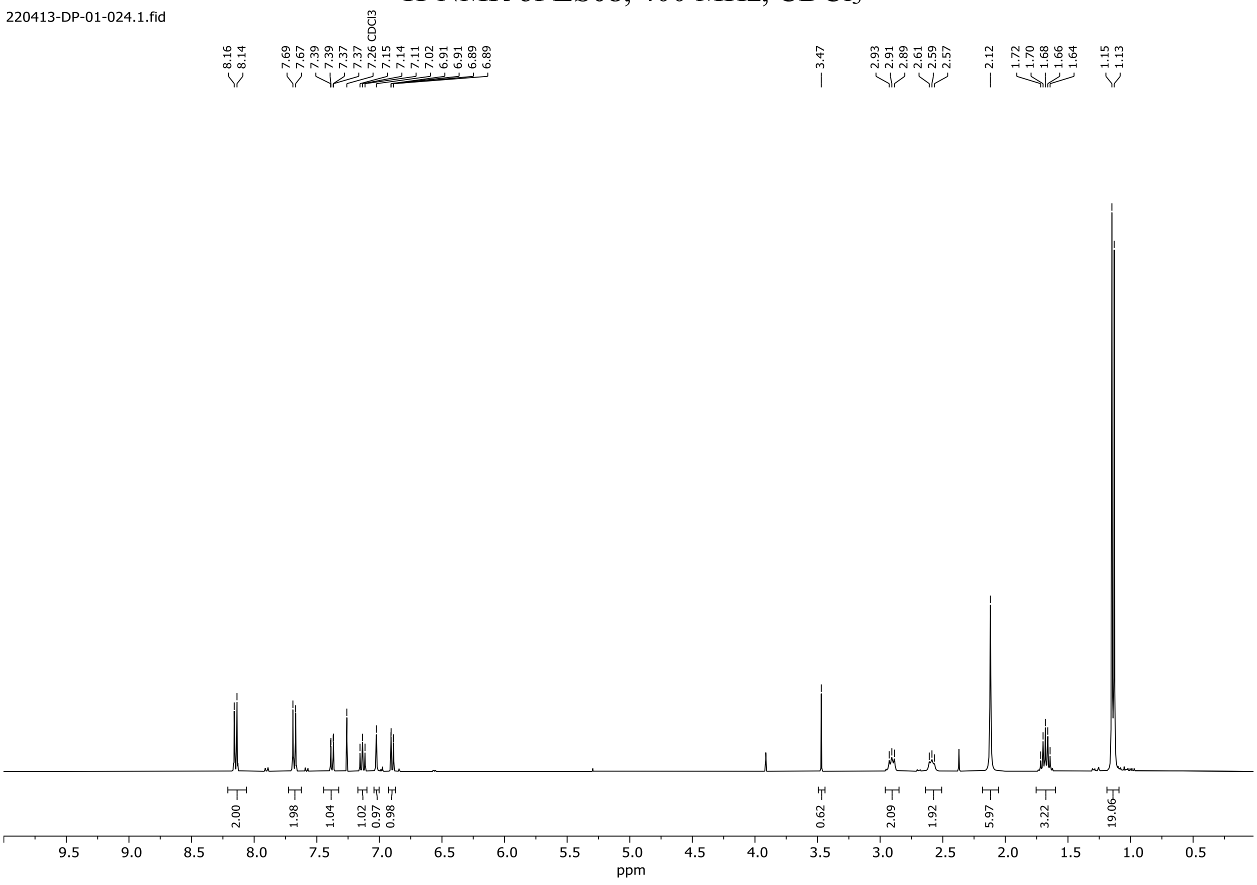

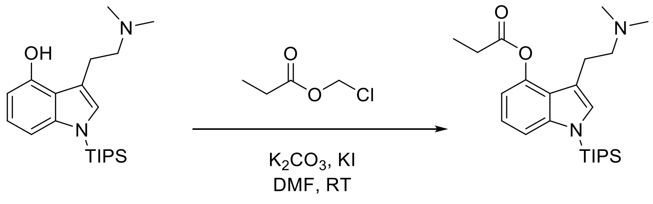


7 ES09

**3-(2-(Dimethylamino)ethyl)-1-(triisopropylsilyl)-1*H*-indol-4-yl propionate (ES09)**

To a solution of *N*-TIPS-Psilocin **7** (180 mg, 0.5 mmol) in dry DMF (4 ml) under argon was added potassium carbonate (69 mg, 0.5 mmol) and potassium iodide (82.8 mg, 0.5 mmol). The mixture was stirred at room temperature for 10 minutes and then chloromethyl propanoate (61.2 mg, 500 mmol) was added. The resulting mixture was stirred at room temperature for 2 hours. The reaction was quenched by addition of water. The mixture was then extracted with DCM (3 x 10 mL), the organic layers were combined, washed with brine, dried over anhydrous $MgSO_4$, filtered and concentrated. The crude residue was purified by flash chromatography on silica gel (0% to 10% MeOH/DCM) to afford the product **ES09** as a pale-yellow oil (140.4 mg, 67.5%).

HRMS-HESI: calculated: 417.2914; observed: 417.2932 m/z $[M+H]^+$

$^1$H NMR (400 MHz, MeOD) δ 7.43 (dd, J = 8.4, 0.8 Hz, 1H), 7.22 (s, 1H), 7.13 (t, J = 8.1 Hz, 1H), 6.81 (dd, J = 7.7, 0.7 Hz, 1H), 3.10 – 2.92 (m, 4H), 2.78 (q, J = 7.5 Hz, 2H), 2.59 (s, 6H), 1.76 (h, J = 7.5 Hz, 3H), 1.30 (t, J = 7.5 Hz, 3H), 1.18 (d, J = 7.5 Hz, 18H).

$^1$H-NMR of **ES09**, 400 MHz, MeOD

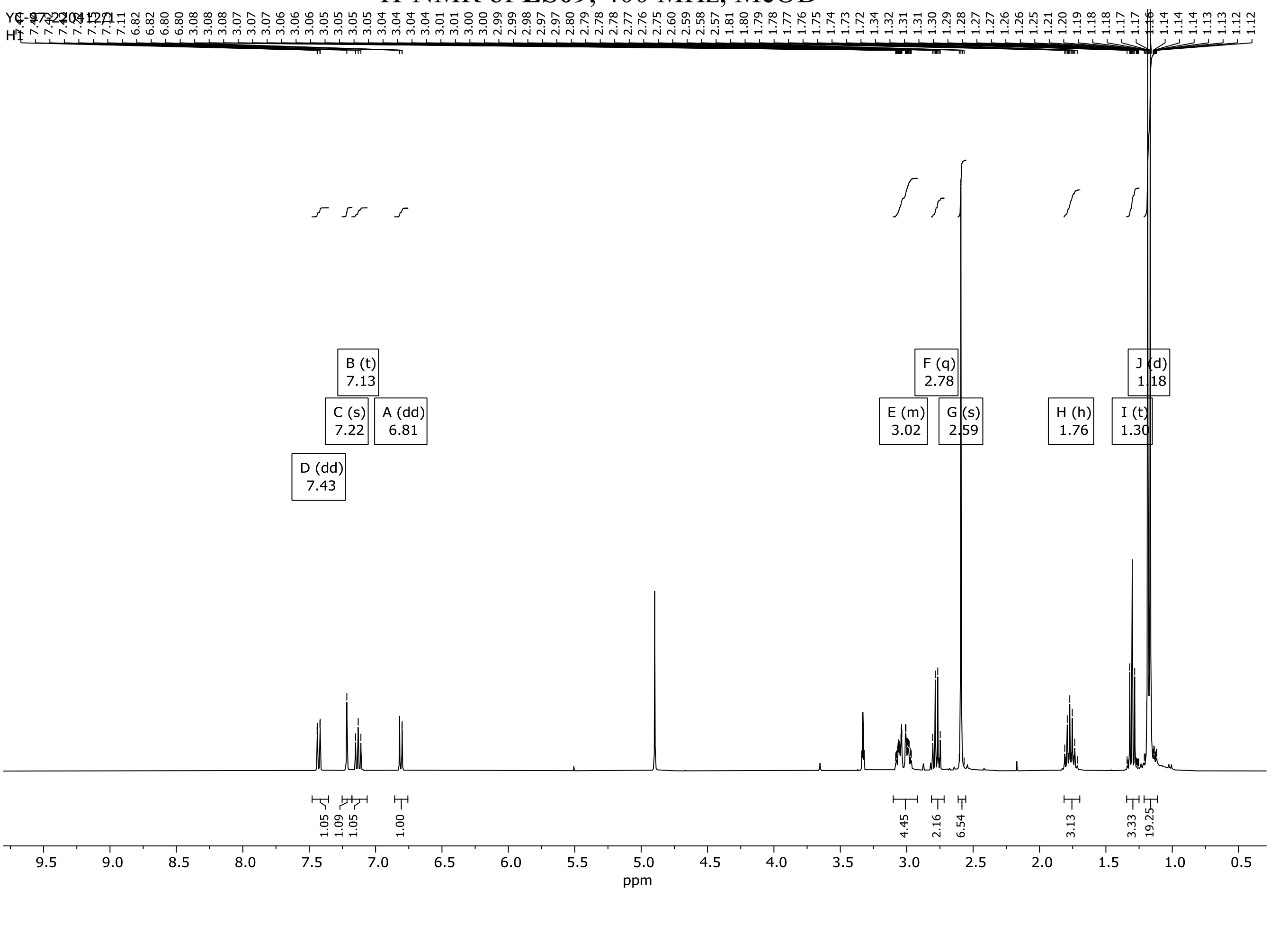

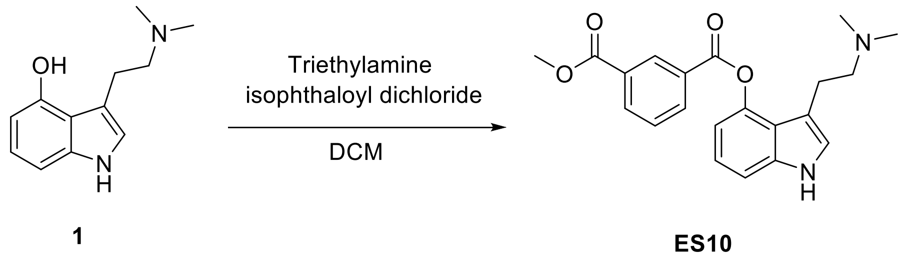

### 3-(2-(Dimethylamino)ethyl)-1*H*-indol-4-yl methyl isophthalate (ES10)

To a suspension of compound **1** (102 mg, 0.50 mmol) in dry DCM (1.5 mL) under argon atmosphere was added triethylamine (70 µL, 0.50 mmol) followed by isophthaloyl dichloride (51 mg, 0.25 mmol) dissolved in dry DCM (0.5 mL). The reaction mixture was allowed to stir at room temperature for 18 hours. The crude reaction mixture was concentrated and directly purified by flash chromatography on silica gel (4 g, 10 to 20% MeOH/DCM) to afford a mixture of products. This mixture was further purified by a second flash column chromatography on silica gel (4 g, 10% MeOH/DCM) to yield **ES10** (7 mg, 8%) as a colorless oil.

HRMS-HESI: calculated: 367.1652; observed: 367.1650 m/z $[M+H]^+$

$^1$H NMR (400 MHz, MeOD) δ 8.91 (td, J = 1.8, 0.6 Hz, 1H), 8.54 (ddd, J = 7.8, 1.8, 1.2 Hz, 1H), 8.40 (dt, J = 7.9, 1.4 Hz, 1H), 7.80 (td, J = 7.8, 0.6 Hz, 1H), 7.38 (dd, J = 8.2, 0.8 Hz, 1H), 7.27 (d, J = 0.8 Hz, 1H), 7.21 (t, J = 7.9 Hz, 1H), 6.91 (dd, J = 7.7, 0.8 Hz, 1H), 4.00 (s, 3H), 3.29 (dd, J = 8.6, 6.8 Hz, 2H), 3.12 – 3.06 (m, 2H), 2.65 (s, 6H).

$^1$H-NMR of **ES10**, 400 MHz, MeOD

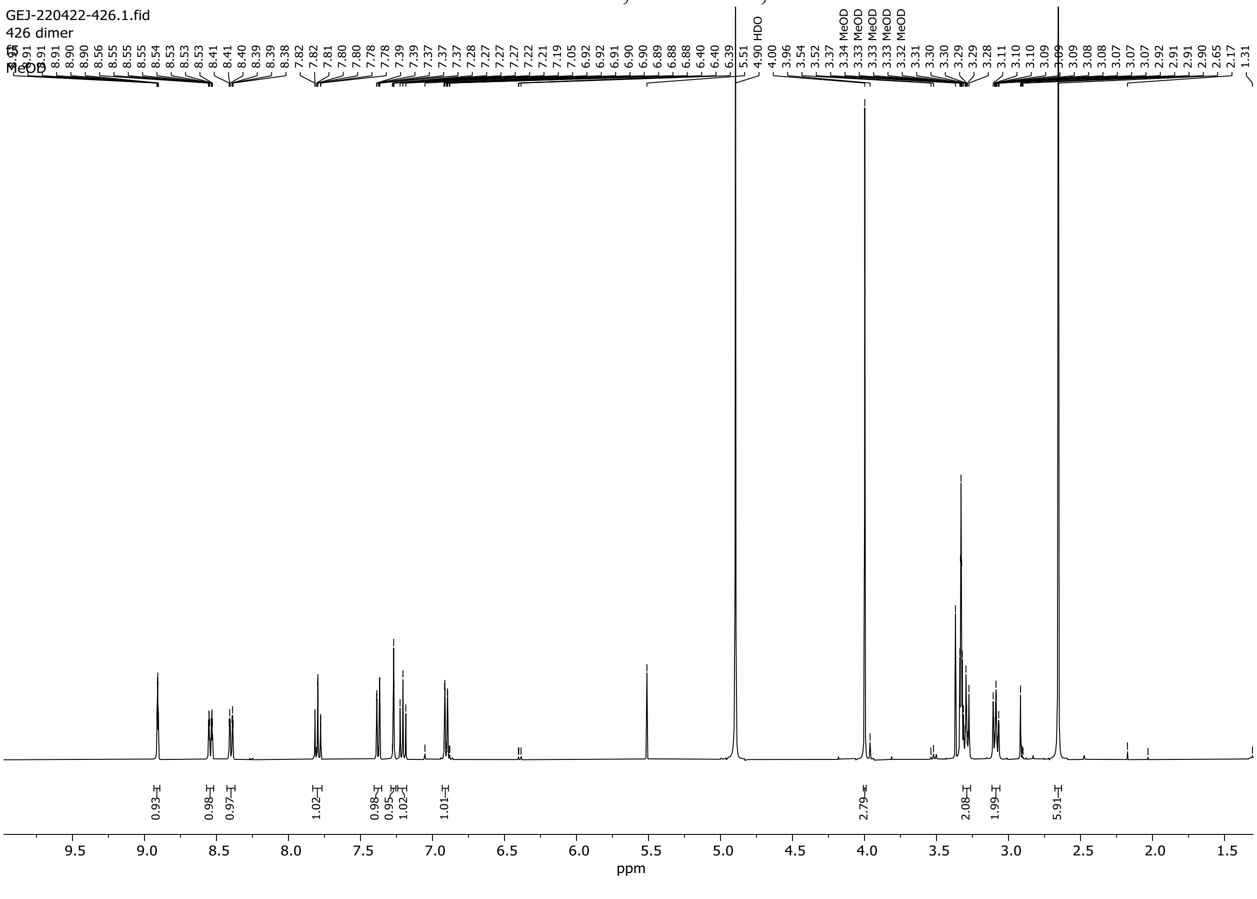

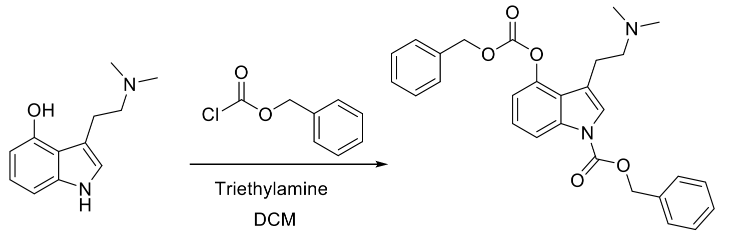


**1** **C01**

**Benzyl 4-(((benzyloxy)carbonyl)oxy)-3-(2-(dimethylamino)ethyl)-1*H*-indole-1-carboxylate (C01)**

To a suspension of psilocin **1** (98 mg, 0.48 mmol) in dry DCM (2.0 mL) under argon atmosphere was added triethylamine (0.12 mL, 0.87 mmol) followed by benzyl chloroformate (75 µL, 0.52 mmol). The reaction mixture was stirred at room temperature for 4 hours until the starting material was no longer visualized on TLC (20% MeOH/DCM). Following dilution with DCM (10 mL) and methanol (1 mL), the reaction mixture was washed with brine, dried over anhydrous $MgSO_4$, filtered and concentrated to afford the crude product as an amber oil. Purification by flash column chromatography on silica gel (12 g, 0 to 10% MeOH/DCM) yielded **C01** (58 mg, 25%) as a white solid.

HRMS-HESI: calculated: 473.2071; observed: 473.2059 m/z $[M+H]^+$

$^1H$ NMR (400 MHz, MeOD) δ 7.57 – 7.35 (m, 10H), 7.31 (d, J = 8.1 Hz, 1H), 7.26 (s, 1H), 7.11 (tt, J = 8.1, 1.5 Hz, 1H), 6.96 (d, J = 7.7 Hz, 1H), 5.26 – 5.19 (m, 2H), 4.45 (d, J = 2.4 Hz, 2H), 3.53 – 3.43 (m, 2H), 3.26 – 3.15 (m, 2H), 2.95 – 2.90 (m, 6H).

$^1H$-NMR of **C01**, 400 MHz, $CDCl_3$

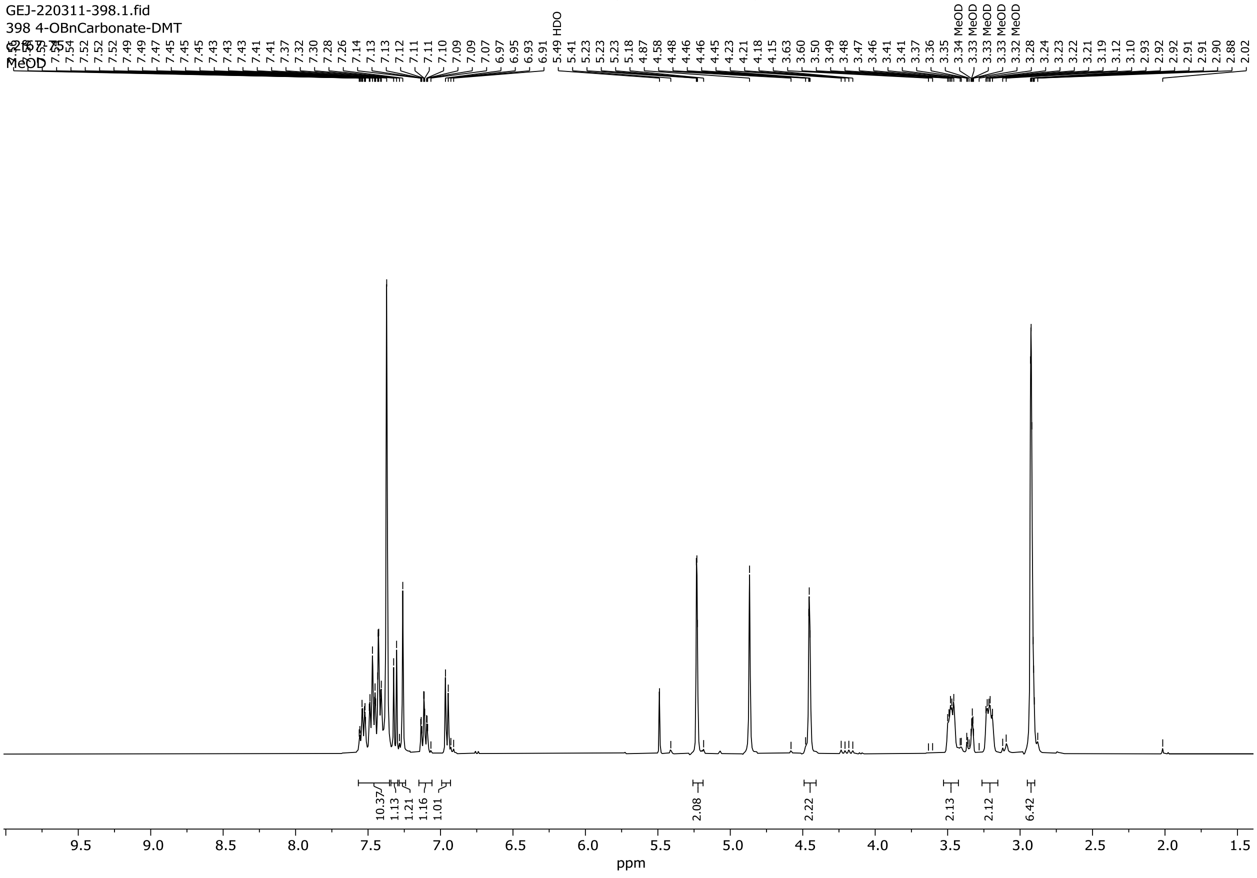

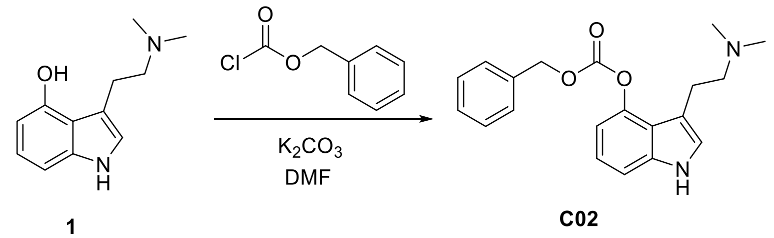

#### Benzyl (3-(2-(dimethylamino)ethyl)-1*H*-indol-4-yl) carbonate (C02)

To a suspension of psilocin **1** (100 mg, 0.49 mmol) and potassium carbonate (68 mg, 0.49 mmol) in dry DMF (1.2 mL) under argon atmosphere was added benzyl chloroformate (70 µL, 0.49 mmol). The reaction was allowed to stir at room temperature for 4 hours until completion as determined by TLC (20% MeOH/DCM). Following dilution with water (10 mL), the reaction mixture was extracted with DCM (3 x 10 mL). The combined organic extracts were washed with brine (10 mL), dried over anhydrous $MgSO_4$, filtered and concentrated. Purification by flash column chromatography on silica gel (12 g, 0 to 10% MeOH/DCM) yielded **C02** (38 mg, 23%) as a light-yellow oil.

HRMS-HESI: calculated: 339.1703; observed: 339.1701 m/z $[M+H]^+$

$^1$H NMR (400 MHz, MeOD) δ 7.63 (d, J = 8.3 Hz, 1H), 7.55 – 7.49 (m, 2H), 7.46 – 7.36 (m, 3H), 7.33 (s, 1H), 7.09 (t, J = 8.1 Hz, 1H), 6.60 (dd, J = 7.9, 0.8 Hz, 1H), 5.44 (s, 2H), 3.02 (td, J = 7.3, 1.0 Hz, 2H), 2.78 (t, J = 7.1 Hz, 2H), 2.38 (s, 6H).

$^1$H-NMR of **C02**, 400 MHz, MeOD

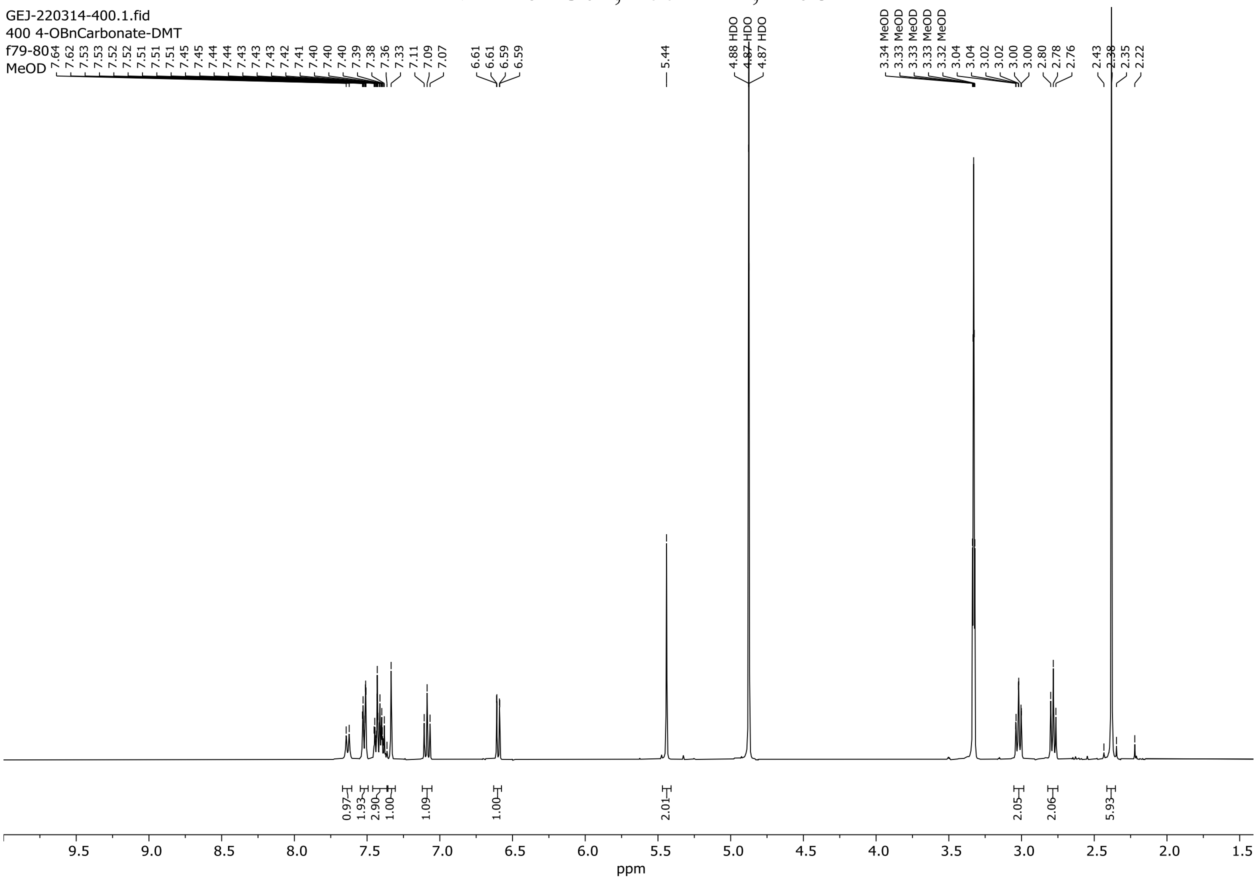

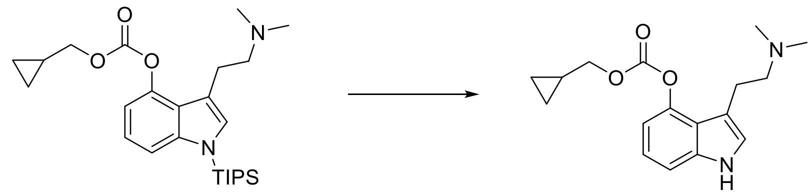

**C05** **C03**

**Cyclopropylmethyl (3-(2-(dimethylamino)ethyl)-1*H*-indol-4-yl) carbonate (C03)**

To a solution of **C05** (165 mg, 360 µmol) in dry THF (1.80 mL) was added TBAF (tetrabutylammonium fluoride) solution (1 M in THF, 540 µL, 540 µmol) dropwise at 0 °C. After 1 h, water (2 mL) was added, the aq. layer was separated and extracted with DCM (3 x 15 mL). The combined organic layers were washed with brine, dried over anhydrous $Na_2SO_4$, filtered and concentrated. The crude product was purified by flash chromatography using silica gel (MeOH/DCM 8:92) to afford the pure product **C03** as a colorless oil (32 mg, 29%).

HRMS-HESI: calculated: 303.1703; observed: 303.1702 m/z $[M+H]^+$

$^1$H NMR (400 MHz, $CDCl_3$) δ 8.35 (s, 1H), 7.22 (d, *J* = 8.1 Hz, 1H), 7.13 (t, *J* = 7.9 Hz, 1H), 6.97 (dt, *J* = 2.1, 1.0 Hz, 1H), 6.91 (d, *J* = 7.7 Hz, 1H), 4.12 (d, *J* = 7.4 Hz, 2H), 3.03 – 2.93 (m, 2H), 2.72 – 2.65 (m, 2H), 2.38 (s, 6H), 1.35 – 1.21 (m, 1H), 0.71 – 0.59 (m, 2H), 0.43 – 0.34 (m, 2H).

$^1$H-NMR of **C03**, 400 MHz, $CDCl_3$

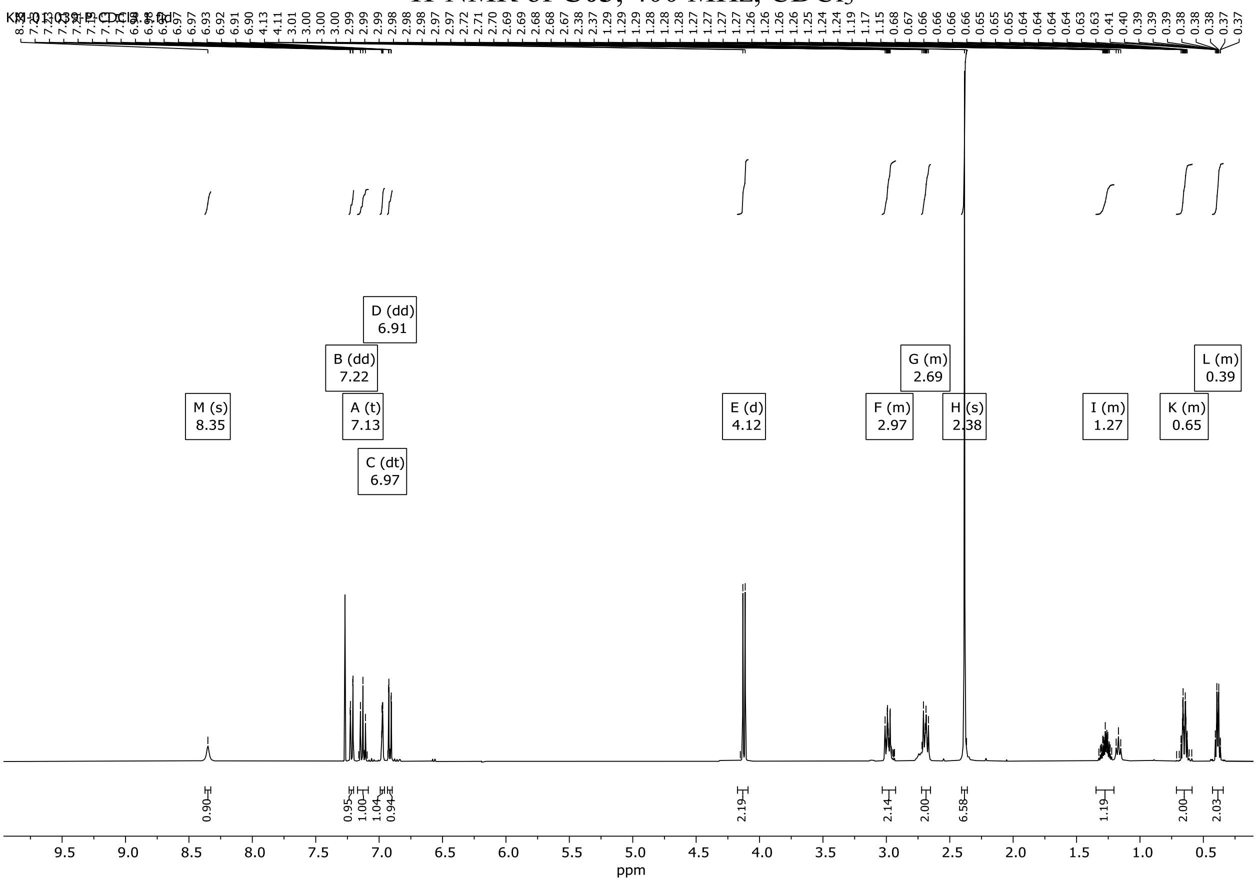

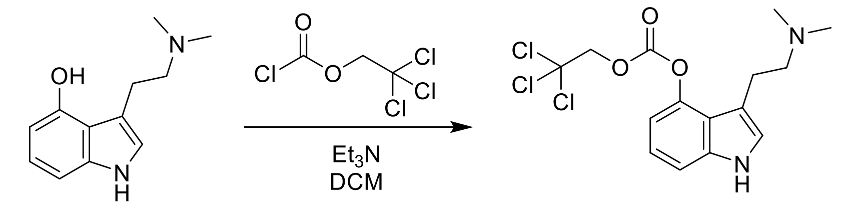


**1** **C04**

**3-(2-(Dimethylamino)ethyl)-1*H*-indol-4-yl (2,2,2-trichloroethyl) carbonate (C04)**

Compound **1** (100 mg, 0.49 mmol) was dissolved in anhydrous dichloromethane (2 mL) under argon. To this solution was added triethylamine (137 µL, 0.98 mmol) followed by 2,2,2-trichloroethyl chloroformate (135 µL, 0.98 mmol). The mixture was stirred at room temperature for 4 h until the reaction was complete as determined by TLC (20% MeOH/DCM). After dilution with dichloromethane (10 mL), the reaction mixture was washed with brine (15 mL), dried over anhydrous $MgSO_4$, and concentrated under reduced pressure. The crude product was purified by flash chromatography on silica gel (12 g, 2.5 to 10% MeOH/DCM) to yield **C04** (10 mg, 5.5%) as a colourless oil.

HRMS-HESI: calculated: 379.0378; observed: 379.0378 m/z $[M+H]^+$

$^1H$ NMR (400 MHz, MeOD) δ 7.69 (dd, J = 8.3, 0.8 Hz, 1H), 7.36 (d, J = 1.0 Hz, 1H), 7.14 (t, J = 8.1 Hz, 1H), 6.65 (dd, J = 7.9, 0.8 Hz, 1H), 5.15 (s, 2H), 3.08 – 3.01 (m, 2H), 2.80 (dd, J = 7.5, 6.7 Hz, 2H), 2.39 (s, 6H).

$^1H$-NMR of **C04**, 400 MHz, MeOD

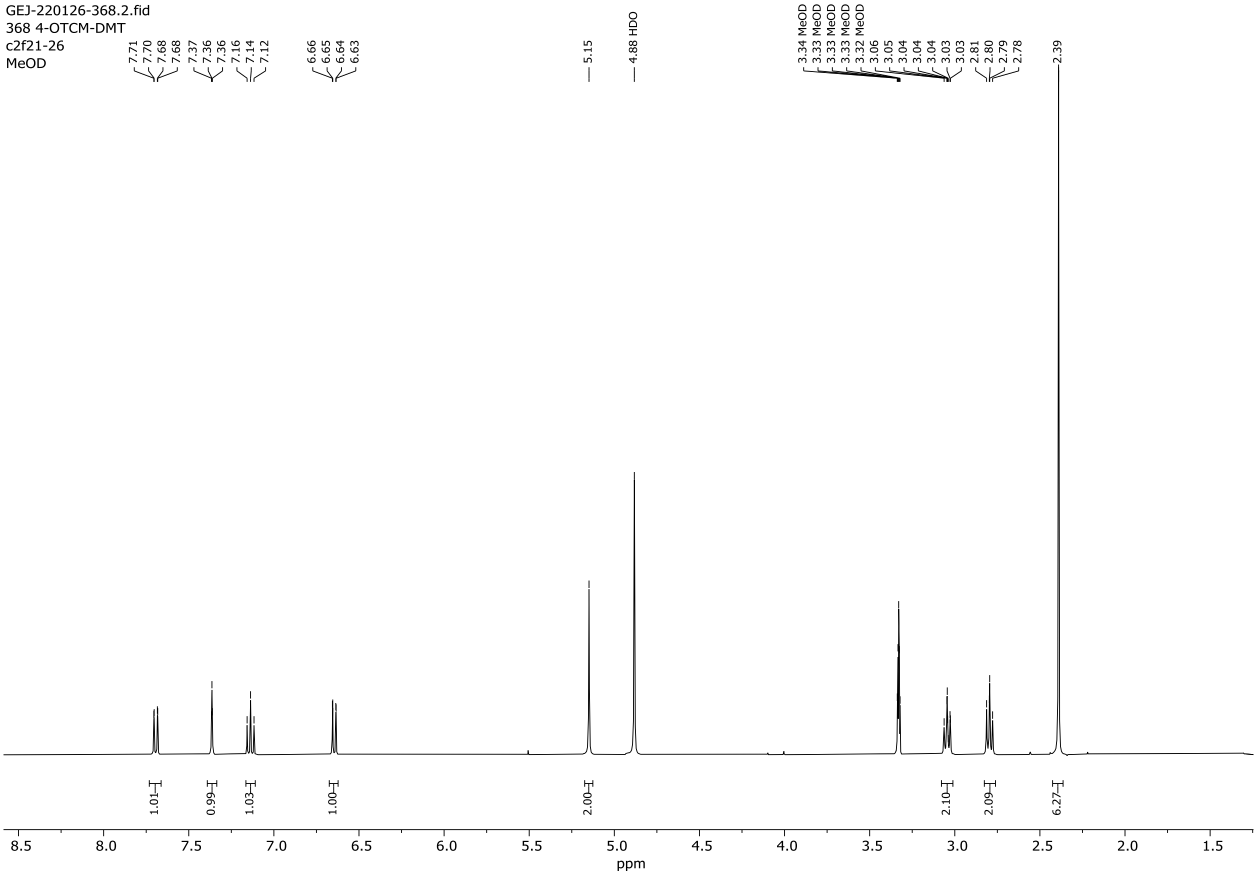

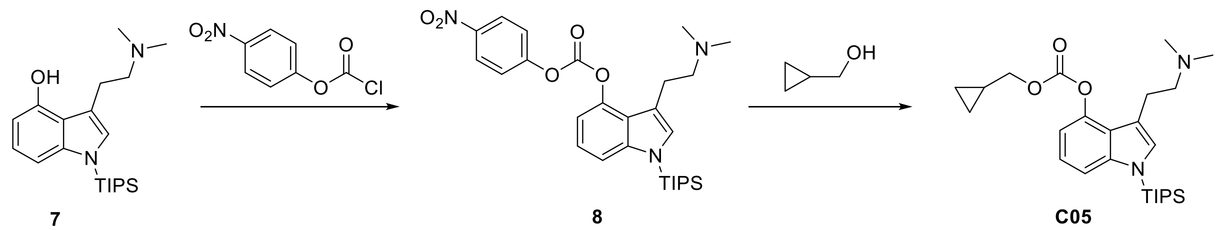

**Cyclopropylmethyl (3-(2-(dimethylamino)ethyl)-1-(triisopropylsilyl)-1*H*-indol-4-yl) carbonate (C05)**

A solution of **7** (250 mg, 693 μmol) and 4-nitrophenyl chloroformate (154 mg, 763 μmol) in DCM (3.47 mL) was cooled down to 0 °C, and to it was added N,N-diisopropylethylamine (242 μL, 1.39 mmol) dropwise. The reaction was warmed up to RT and stirred for 2 h. After 2 h, TLC (MeOH/DCM 1:9) showed almost complete conversion to the desired intermediate, **8**.

To the reaction mixture (crude **8**) was added cyclopropanemethanol (168 μL, 2.08 mmol) and N,N-diisopropylethylamine (362 μL, 2.08 mmol) with vigorous stirring at RT. After 2 h, volatiles were removed in vacuo and the crude mixture was purified on silica gel (MeOH/DCM 1:9) to afford the semi-pure product that contained p-nitrophenol. This material was dissolved in DCM (10 mL) and washed with sat'd aq. $NaHCO_3$ (6 x 20 mL) to remove the residual *p*-nitrophenol. **C05** was obtained as a light-yellow oil (185 mg, 58%).

HRMS-HESI: calculated: 459.3037; observed: 459.3031 m/z $[M+H]^+$

$^1$H NMR (400 MHz, $CDCl_3$) δ 7.34 (d, *J* = 8.3 Hz, 1H), 7.09 (t, *J* = 8.1 Hz, 1H), 7.01 (s, 1H), 6.92 (d, *J* = 7.7 Hz, 1H), 4.12 (d, *J* = 7.4 Hz, 2H), 3.02 – 2.94 (m, 2H), 2.66 (t, *J* = 8.2 Hz, 2H), 2.38 (s, 6H), 1.66 (h, *J* = 7.5 Hz, 3H), 1.31 – 1.22 (m, 1H), 1.13 (d, *J* = 7.5 Hz, 18H), 0.69 – 0.58 (m, 2H), 0.43 – 0.34 (m, 2H).

$^1$H-NMR of **C05**, 400 MHz, $CDCl_3$

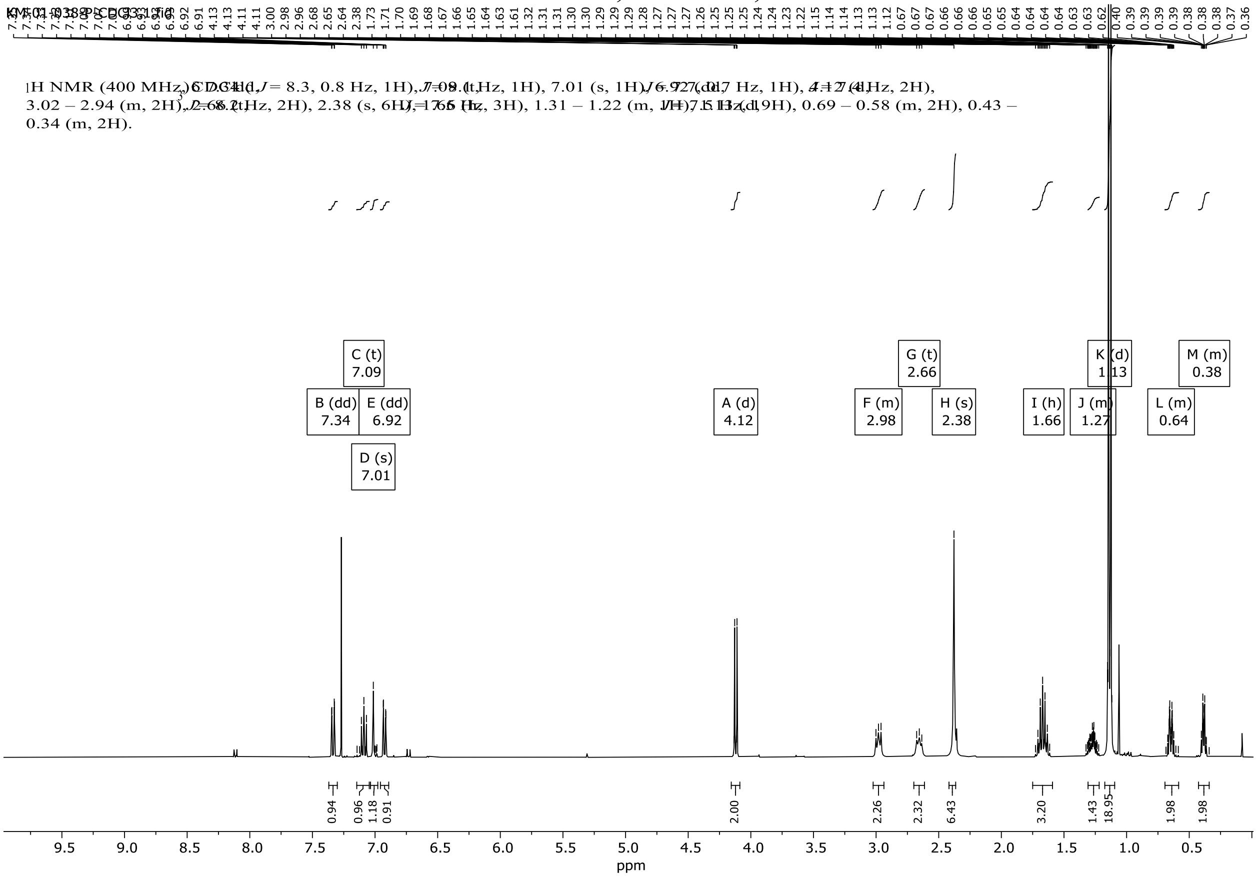

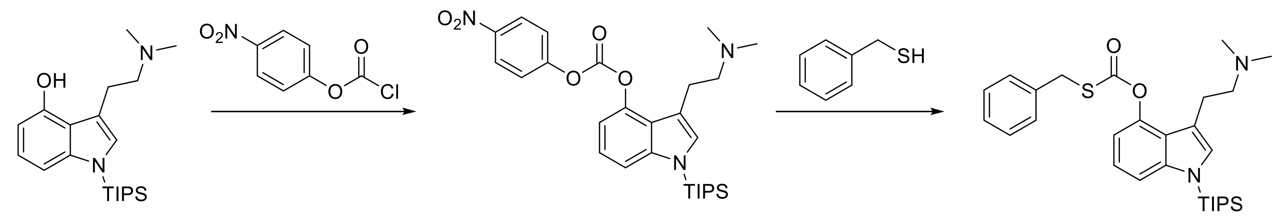


**7** **8** **9**

***S*-Benzyl *O*-(3-(2-(dimethylamino)ethyl)-1*H*-indol-4-yl) carbonothioate (T01)**

A solution of **7** (100 mg, 277 µmol) and 4-nitrophenyl chloroformate (58.7 mg, 291 µmol) in DCM (1.39 mL) was cooled down to 0 °C, and to it was added N,N-diisopropylethylamine (96.6 µL, 555 µmol) dropwise. The reaction was warmed up to RT and stirred for 2 h. After 2 h, TLC (MeOH/DCM 12:88) showed almost complete conversion to the desired product **7**.

To the reaction mixture (**8**) was added a solution of benzyl mercaptan (49.3 µL, 416 µmol) and N,N-diisopropylethylamine (96.5 µL, 554 µmol) in DCM (0.5 mL) with vigorous stirring at RT. After 2 h, volatiles were removed in vacuo and the crude residue (**9**) was used in the next step without further purification.

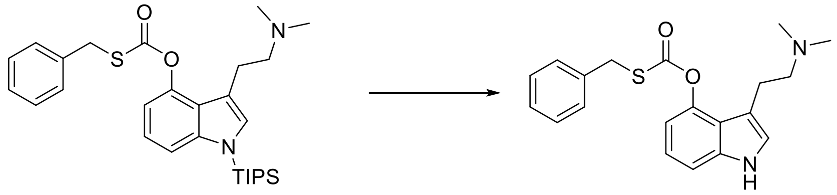


**9** **T01**

To a solution of crude **9** (141 mg, 276 µmol) in dry THF (1.38 mL) was added tetrabutylammonium fluoride solution (1 M in THF, 414 µL, 414 µmol) dropwise at 0 °C. After 30 min, water (2 mL) was added, the aq. layer was separated and extracted with DCM (3 x 15 mL). The combined organic layers were washed with brine, dried over anhydrous $Na_2SO_4$, filtered and concentrated. The crude product was purified by flash chromatography using silica gel (MeOH/DCM 1:9) to afford the pure product **T01** as a yellow waxy solid (33 mg, 34%).

HRMS-HESI: calculated: 355.1475; observed: 355.1467 m/z $[M+H]^+$

$^1H$ NMR (400 MHz, $CDCl_3$) δ 8.32 (s, 1H), 7.41 – 7.26 (m, 5H), 7.21 (d, J = 8.2 Hz, 1H), 7.15 – 7.05 (m, 1H), 6.99 – 6.94 (m, 1H), 6.88 (dd, J = 7.6, 0.8 Hz, 1H), 4.20 (s, 2H), 2.92 (dd, J = 9.0, 6.7 Hz, 2H), 2.66 – 2.58 (m, 2H), 2.30 (s, 6H).

$^1$H-NMR of **T01**, 400 MHz, $CDCl_3$

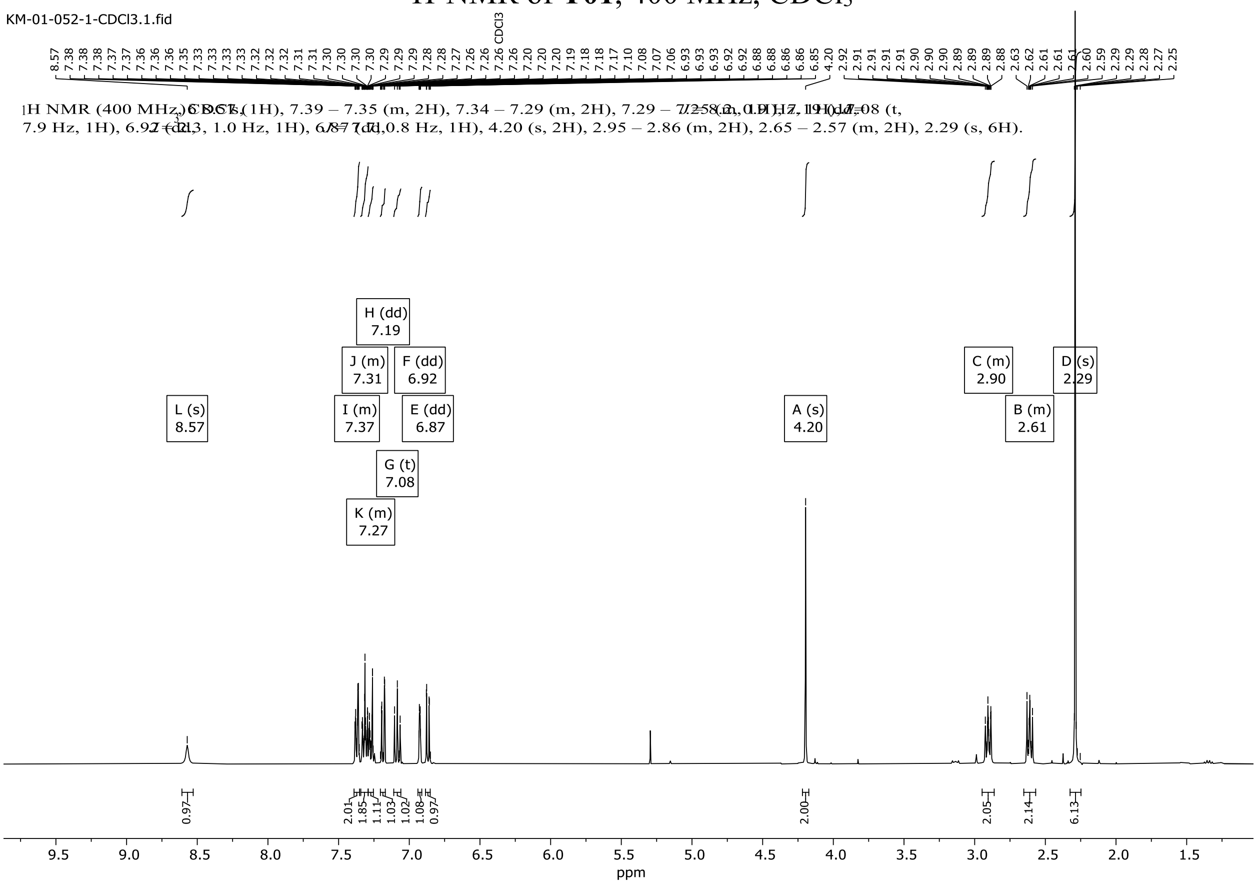

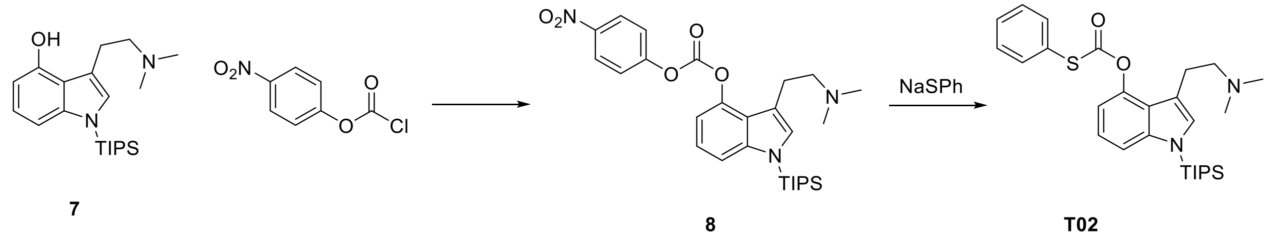


***O*-(3-(2-(Dimethylamino)ethyl)-1-(triisopropylsilyl)-1*H*-indol-4-yl) *S*-phenyl carbonothioate (T02)**

A solution of **7** (150 mg, 416 µmol) and 4-nitrophenyl chloroformate (91.7 mg, 437 µmol) in DCM (2.0 mL) was cooled down to 0 °C. To this was added N,N-diisopropylethylamine (145 µL, 832 µmol) in a dropwise manner. The reaction was warmed up to RT and stirred for 3 h. At this point, TLC (UV) confirmed that the conversion to the reactive intermediate **8** had occurred and the mixture was immediately used in the next step. The DCM was evaporated and replaced with dry THF. Added to this was sodium thiophenolate (67.2 mg, 458 µmol), and the resulting mixture was left to stir overnight at room temperature. The reaction mixture was poured into a separatory funnel containing 10 mL of water and 10 mL of DCM. The aqueous phase was extracted with DCM (3x10 mL); all organic phases were combined, washed with brine, and dried over $MgSO_4$. After filtering, the organic phase was concentrated to leave an oil containing the crude product. Purification was carried out with flash chromatography (9:1 DCM:MeOH) to provide **T02** (113 mg, 55%) as a yellow oil.

MS-ESI: calculated: 497.2653; observed: 497.2647 m/z $[M+H]^+$

$^1H$ NMR (400 MHz, $CDCl_3$) δ 7.64 – 7.58 (m, 2H), 7.45 – 7.39 (m, 3H), 7.34 (dd, J = 8.3, 0.8 Hz, 1H), 7.10 – 7.03 (m, 2H), 6.96 (dd, J = 7.8, 0.7 Hz, 1H), 3.10 (m, 2H), 2.78 (m, 2H), 2.48 (s, 6H), 1.66 (m, J = 7.5 Hz, 3H), 1.12 (d, J = 7.5 Hz, 18H).

$^{1}$H-NMR of **T02**, 400 MHz, $CDCl_3$

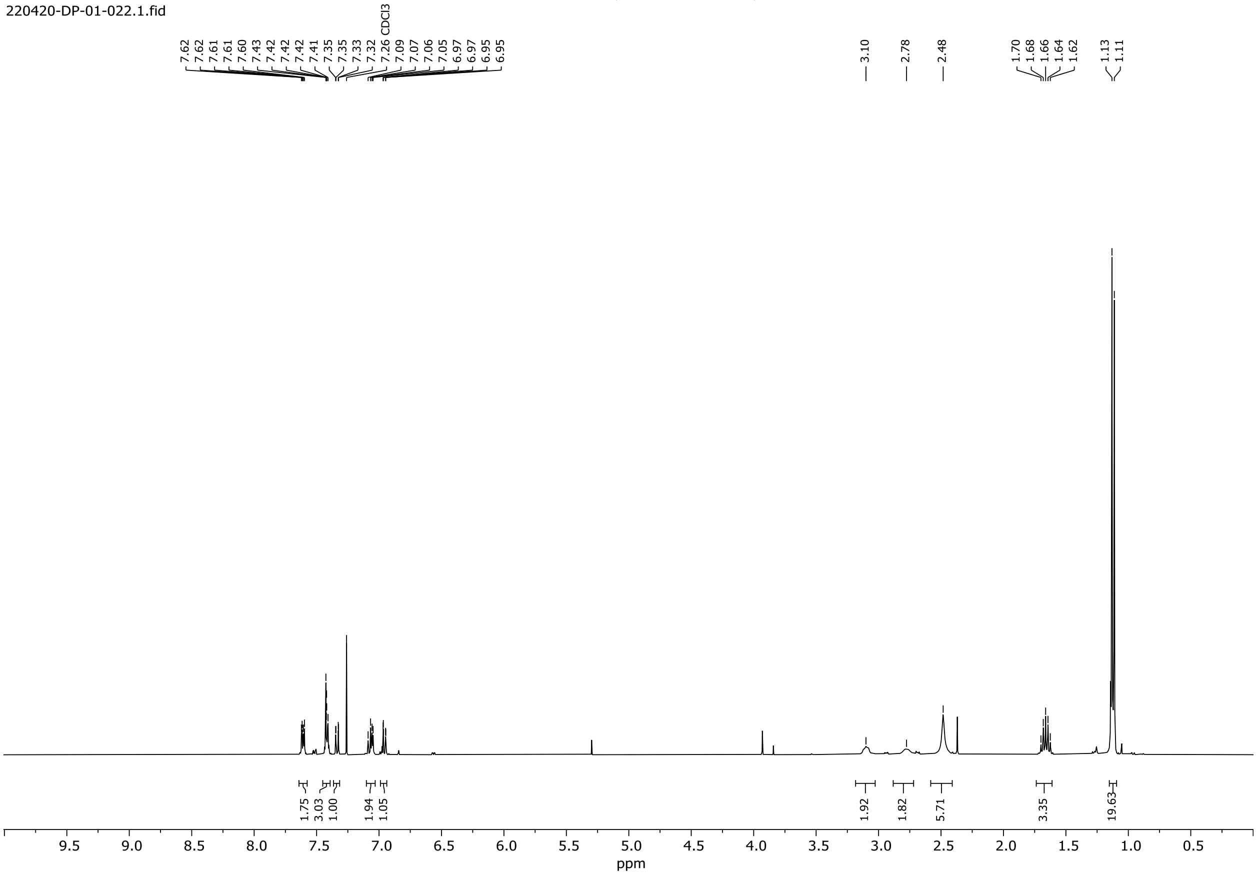

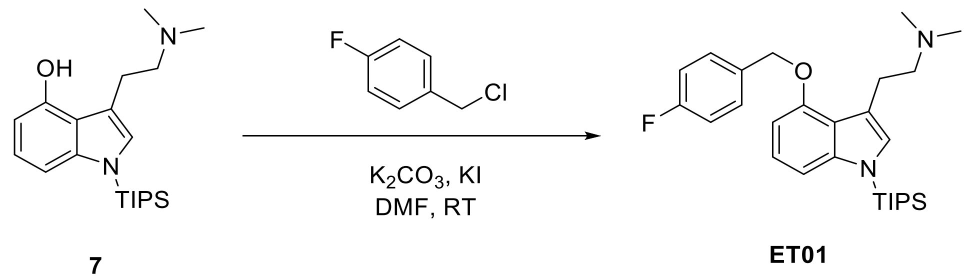

**2-(4-((4-Fluorobenzyl)oxy)-1-(triisopropylsilyl)-1*H*-indol-3-yl)-*N*,*N*-dimethylethanamine (ET01)**

To a solution of *N*-TIPS-Psilocin **7** (22.5 mg, 62.6 µmol) in dry DMF (2 ml) under argon was added potassium carbonate (10.4 mg, 75.1 µmol) and potassium iodide (12.5 mg, 75.1µmol). The mixture was stirred at room temperature for 10 minutes and then 4-fluorobenzyl chloride (10.9 mg, 75.1 mmol) was added. The resulting mixture was stirred at room temperature for 2 hours and the reaction was monitored by TLC. The reaction was quenched by addition of water. The mixture was then extracted with dichloromethane (3x10 mL) and the organic phases were combined, washed with brine, dried over anhydrous $MgSO_4$, filtered and concentrated. The crude residue was purified by flash chromatography on silica gel (12 g, 0% to 10% MeOH/DCM) to afford the product **ET01** as a pale-yellow solid (7.3 mg, 25%).

HRMS-HESI: calculated: 469.3045; observed: 469.3046 m/z $[M+H]^+$

$^1$H NMR (400 MHz, MeOD) δ 7.72 – 7.57 (m, 2H), 7.51 (s, 1H), 7.07 – 6.91 (m, 4H), 6.78 (d, J = 7.5 Hz, 1H), 4.89 (s, 2H), 3.85 – 3.71 (m, 2H), 3.41 – 3.28 (m, 2H), 3.19 (s, 6H), 1.63 (h, J = 7.5 Hz, 3H), 1.10 (d, J = 7.5 Hz, 18H).

$^{1}$H-NMR of **ET01**, 400 MHz, MeOD

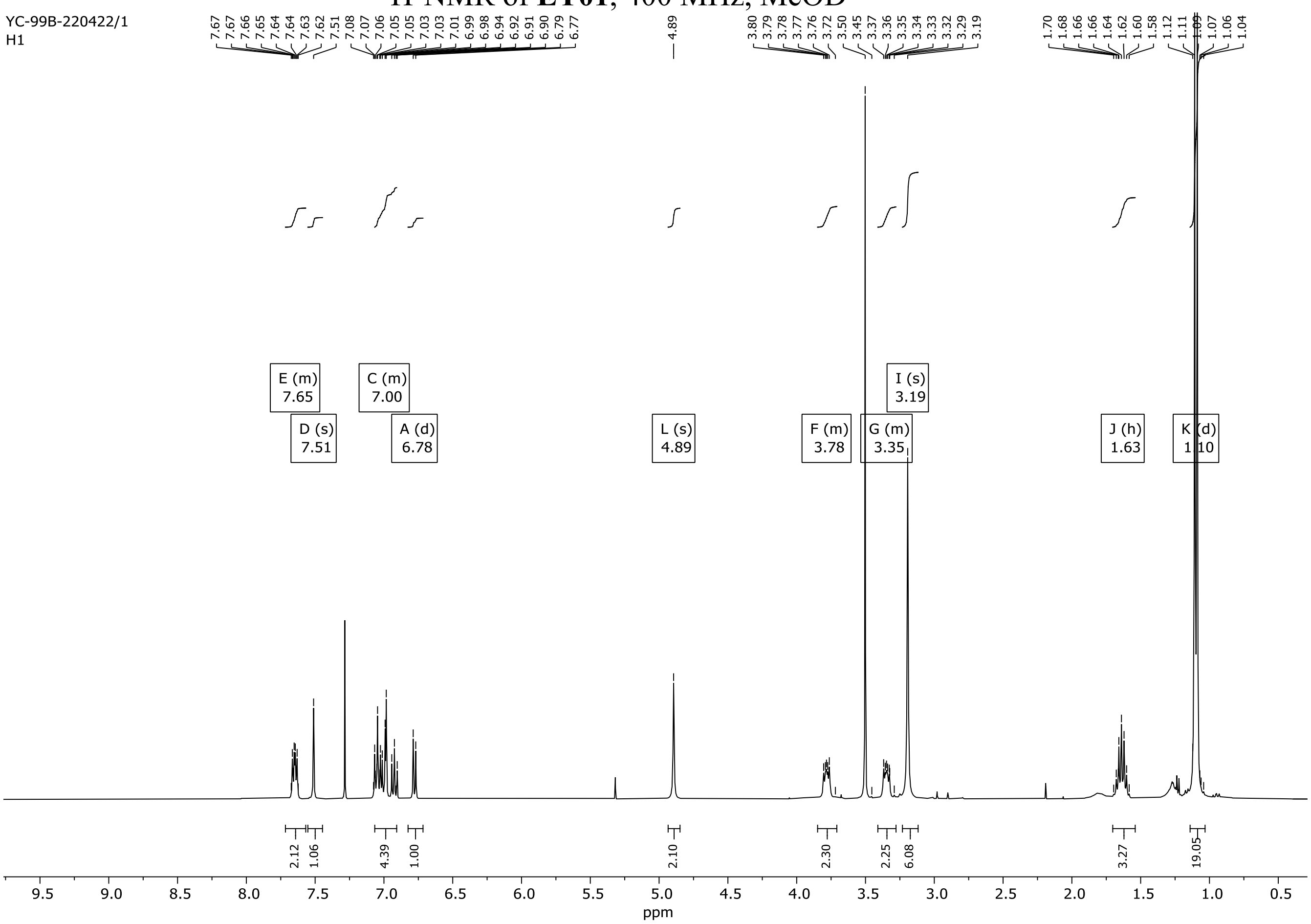

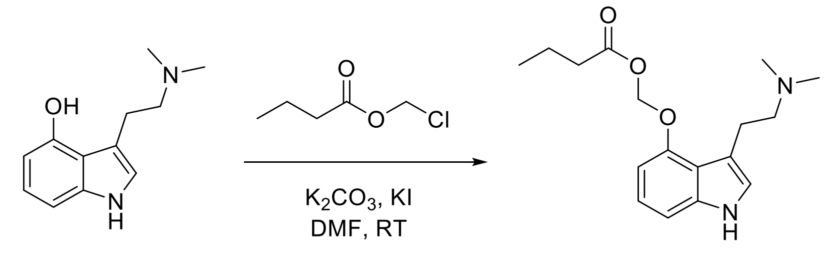


**1** **EE01**

**((3-(2-(Dimethylamino)ethyl)-1*H*-indol-4-yl)oxy)methyl butyrate (EE01)**

To a solution of psilocin **1** (51 mg, 250 µmol) in dry DMF (2 ml) under argon was added potassium carbonate (34.5 mg, 250 µmol) and potassium iodide (41.5 mg, 250 µmol). After stirring at room temperature for 10 minutes, chloromethyl butyrate (34.1 mg, 250 µmol) was added, and the resulting mixture was stirred for 3 days at room temperature. The reaction was quenched by addition of water. The mixture was then extracted with DCM (3 x 5 mL), the organic layers were combined and washed with brine, dried over anhydrous $MgSO_4$, filtered and concentrated. The crude residue was purified by flash chromatography on silica gel (12 g, 7.5% to 10% MeOH/DCM) to afford the product **EE01** as a white solid (41.3 mg, 54.3%).

HRMS-HESI: calculated: 305.1865; observed: 305.1860 m/z $[M+H]^+$

$^1$H NMR (400 MHz, MeOD) δ 7.46 (dd, J = 8.3, 0.8 Hz, 1H), 7.36 (d, J = 1.0 Hz, 1H), 7.29 – 7.18 (m, 1H), 6.87 (td, J = 8.1, 0.9 Hz, 1H), 5.60 (s, 2H), 3.56 – 3.44 (m, 2H), 3.25 – 3.10 (m, 2H), 2.93 (s, 6H), 2.73 (t, J = 7.3 Hz, 2H), 1.84 (h, J = 7.4 Hz, 2H), 1.11 (t, J = 7.4 Hz, 3H).

$^1$H-NMR of **EE01**, 400 MHz, MeOD

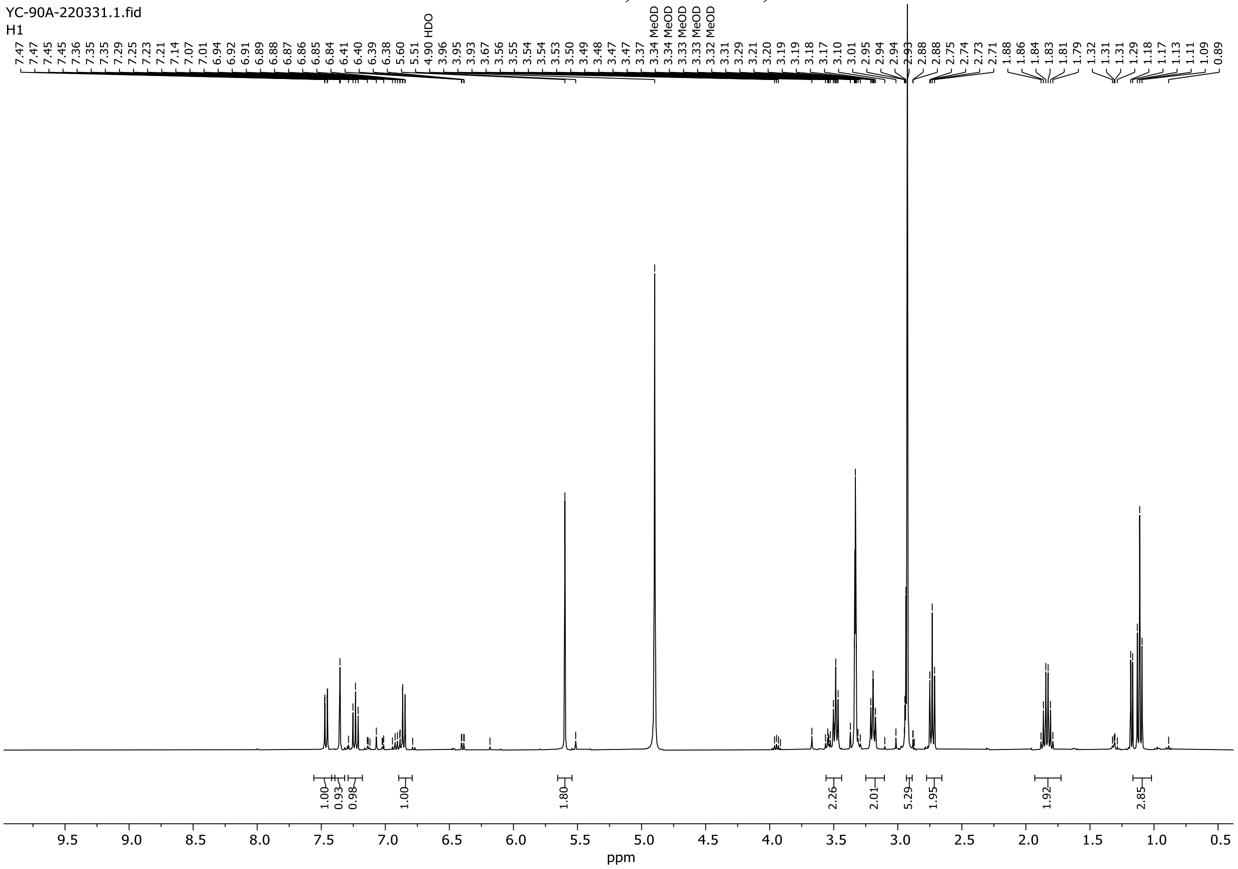

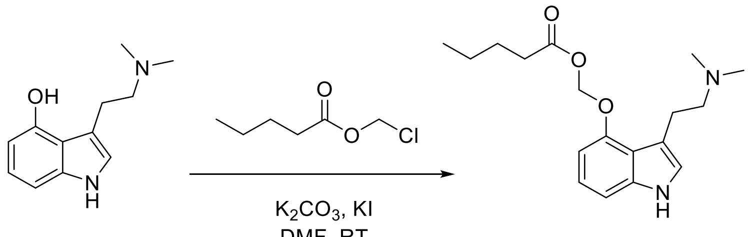


**1** **EE02**

**((3-(2-(Dimethylamino)ethyl)-1*H*-indol-4-yl)oxy)methyl pentanoate (EE02)**

To a solution of psilocin **1** (102 mg, 500 µmol) in dry DMF (2 ml) under argon was added potassium carbonate (69 mg, 500 µmol) and potassium iodide (83 mg, 500 µmol). After stirring at room temperature for 10 minutes, chloromethyl pentanoate (75.2 mg, 500 µmol) was added, and the resulting mixture was stirred for 3 days at room temperature. The reaction was quenched by addition of water. The mixture was then extracted with DCM (3 x 5 mL), the organic layers were combined, washed with brine, dried over anhydrous $MgSO_4$, filtered and concentrated. The crude residue was purified by FC on silica gel (12 g, 7.5% to 10% MeOH/DCM) to afford the pure product **EE02** as a white solid (40.7 mg, 25.6%).

HRMS-HESI: calculated: 319.2022; observed: 319.2014 m/z $[M+H]^+$

$^1$H NMR (400 MHz, MeOD) δ 7.46 (dd, J = 8.3, 0.8 Hz, 1H), 7.37 (d, J = 1.0 Hz, 1H), 7.23 (t, J = 8.0 Hz, 1H), 6.85 (dd, J = 7.7, 0.8 Hz, 1H), 5.59 (s, 2H), 3.52 – 3.45 (m, 2H), 3.24 – 3.14 (m, 2H), 2.93 (s, 6H), 2.76 (t, J = 7.4 Hz, 2H), 1.84 – 1.74 (m, 2H), 1.58 – 1.47 (m, 2H), 1.03 (t, J = 7.4 Hz, 3H).

$^1$H-NMR of **EE02**, 400 MHz, MeOD

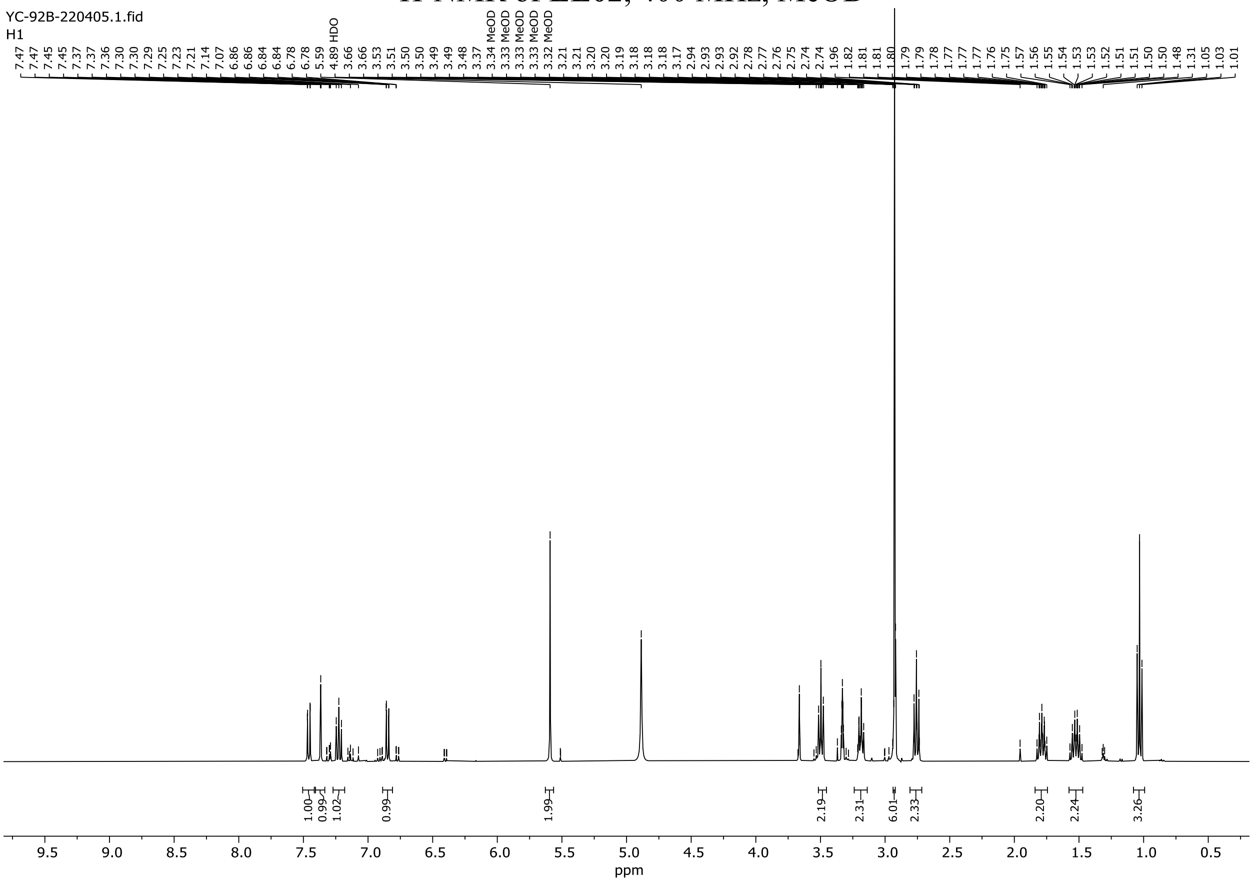

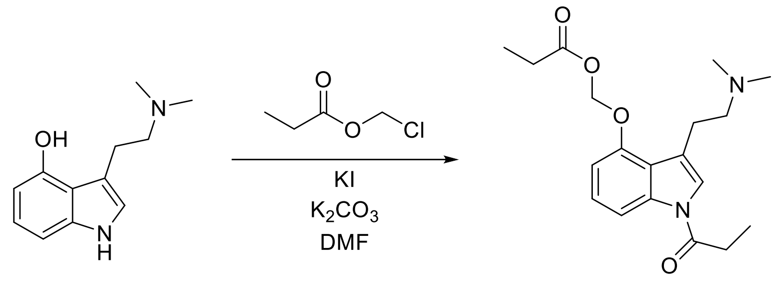

1 EE03

**((3-(2-(Dimethylamino)ethyl)-1-propionyl-1*H*-indol-4-yl)oxy)methyl propionate (EE03)**

To a solution of **1** (101 mg, 494 µmol) in dry DMF (1.00 mL) under argon was added potassium carbonate (68.3 mg, 494 µmol) and potassium iodide (82.1 mg, 494 µmol). After stirring at room temperature for 10 minutes, chloromethyl propionate (55.1 µL, 494 µmol) in dry DMF (500 µL) was added, and the resulting mixture was stirred overnight at 70 °C. The mixture was diluted with water (10 mL) and washed with brine (10 mL). The brine was extracted with DCM (3 x 10 mL). An emulsion formed during the extraction that was subsequently filtered off. The organic layer was dried over anhydrous $MgSO_4$, filtered and concentrated. The crude residue was purified by flash column chromatography on silica gel (12 g, 0 to 10% MeOH/DCM) to yield a mixture of products. This mixture was further purified by a second flash column chromatography on silica gel (4 g, 2.5 to 5% MeOH/DCM) to yield **EE03** (6 mg, 4%) as a colorless oil.

HRMS-HESI: calculated: 347.1965; observed: 347.1963 m/z $[M+H]^+$

$^1$H NMR (400 MHz, MeOD) δ 7.42 (dd, $J$ = 8.3, 0.8 Hz, 1H), 7.24 – 7.17 (m, 2H), 6.81 (dd, $J$ = 7.8, 0.8 Hz, 1H), 6.14 (s, 2H), 2.93 – 2.87 (m, 2H), 2.76 (q, $J$ = 7.5 Hz, 2H), 2.69 – 2.62 (m, 2H), 2.37 – 2.33 (m; 8H), 1.29 (t, $J$ = 7.5 Hz, 3H), 1.09 (t, $J$ = 7.5 Hz, 3H).

[1]H-NMR of **EE03**, 400 MHz, MeOD

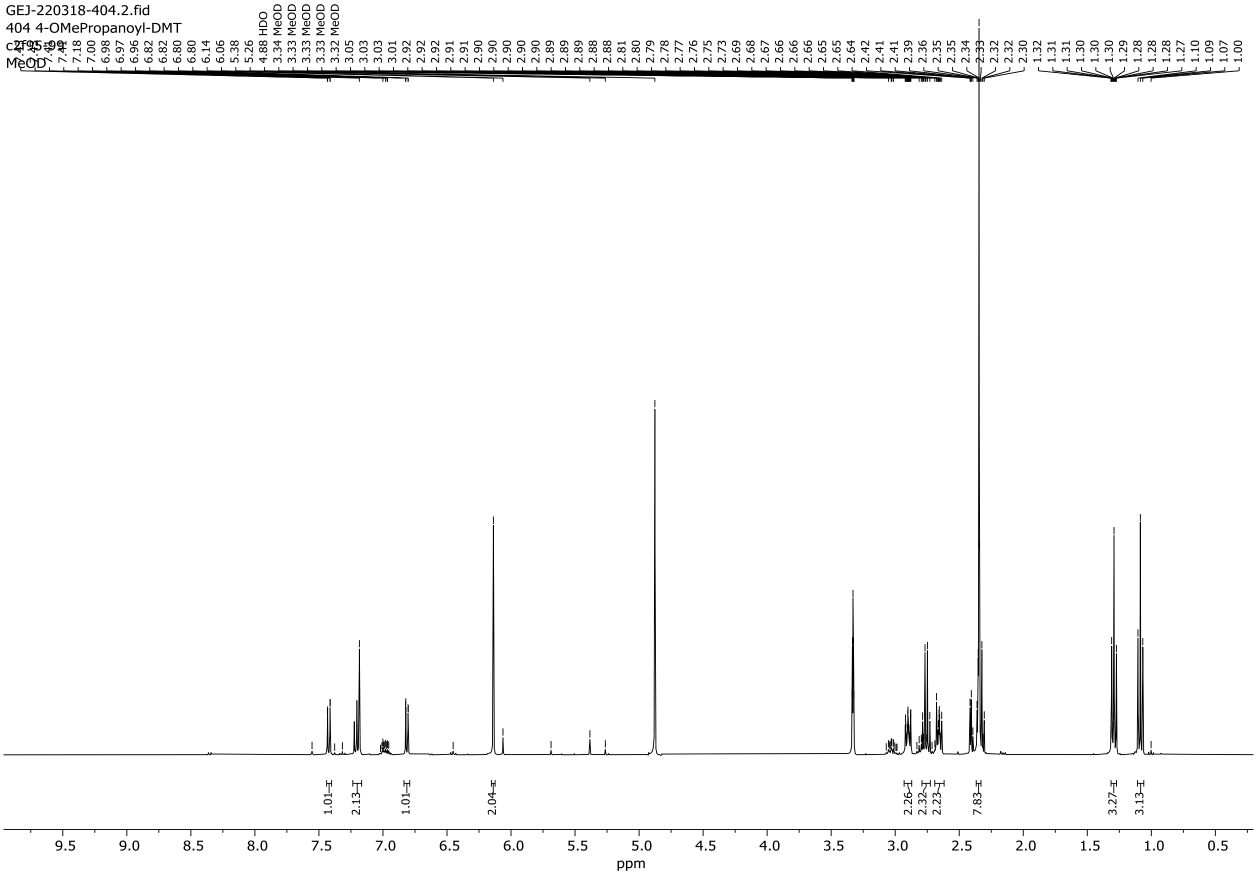

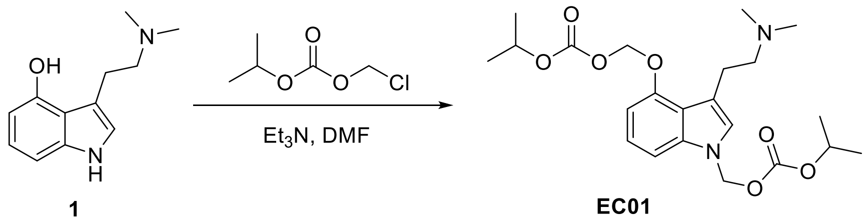

## EC01

A dry round-bottom flask was charged with psilocin **1** (100 mg, 0.49 mmol) and dry DMF (2 mL) under argon. To the flask was added $Et_3N$ (69 µL, 0.49 mmol) followed by chloromethyl isopropyl carbonate (71 mg, 0.47 mmol), and the resulting mixture was stirred at 70 °C for 16 h. Upon completion (TLC), the reaction mixture was diluted with water (20 mL) and extracted with DCM (3 x 15 mL). The combined organic layers were washed with brine, dried over anhydrous $MgSO_4$, filtered and concentrated. The crude residue was purified by flash chromatography using silica gel (MeOH/DCM 0:100 to 20:80, gradient) to afford the pure product **EC01** as a colorless oil (33 mg, 15%).

HRMS-HESI: calculated: 437.2282; observed: 437.2275 m/z $[M+H]^+$

$^1$H NMR (400 MHz, MeOD) δ 7.46 (dd, J = 8.3, 0.8 Hz, 1H), 7.34 (s, 1H), 7.24 (t, J = 8.1 Hz, 1H), 7.06 (dd, J = 7.8, 0.7 Hz, 1H), 5.58 (s, 2H), 5.42 (s, 2H), 5.09 – 4.93 (m, 2H), 3.66 – 3.59 (m, 2H), 3.35 – 3.30 (m, 2H), 3.28 (s, 6H), 1.40 (d, J = 6.3 Hz, 6H), 1.36 (d, J = 6.3 Hz, 6H).

$^1$H-NMR of **EC01**, 400 MHz, MeOD

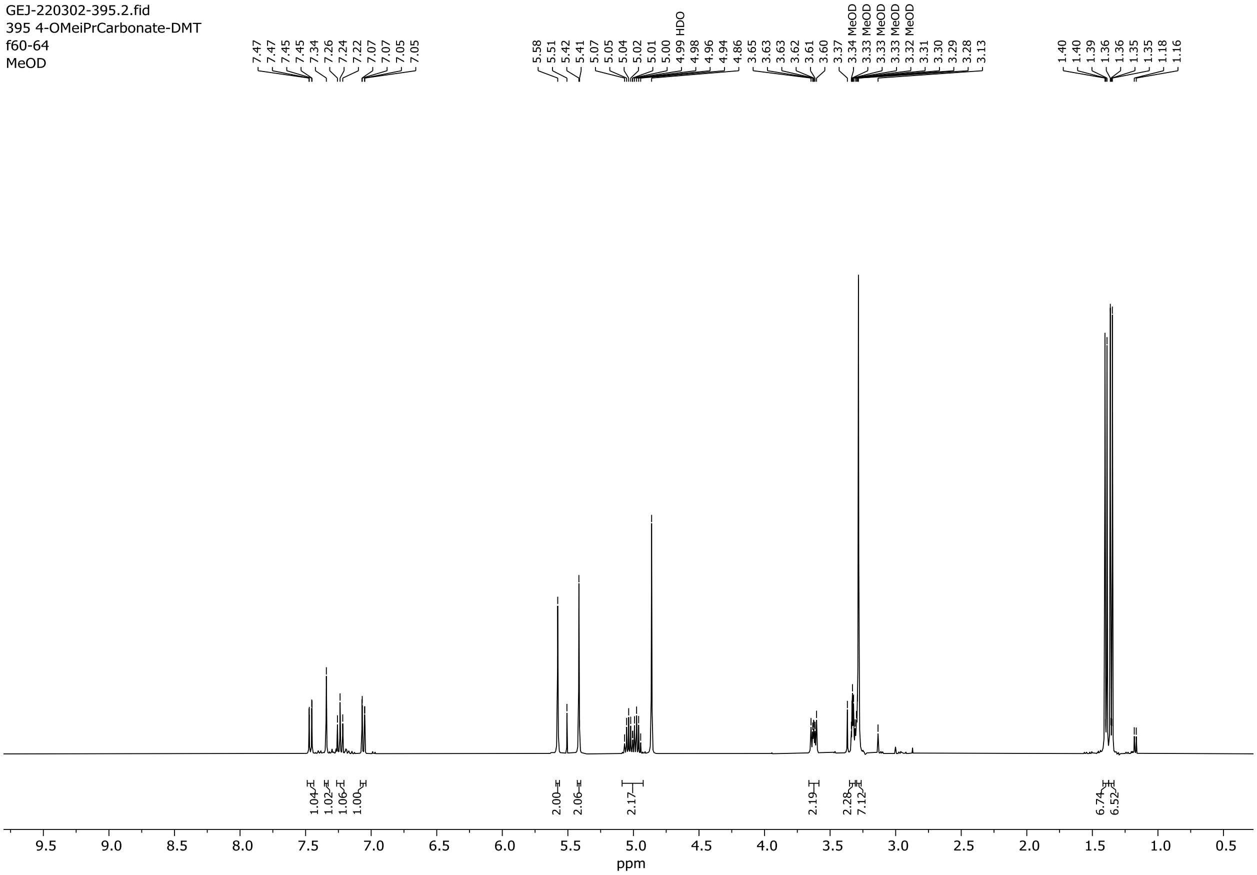

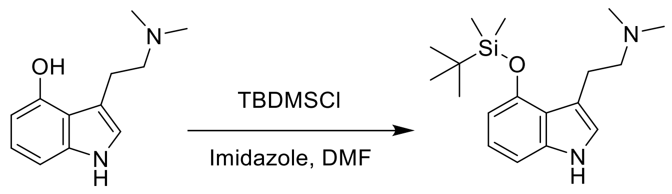


1 SE01

**2-(4-((*tert*-Butyldimethylsilyl)oxy)-1*H*-indol-3-yl)-*N*,*N*-dimethylethanamine (SE01)**

Compound **1** (100 mg, 0.49 mmol) and imidazole (100 mg, 1.5 mmol) were dissolved in anhydrous DMF (1.5 mL) under argon atmosphere, to which *tert*-butyl(chloro)dimethylsilane (89 mg, 0.59 mmol) was added. The solution was allowed to stir for 18 hours until the reaction was complete as determined by TLC (20% MeOH/DCM). The solvent was removed under reduced pressure at 80 °C to dryness, then the residue was dissolved in ethyl acetate (15 mL), washed with water (3 x 10 mL), dried over anhydrous $MgSO_4$, filtered and concentrated under reduced pressure. The crude residue was purified by FC on silica gel (12 g, 0 to 10% MeOH/DCM). The isolated material was dissolved in dichloromethane (15 mL), washed with brine (3 x 10 mL), dried over anhydrous $MgSO_4$, filtered and concentrated to yield **SE01** (80 mg, 80%) as a white solid.

HRMS-HESI: calculated: 319.2200; observed: 319.2195 m/z $[M+H]^+$

$^1$H NMR (400 MHz, MeOD) δ 7.00 – 6.88 (m, 3H), 6.45 (dd, J = 7.3, 1.1 Hz, 1H), 3.22 – 3.14 (m, 2H), 2.92 (dd, J = 8.5, 6.8 Hz, 2H), 2.45 (s, 6H), 1.06 (s, 9H), 0.35 (s, 6H).

$^1$H-NMR of **SE01**, 400 MHz, MeOD

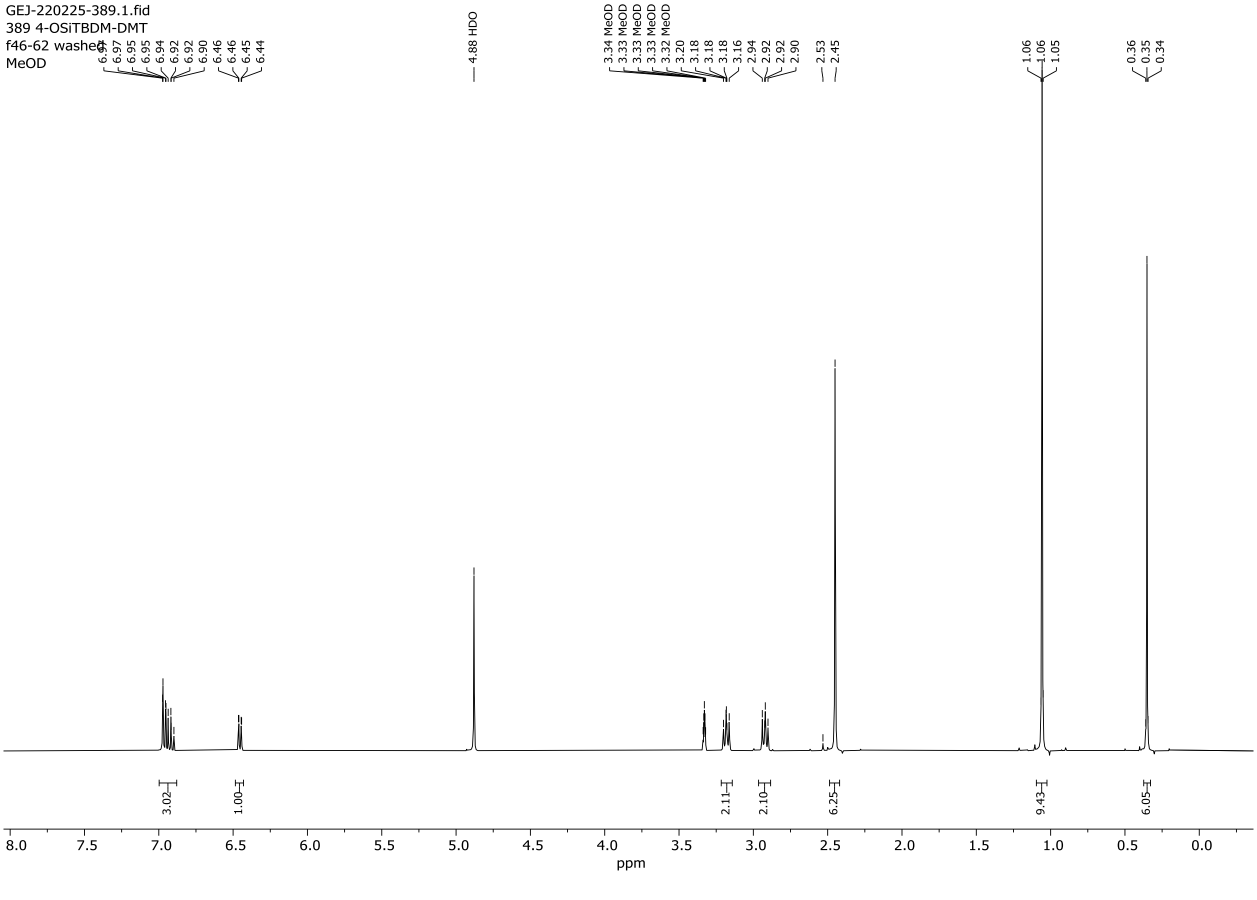

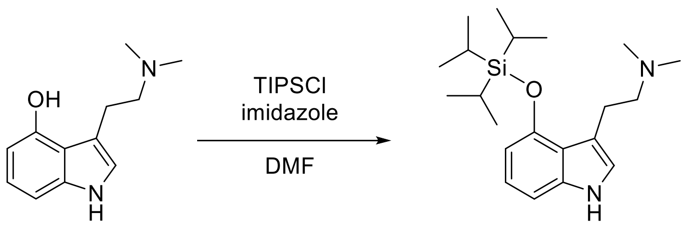


**1** **SE02**

**_N_,_N_-Dimethyl-2-(4-((triisopropylsilyl)oxy)-1_H_-indol-3-yl)ethanamine (SE02)**

A dry round-bottom flask was charged with psilocin **1** (100 mg, 0.49 mmol) and imidazole (100 mg, 1.47 mmol) under argon. Then, anhydrous DMF (1.50 mL) followed by triisopropylsilyl chloride (TIPSCl) (126 µL, 0.59 mmol) were added to the flask, and the resulting solution was stirred at room temperature for 4 h (or until completion). The volatiles were removed under reduced pressure, and the obtained residue was re-dissolved in EtOAc/MeOH 9:1 (10 mL). The resulting solution was washed with sat'd aq. $NaHCO_3$, followed by brine. The organic layer was dried over anhydrous $MgSO_4$, filtered and concentrated. The crude residue was purified by flash chromatography using silica gel (MeOH/DCM, 0:100 to 10:90, gradient) to afford the pure product **SE02** as an off-white solid (22 mg, 22%).

HRMS-HESI: calculated: 361.2670; observed: 361.2663 m/z $[M+H]^+$

$^1$H NMR (400 MHz, MeOD) δ 6.96 – 6.86 (m, 3H), 6.44 – 6.40 (m, 1H), 3.20 – 3.14 (m, 2H), 2.80 – 2.73 (m, 2H), 2.33 (s, 6H), 1.50 – 1.39 (m, 3H), 1.19 (d, $J$ = 7.5 Hz, 18H).

$^1$H-NMR of **SE02**, 400 MHz, MeOD

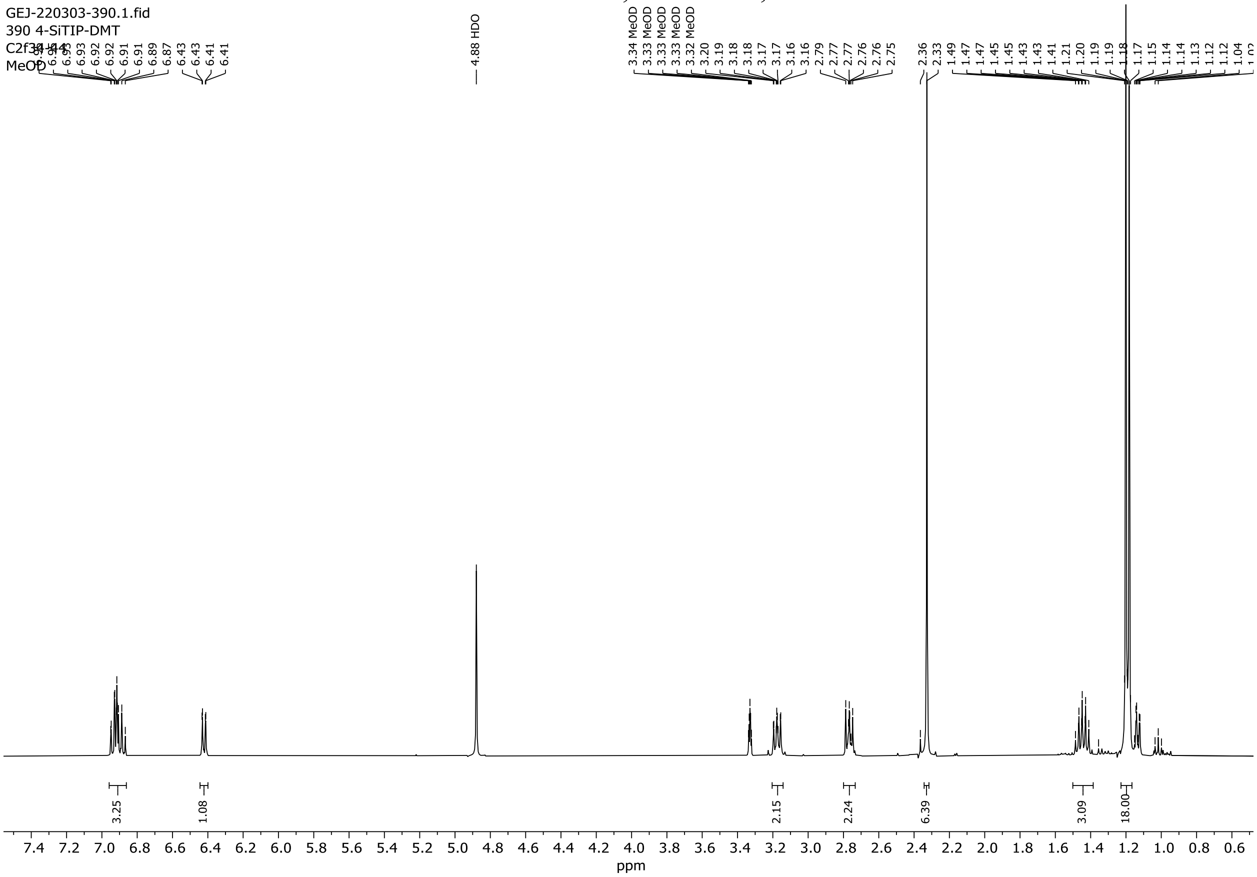

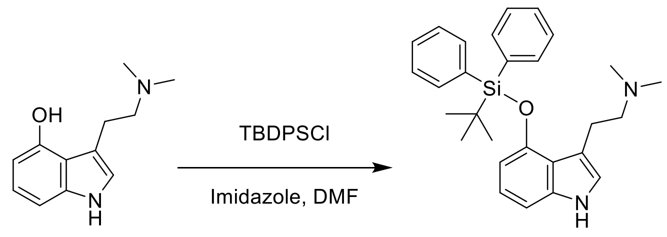


**1** **SE03**

**2-(4-((*tert*-Butyldiphenylsilyl)oxy)-1*H*-indol-3-yl)-*N*,*N*-dimethylethanamine (SE03)**

Compound **1** (100 mg, 0.49 mmol) and imidazole (100 mg, 1.5 mmol) were dissolved in anhydrous DMF (1.5 mL) under argon atmosphere, to which *tert*-butyl(chloro)diphenylsilane (153 µL, 0.59 mmol) was added. The solution was allowed to stir for 27 hours until the reaction was complete as determined by TLC (20% MeOH/DCM). The solvent was removed under reduced pressure at 80 °C to dryness. The crude product was directly purified by flash column chromatography on silica gel (12 g, 0 to 10% MeOH/DCM). The resulting material was dissolved in ethyl acetate (15 mL) and washed with water (3 x 10 mL) to remove residual imidazole. The organic phase was dried over anhydrous $MgSO_4$, filtered and concentrated to yield **SE03** (72 mg, 33%) as a white solid.

HRMS-HESI: calculated: 443.2513; observed: 443.2509 m/z $[M+H]^+$

$^1$H NMR (400 MHz, MeOD) δ 7.85 – 7.78 (m, 4H), 7.49 – 7.34 (m, 6H), 6.99 (d, J = 1.3 Hz, 1H), 6.86 (dt, J = 8.1, 0.6 Hz, 1H), 6.57 – 6.48 (m, 1H), 6.02 – 5.96 (m, 1H), 3.38 (d, J = 7.6 Hz, 2H), 2.91 – 2.84 (m, 2H), 2.37 – 2.30 (m, 6H), 1.15 (s, 9H).

$^1$H-NMR of **SE03**, 400 MHz, MeOD

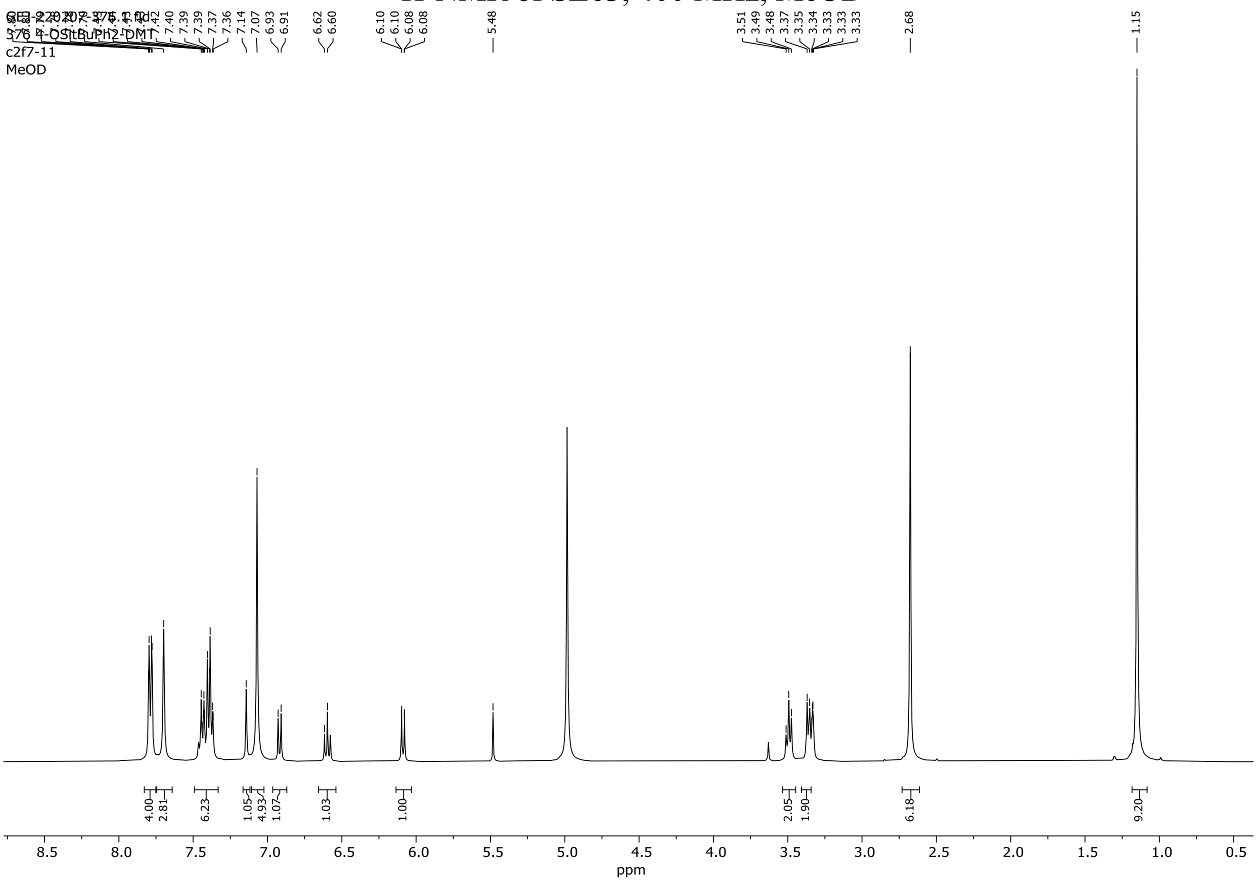

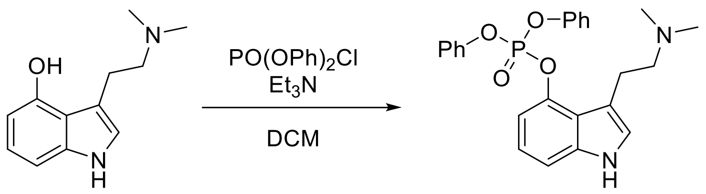


**1** **P02**

**3-(2-(Dimethylamino)ethyl)-1*H*-indol-4-yl diphenyl phosphate (P02)**

A dry round-bottom flask was charged with psilocin **1** (137 mg, 0.67 mmol) and dry DCM (1 mL) under argon. To this suspension, was added $Et_3N$ (188 μL, 1.34 mmol) and the resulting mixture was cooled down to 0 °C. To the reaction mixture was added a solution of diphenylphosphoryl chloride (278 μL, 1.34 mmol) in DCM (0.5 mL), and the resulting mixture was stirred at room temperature for 16 h. Upon completion (TLC), the mixture was diluted with DCM (10 mL) and washed with sat'd aq. $NaHCO_3$ (2 x 25 mL) and brine. The organic layer was dried over anhydrous $MgSO_4$, filtered and concentrated. The crude material was purified by flash chromatography using silica gel (MeOH/DCM, 0:100 to 10:90, gradient) to afford the pure product **P02** as a light-yellow oil (109 mg, 37%).

HRMS-HESI: calculated: 437.1625; observed: 437.1620 m/z $[M+H]^+$

$^1$H NMR (400 MHz, MeOD) δ 7.37 – 7.10 (m, 14H), 3.11 – 2.97 (m, 2H), 2.81 – 2.69 (m, 2H), 2.35 (s, 6H).

$^1$H-NMR of **P02**, 400 MHz, MeOD

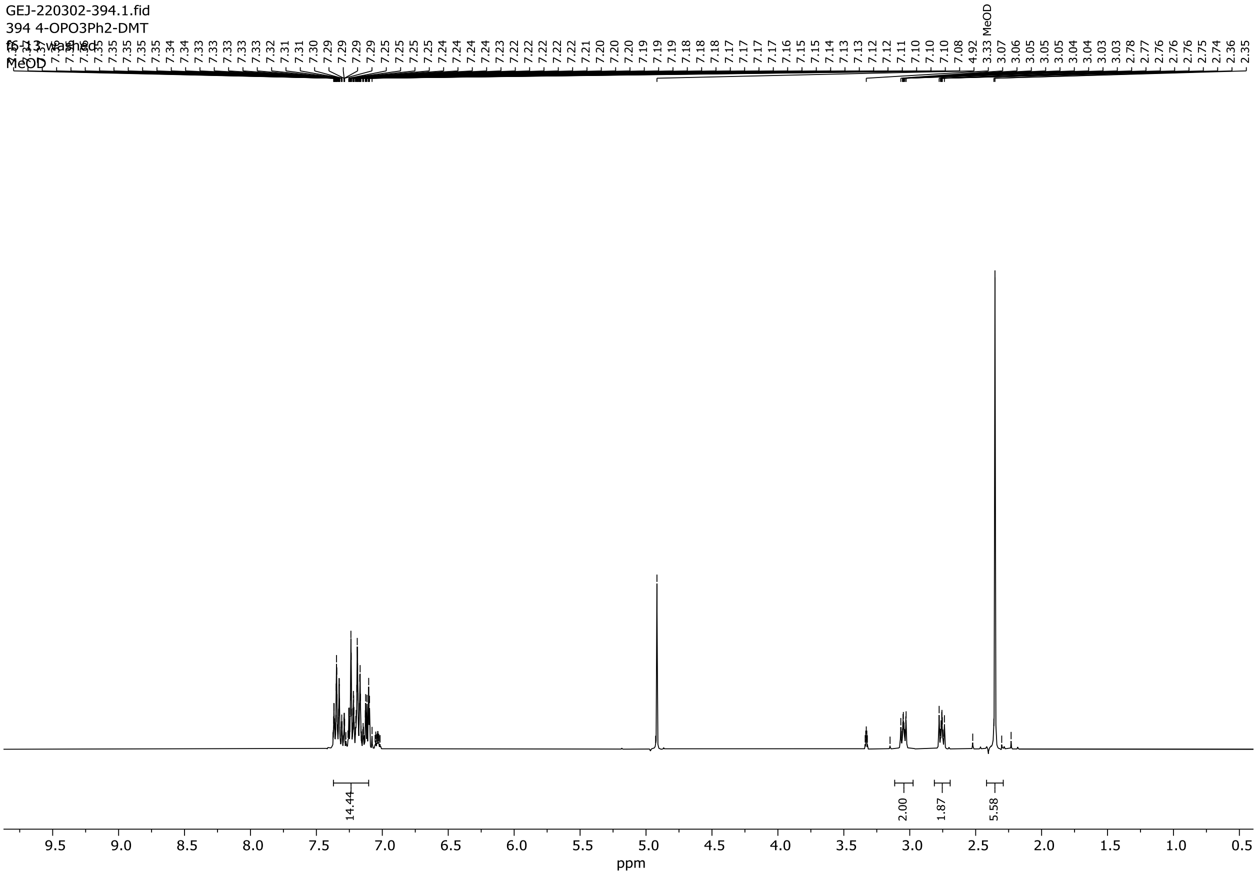

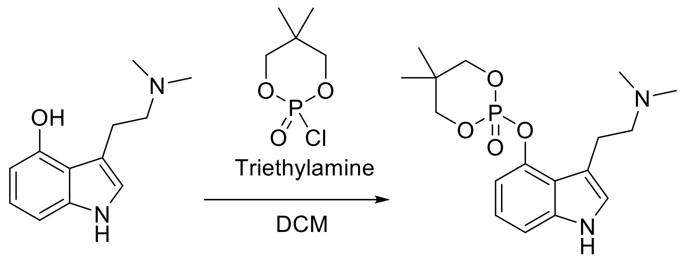

1 P03

**2-((3-(2-(Dimethylamino)ethyl)-1*H*-indol-4-yl)oxy)-5,5-dimethyl-1,3,2-dioxaphosphinane 2-oxide (P03)**

Compound **1** (101 mg, 0.49 mmol) was suspended in anhydrous DCM (1 mL) under argon atmosphere. Triethylamine (0.14 mL, 0.98 mmol) was added, followed by 2-chloro-5,5-dimethyl-1,3,2-dioxaphosphorinane-2-oxide (182 mg, 0.98 mmol) in anhydrous DCM (0.5 mL) was added. The resulting mixture was stirred at room temperature for 18 hours and monitored by TLC (20% MeOH/DCM). The mixture was diluted with DCM (10 mL) and washed with saturated aq. $NaHCO_3$ (10 mL) and brine (2 x 10 mL). The organic phase was dried over anhydrous $MgSO_4$, filtered and concentrated to yield an amber oil. The crude residue was purified by flash column chromatography on silica gel (12 g, 0 to 40% MeOH/DCM) to yield **P03** (55 mg, 29%) as a white solid.

HRMS-HESI: calculated: 353.1625; observed: 353.1624 m/z $[M+H]^+$

$^1H$ NMR (400 MHz, $CDCl_3$) δ 9.41 (s, 1H), 7.08 – 7.02 (m, 1H), 7.00 – 6.94 (m, 2H), 6.75 (d, J = 2.4 Hz, 1H), 4.31 (ddd, J = 11.5, 2.6, 1.4 Hz, 2H), 4.10 – 3.93 (m, 2H), 3.12 – 2.98 (m, 2H), 2.73 – 2.58 (m, 2H), 2.35 (s, 6H), 1.34 (s, 3H), 0.87 (s, 3H).

$^{1}$H-NMR of **P03**, 400 MHz, $CDCl_3$

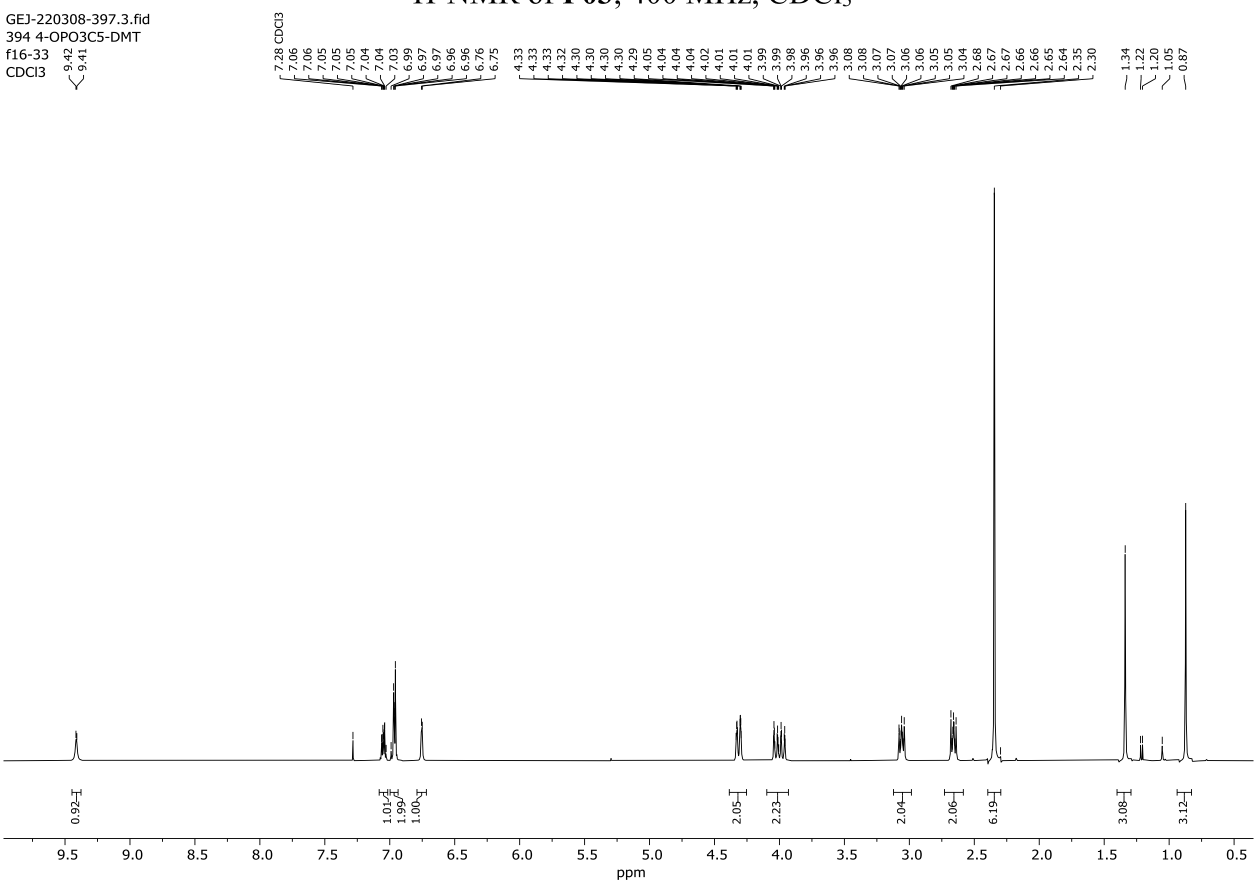

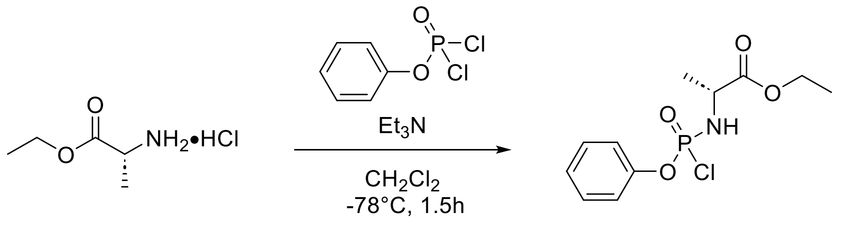


**10**

**(2*R*)-Ethyl 2-((((3-(2-(dimethylamino)ethyl)-1-(triisopropylsilyl)-1*H*-indol-4-yl)oxy)(phenoxy)phosphoryl)amino)propanoate (PA01)**

Phenyldichlorphosphate (354 µL, 2.25 mmol) and *L*-Alanine ethyl ester hydrochloride (349 mg, 2.25 mmol) were dissolved in DCM (5.00 mL). The mixture was cooled down to -78˚C followed by dropwise addition of triethylamine (631 µL, 4.50 mmol). The reaction was stirred for 30 minutes at -78 °C, then slowly warmed up to room temperature over an hour. The crude mixture was concentrated *in vacuo*, redissolved in $Et_2O$, and filtered. The filtrate was purified by FC on silica gel (EA/hex 3:7) to afford compound **10** (520 mg, 79%) as a colorless oil.

$^1$H NMR (400 MHz, $CDCl_3$) δ 7.41 – 7.34 (m, 2H), 7.29 – 7.22 (m, 3H), 4.37 – 4.09 (m, 4H), 1.54 – 1.49 (m, 2H), 1.35 – 1.27 (m, 3H).

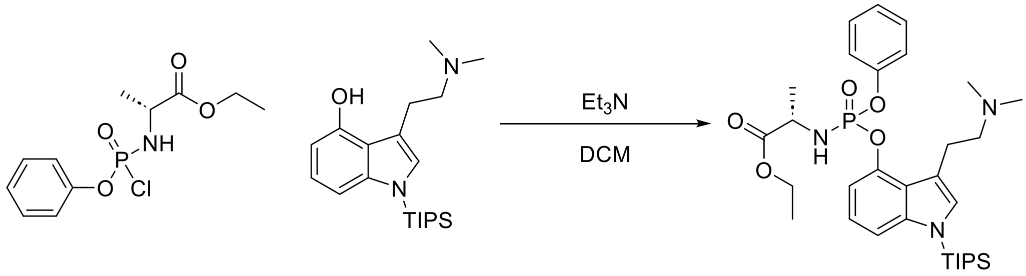


**10** **7** **PA01**

Compound **7** (40.0 mg, 111 µmol) was added to anhydrous DCM (1.00 mL) followed by the addition of triethylamine (31.1 µL, 222 µmol), the mixture was stirred for 5 minutes before the addition of compound **10** (64.7 mg, 222 µmol). The mixture was then stirred at room temperature overnight. The reaction mixture was washed with water and saturated aq. $NaHCO_3$, then dried over anhydrous $MgSO_4$, filtered and concentrated. The crude residue was purified on silica gel (0-10% MeOH/DCM) to afford **PA01** (17 mg, 25%) as a colorless oil.

HRMS-HESI: calculated: 616.3330; observed: 616.3330 m/z [M+H]+

$^1$H NMR (400 MHz, $CDCl_3$) δ 7.31 – 7.21 (m, 6H), 7.14 – 7.03 (m, 3H), 5.36 (s, 1H), 4.23 – 4.10 (m, 1H), 4.05 (q, J = 7.1 Hz, 2H), 3.58 (s, 2H), 3.29 (dt, J = 15.5, 8.7 Hz, 1H), 3.04 (d, J = 14.0 Hz, 1H), 2.82 (s, 6H), 1.65 (h, J = 7.5 Hz, 3H), 1.34 (dd, J = 7.2, 1.2 Hz, 3H), 1.17 (t, J = 7.1 Hz, 3H), 1.11 (d, J = 7.5 Hz, 18H).

#### $^1$H-NMR of **10**, 400 MHz, $CDCl_3$

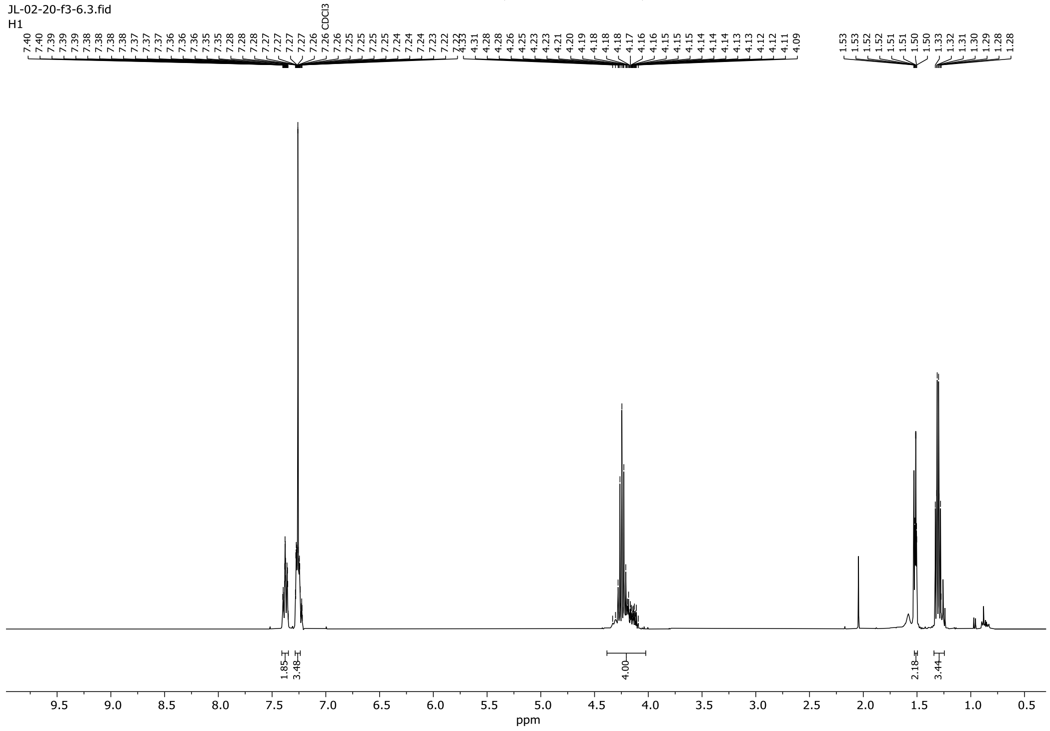

#### $^1$H-NMR of **PA01**, 400 MHz, $CDCl_3$

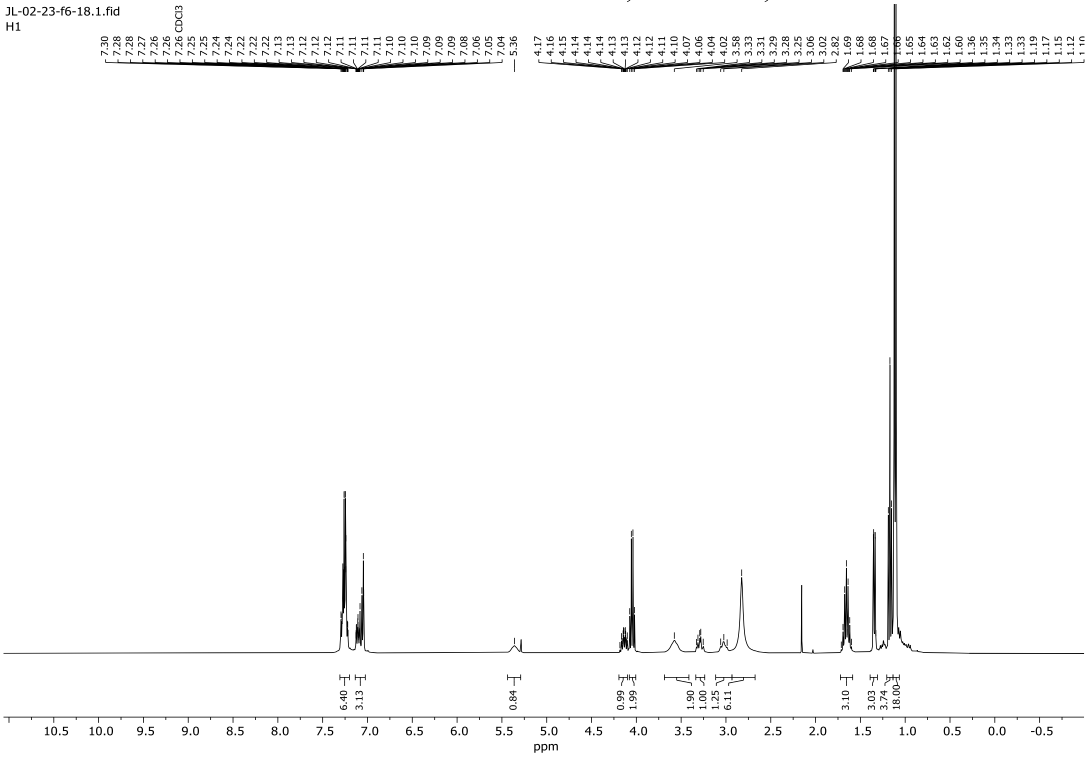
